# Plasmalogen biosynthesis genes decline with age across mammalian tissues and glia, and plasmalogen deficiency fragments brain mitochondria

**DOI:** 10.64898/2026.11.25.769321

**Authors:** Niwase Shamim, Masanori Honsho, Shiro Mawatari, Takehiko Fujino

## Abstract

Plasmalogens are peroxisome-derived ether phospholipids enriched in brain, myelin and immune cells, yet whether ageing switches off their synthesis, and where, has been unknown. Across 105 human, macaque and mouse tissue age models, transcripts of *GNPAT* and *AGPS*, the peroxisomal enzymes that commit precursors to the ether-lipid route, and of the fatty-acyl-CoA reductase *FAR2* fell with age independently of changing cell composition. Transcripts of *FAR1*, the rate-limiting reductase, did not fall, consistent with its control at the protein level, but FAR1 protein fell in 8 of 9 ageing macaque tissues. The decline was confirmed in an independent human lifespan resource and was specific to mammals, absent from worm, fly and fish. It was not confined to vulnerable hotspots: spatial transcriptomics placed it in all 40 niches of ageing mouse brain, kidney, heart and spleen, and it replicated in three independent brain cohorts (meta p = 4 × 10⁻⁴). In the brain the loss was glial. It occurred within oligodendrocytes, oligodendrocyte precursors, microglia and astrocytes but not neurons, and the same glia became inflamed as microglia shifted towards disease-associated and interferon-responsive states. Finally, plasmalogen-deficient *Gnpat*-knockout brains had fewer astrocytes and fragmented mitochondria, and plasmalogen supplementation restored mitochondrial networks in inflamed astrocytes. Declining plasmalogen synthesis is therefore a pervasive, glial feature of mammalian ageing that links inflammation to mitochondrial damage.

**Highlights:**

- Transcripts of the committing enzymes *GNPAT* and *AGPS* and of the reductase *FAR2* decline with age in 105 human, macaque and mouse tissue models.
- The decline is not explained by changing cell composition and replicates in an independent human lifespan resource.
- It is specific to mammals and absent from worm, fly and fish.
- Spatial transcriptomics finds the decline throughout ageing mouse tissues, not only in vulnerable hotspots.
- In the brain the loss is glial: it occurs within oligodendrocytes, their precursors, microglia and astrocytes, not neurons.
- The same glia become inflamed, and the rate-limiting reductase FAR1 falls at the protein level in ageing macaque tissues.
- Plasmalogen deficiency fragments brain mitochondria, and plasmalogen supplementation protects astrocyte mitochondria.

## Introduction

Plasmalogens are ether phospholipids whose synthesis begins in the peroxisome, where FAR1 or FAR2, GNPAT and AGPS commit fatty alcohols to the ether-lipid route^1,2^. They make up to a fifth of human phospholipid and are enriched in brain, heart and immune cells, and in myelin^1,3^. Their loss matters for the two most consistently reported features of ageing, mitochondrial dysfunction and chronic low-grade inflammation^4–6^. Without ether lipids, nerve terminals respire less and mitochondrial fission fails^7,8^, and an age-dependent decline of plasmalogen biosynthesis impairs stress-induced mitochondrial responses^9^. In glia, inflammation lowers plasmalogens through NF-κB-dependent repression of GNPAT, and plasmalogen deficiency in turn amplifies NF-κB and TLR4 signalling^10–13^. Plasmalogens fall in the serum of older people^14,15^, brain plasmalogens are depleted at the mildest stages of Alzheimer’s and Parkinson’s disease^16–18^, and supplementation reduces microglial inflammation in aged mice^19^.

Three questions have remained open. First, whether the expression of plasmalogen-synthesis genes declines in normally ageing tissue across species, and whether it does so within cells rather than through changing cell mixtures^20^. Second, where in a tissue and in which cells the decline occurs: in a few vulnerable regions, such as the white matter that dominates inflammatory brain ageing^21,22^, or throughout, and in neurons or in the glia that become inflamed with age. Third, whether losing plasmalogens is enough to disturb brain mitochondria, as the fission defects of ether-lipid-deficient cells and flies suggest^7–9^.

We addressed these questions with bulk, single-cell and spatial transcriptomes of eight species, lipids and proteins measured in the same macaques, an independent human lifespan resource, and experiments in plasmalogen-deficient *Gnpat*-knockout mice and in inflamed astrocytes supplemented with plasmalogen. Every statistic is compared with a null built from the same data, and every cohort is reported whatever its outcome. The data identify plasmalogen-synthesis decline as a conserved, spatially pervasive and, in the brain, glial feature of mammalian ageing, and show that plasmalogen loss alone is enough to fragment brain mitochondria. How the decline relates in time and mechanism to mitochondrial decline, inflammation, epigenetic ageing and geroprotective interventions is analysed in a companion paper.

## Results

### Plasmalogen-synthesis transcripts decline across mammalian tissues, beyond cell composition

**Fig. 1** summarises the result. Across eight species, plasmalogen-synthesis expression falls with age only in mammals, where tissue transcripts before and after cell-composition control, FAR1 protein in macaque (8 of 9 tissues), every spatial niche of four ageing mouse tissues (40 of 40) and glial cell types point the same way (**Fig. 1A**); the mouse brain plasmalogen lipid fraction by middle age and adult human cortex cell types are the exceptions. **Fig. 1B** sketches the four findings developed below.

**Figure 1.**
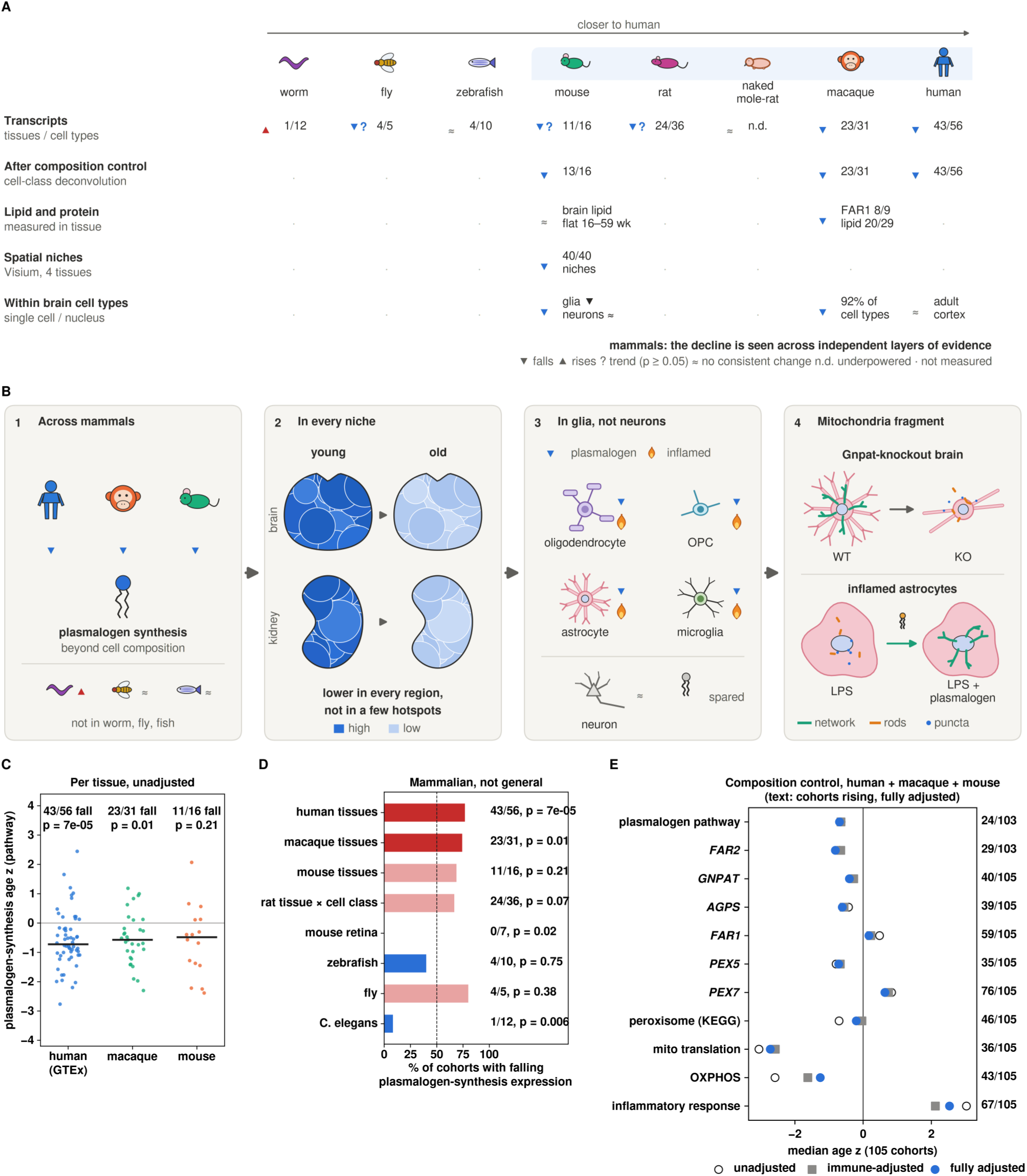
Plasmalogen-synthesis expression declines across mammals: overview. **A,** Evidence matrix: each line of evidence for falling plasmalogen synthesis in eight species, ordered by evolutionary distance from human (shaded, mammals). Transcripts: cohorts with falling plasmalogen-synthesis expression / cohorts tested (mammals, bulk tissues, unadjusted; rat, tissue × cell class; worm, fly and zebrafish, pre-specified single-cell-type cohorts). After composition control: the same after deconvolution. Lipid and protein: macaque FAR1 protein (8 of 9 tissues, p = 0.04) and ether-PE (plasmalogen) fraction (20 of 29, p = 0.06); mouse brain ether-PE flat from 16 to 59 weeks. Spatial niches: niches with a negative age slope in mouse brain, kidney, heart and spleen (40 of 40; not liver or lung; Fig. 4B). Within brain cell types: mouse glia but not neurons (Fig. 5A); macaque, 92% of units (sign-test q < 0.05; Fig. 2B); adult human cortex, no consistent fall (Fig. 2B). Glyphs: ▾/▴, most cohorts move in that direction with sign-test p < 0.05; ▾?, most fall but p ≥ 0.05; ≈, no consistent change; n.d., naked mole-rat, underpowered (T cells only); ·, not measured. **B,** Schematic of the four main findings: 1, mammalian and beyond cell composition (**A**, **C–E**; Fig. 2); 2, in every niche of ageing brain and kidney (Figs. 3, 4); 3, within glia, which also become inflamed, not neurons (Figs. 5, 6); 4, *Gnpat*-knockout astrocytes with fragmented mitochondria, and plasmalogen rescue in LPS-treated astrocytes (Fig. 8). Tissue colours show the direction of change only. **C,** Per-tissue pathway z, unadjusted; bar, median; text, tissues falling and sign-test p. **D,** Percentage of cohorts with falling plasmalogen-synthesis expression per species (cohorts as in **A**); text, cohorts falling / tested and sign-test p. **E,** Bulk atlas of 105 human (GTEx v10), macaque and mouse tissue age models: median age z (against abundance-matched random sets) for the plasmalogen pathway (Reactome wax and plasmalogen biosynthesis), the committed enzymes, PEX5 and PEX7, KEGG peroxisome and comparators, unadjusted (open), adjusted for immune fractions (grey) and fully adjusted for estimated cell composition (blue). Text, cohorts rising after full adjustment. Source data: Supplementary Data 1, 2 and 4.

The resource comprises 105 bulk tissue age models in human (GTEx v10; 56 tissues, 946 donors^23^), rhesus macaque (32 tissues, 17 animals^24^) and mouse (17 tissues, 56 animals^25^) (Supplementary Fig. 1A,B), extended by single-cell atlases of seven species, whose cell identities and marker genes are shown in Supplementary Figs. 2A–E and 3A–E, and by spatial transcriptomes (**Fig. 1A**; Methods). Gene sets were scored against random sets matched for transcript abundance, because per-gene age effects trend with abundance: the cross-species correlation of per-gene age effects tracks the product of the two datasets’ abundance trends (ρ = +0.51, p = 5 × 10⁻⁴, 42 comparisons; Supplementary Fig. 1C), and gene-level worm–mammal agreement is near zero (median ρ ≤ +0.024; Supplementary Fig. 1D).

Several analysis choices follow from controls (Supplementary Note 1; Methods): cross-species conservation is mapped at pathway rather than gene resolution, agreement is tested by sign tests over cohorts, age effects are estimated within tissue, and the flat macaque signal is not a lack of power. At pathway resolution, the most conserved of 759 shared terms are signalling, lysosome, endocytosis and phagocytosis terms that rise with age and mitochondrial terms (OXPHOS, mitochondrial translation, ATP synthesis) that fall in most species (Supplementary Figs. 4 and 5A); conservation grows with effect size (Supplementary Fig. 5B); and interferon, cGAS–STING and chronic-inflammation programmes rise and chaperones fall in all three mammals (Supplementary Fig. 5C).

Bulk tissue changes its cell mixture with age, so every age model was refitted with estimated cell-class pro-portions (11 classes, 440 markers, no plasmalogen enzyme among them; **Fig. 1E**; Supplementary Fig. 6A, comparator programmes). The organelle-wide peroxisome decline is compositional: KEGG peroxisome loses its signal under adjustment (Supplementary Fig. 6B). The plasmalogen-synthesis branch does not. The Reactome wax and plasmalogen biosynthesis pathway rises in only 24 of 103 cohorts after full adjustment (p = 5 × 10⁻⁸; Supplementary Fig. 6B), and transcripts of its committed enzymes fall: *FAR2* (29/103 rising, p = 10⁻⁵), *GNPAT* (40/105, p = 0.02) and *AGPS* (39/105, p = 0.01). *FAR1* transcripts do not fall (59/105), consistent with control of FAR1 at the protein level by cellular plasmalogens^26^. Per tissue, the pathway falls in 43 of 56 human (p = 7 × 10⁻⁵), 23 of 31 macaque (p = 0.01) and 11 of 16 mouse tissues (**Fig. 1C**). Brain shows the steepest decline; refitted against a primate brain-cell reference that captures falling neuronal and rising glial fractions (Supplementary Fig. 7B), the pathway rises in only 2 of 16 brain cohorts and *FAR2*, *GNPAT* and *AGPS* in 2–3 of 16 (**Fig. 2A**; Supplementary Fig. 7A). An earlier between-cohort adjustment, which regresses each cohort’s z on its immune-marker shift, likewise shows that the decline is not explained by between-cohort immune infiltration: there, inflammatory, complement and interferon-γ pathways (ρ ≥ 0.64 with the immune shift) lose most of their signal, whereas OXPHOS, mitochondrial translation and p53 keep more than half (Supplementary Data 1, sheet S10_composition).

**Figure 2.**
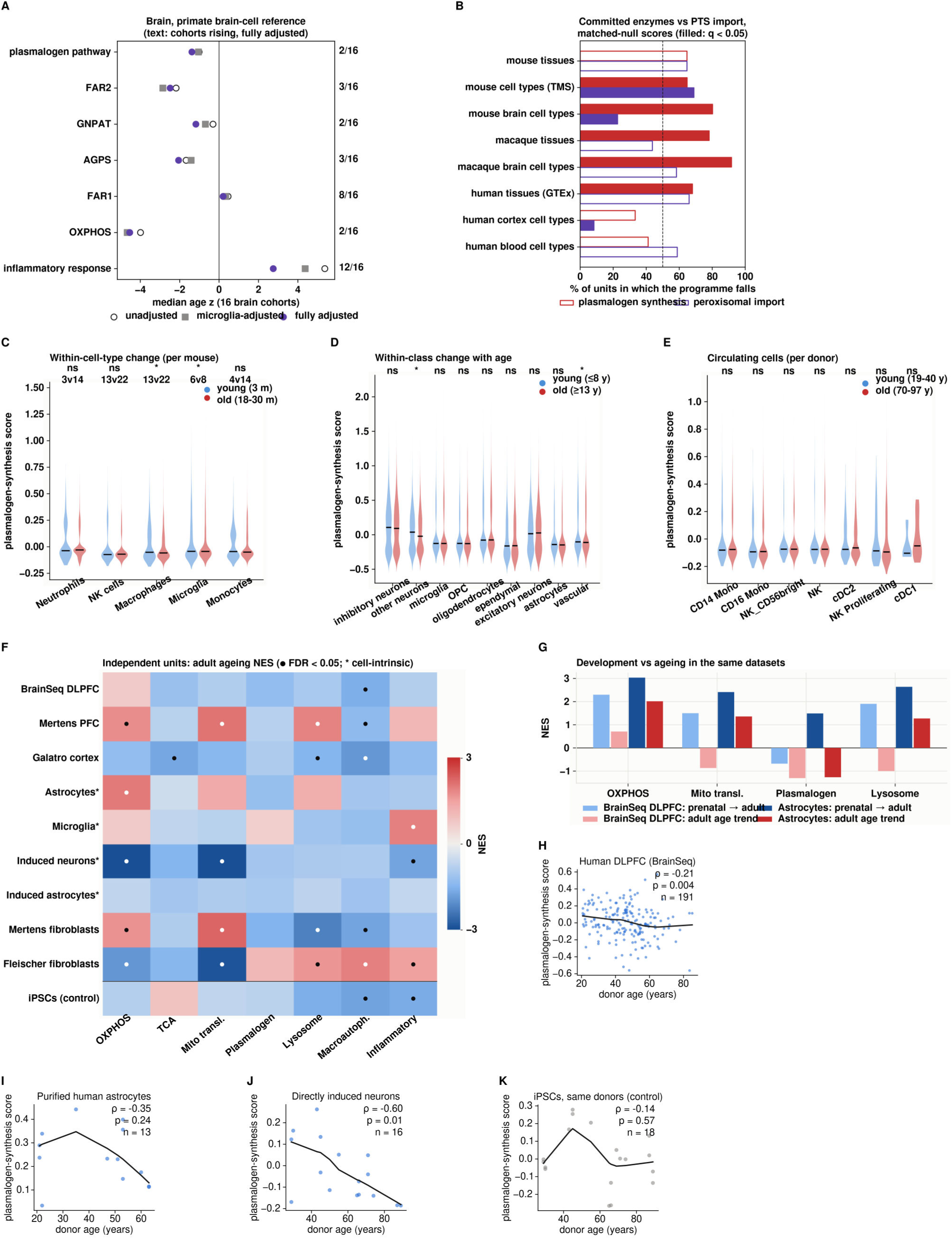
The decline persists beyond cell composition, within single cell types and in an independent human lifespan resource. **A,** The 16 brain cohorts refitted against a primate brain-cell reference (9 classes): unadjusted, microglia-adjusted and fully adjusted; text, cohorts rising. **B,** Matched-null scores in a second, independent implementation on donor- and cell-type-level pseudobulk: percentage of units in which committed plasmalogen synthesis (red) or peroxisomal import (purple) falls, per dataset; filled, sign-test q < 0.05. **C–E,** Score of the committed enzymes GNPAT, AGPS, FAR1 and FAR2 within single cell types, young versus old (violins, cells; tests per animal or donor, Welch’s t-test; *, p < 0.05; ns, not significant): **C,** mouse immune cells (Tabula Muris Senis; 3 versus 18–30 months); **D,** macaque brain nuclei (≤ 8 versus ≥ 13 years); **E,** human blood (OneK1K; 19–40 versus 70–97 years). **F–K,** Independent human lifespan resource: eight public bulk RNA-seq datasets (4,513 samples, 8 post-conception weeks to 102 years) analysed as 25 dataset × cell-type units with limma-voom and pre-ranked GSEA on the moderated t (full resource: Supplementary Figs. 12–15). **F,** Adult ageing normalised enrichment score (NES) per independent unit and in iPSCs for OXPHOS, TCA cycle, mitochondrial translation, plasmalogen synthesis, lysosome, macroautophagy and inflammatory response; ●, FDR < 0.05 (Benjamini–Hochberg); *, cell-intrinsic unit. **G,** NES for development (prenatal to adult) and for the adult age trend in BrainSeq DLPFC and purified astrocytes. **H–K,** Per-sample plasmalogen-synthesis module score (mean z of member genes) against donor age in adult donors (≥ 20 years) of BrainSeq dorsolateral prefrontal cortex (**H**; n = 191, Spearman ρ = −0.21, p = 0.004), purified human astrocytes (**I**; n = 13, ρ = −0.35, p = 0.24), directly induced neurons (**J**; n = 16, ρ = −0.60, p = 0.01) and iPSCs reprogrammed from the same donors (**K**; n = 18, ρ = −0.14, p = 0.57); line, LOWESS. These panels are descriptive; the unit-level test is the GSEA of **F**. Adults only are shown because prenatal samples drive an apparent rise in the astrocyte unit. Source data: Supplementary Data 1 and 4.

A second, independent re-implementation of the scoring on donor- and cell-type-level pseudobulk confirms the direction and separates the committed enzymes from peroxisomal import (**Fig. 2B**; per unit, Supplementary Fig. 9). In this matched-null atlas of 44 programmes, plasmalogen synthesis rises in only 8–35% of units in six datasets, whereas interferon programmes rise in most units of every dataset (Supplementary Fig. 8); 5–32% of units change direction between a simple and the matched-null score (15% for plasmalogen synthesis), and the median plasmalogen-synthesis z is negative in all three mammals (Supplementary Fig. 10A,B). Committed plasmalogen synthesis falls in most units of six of eight datasets — mouse cell types (65%), mouse brain cell types (80%), macaque tissues (78%), macaque brain cell types (92%) and human tissues (68%), each sign-test q < 0.05, and mouse tissues in direction (65%) — but not within adult human cortex or blood cell types. The PTS-import receptors PEX5 and PEX7 behave differently (PEX7 rises in 31 of 32 macaque tissues), which is why they are kept out of the committed-enzyme set; with them included, a simple score inverts the macaque result. Scored separately from import, plasmalogen synthesis falls in 23 of 32 macaque tissues before and after composition adjustment (p = 0.02) and in 11 of 12 macaque brain cell types (p = 0.006), against 20 of 32 with the simple score and 14–15 of 32 with PEX5 and PEX7 included (Supplementary Fig. 11A,B; reproduction check, Supplementary Fig. 11C); per gene, *FAR2*, *GNPAT* and *AGPS* fall in most tissues and *PEX7* rises (Supplementary Fig. 11D), and every brain cell type but microglia declines (Supplementary Fig. 11E).

Within single cell types, the committed-enzyme score falls in mouse microglia (p = 0.01) and macrophages (p = 0.02) and in macaque brain vascular cells (p = 0.01) and cerebellar and midbrain neurons (p = 0.03), with a trend in astrocytes (p = 0.08), but not in any circulating human NK-cell, monocyte or dendritic-cell subset (**Fig. 2C–E**). The decline is mammalian (**Fig. 1D**). Under analysis plans fixed before the data were examined, plasmalogen-synthesis expression falls in rat tissue × cell-class cohorts in direction (24/36; p = 0.07), but not in zebrafish (4/10), fly (4/5, too weakly expressed to score in most cell types) or mouse retina (0/7), and the worm ether-lipid pathway rises (11 of 12 cohorts; Supplementary Fig. 6C; Methods). Details per species are given in Supplementary Note 1.

### The decline replicates in an independent human lifespan resource and within purified and converted cells

Because the human arm rests on GTEx, we tested the decline in a second human resource of eight public datasets with explicit age groups (4,513 samples from 8 post-conception weeks to 102 years), analysed as 25 dataset × cell-type units (**Fig. 2**; Methods; 19 focus modules in every unit for development and ageing, Supplementary Fig. 12A,B). Gene-level ageing signatures agree between bulk brain series (BrainSeq versus GTEx frontal cortex, ρ = 0.70) but little between purified, converted and cultured cells, whose ageing signals are small or cell-type specific (Supplementary Fig. 13). Nine units use data not used elsewhere in this study; four of them are cell-intrinsic — purified astrocytes and microglia and directly induced neurons and astrocytes, in which a change cannot come from changing tissue composition — and iPSCs reprogrammed from the induced-neuron donors serve as a negative control because reprogramming resets age. The 14 GTEx v8 units are not independent, because their donors are in GTEx v10, and serve as a consistency check.

The plasmalogen-synthesis pathway rises in only 2 of 9 independent units, 1 of 4 cell-intrinsic units and 1 of 14 GTEx v8 brain and fibroblast units (Supplementary Fig. 12B; Supplementary Data 1). It falls in purified astrocytes (normalised enrichment score −1.3), directly induced neurons (−1.1) and induced astrocytes (−1.0) and rises in purified microglia (+0.6) (**Fig. 2F**). *FAR2* and *AGPS* rise in 2 of 9 independent units each and fall significantly in 8 and 6 of 14 GTEx brain regions; in the cell-intrinsic units *GNPAT* rises in none and *FAR2* and *AGPS* in one (microglia). None of these pathway scores is individually significant, and the enzyme-transcript falls are significant only in induced astrocytes (3 versus 3 donors; *GNPAT* t = −14.9, *AGPS* −10.1, *FAR2* −4.9, *FAR1* −23.3), so the replication in single cell types is one of direction. Per-sample scores show the same pattern across adult donors: the plasmalogen-synthesis score falls with age in BrainSeq dorsolateral prefrontal cortex (ρ = −0.21, p = 0.004, 191 adults) and in directly induced neurons (ρ = −0.60, p = 0.01), in direction in purified astrocytes (ρ = −0.35, p = 0.24), and not in iPSCs from the same donors (ρ = −0.14, p = 0.57) (**Fig. 2H–K**). Across the whole lifespan of BrainSeq DLPFC, mitochondrial and lysosomal modules rise steeply in development and then plateau or decline, whereas plasmalogen synthesis peaks in adolescence and drifts down through adulthood (Supplementary Fig. 14); in directly induced neurons, whose composition is fixed, the modules follow donor age (Supplementary Fig. 15A). Reprogramming erases the age signal more generally: 259 genes differ between older and newborn donors in fibroblasts, 19 in induced neurons and none in iPSCs, which show no plasmalogen-synthesis or OXPHOS expression trend with age (Supplementary Fig. 15B).

In this resource inflammatory programmes rise in tissue but not within cell-intrinsic units, and from prenatal to adult life OXPHOS rises while plasmalogen-enzyme transcripts split, which is why the adult age trend is the primary contrast (**Fig. 2G**; Supplementary Note 1).

### The decline is spatially pervasive within ageing tissues

To ask where in a tissue the decline occurs, we analysed spatial transcriptomes under a pre-registered plan, with the mouse as the statistical unit (**Figs. 3, 4**; Methods). In the discovery cohort (10x Visium; brain, heart, kidney, liver, lung and spleen at 4, 17 and 26 months), 57 niches were defined from marker genes before any age test; in brain, 19 niches reproduce the anatomy at every age (**Fig. 3A,B**), as do the niches of heart, kidney and liver (Supplementary Fig. 16A–D, E–H and I–L) and of lung and spleen (Supplementary Fig. 17A–D and E–H). Tissue composition changed little. Among niches, the kidney immune infiltrate expanded (+0.75, q = 0.028) and the brain striatum fraction fell, probably with section level (Supplementary Fig. 18A–F); among deconvolved cell types, brain oligodendrocytes rose and spleen T cells fell while mature NK T cells rose (Supplementary Fig. 19A–F); per-mouse trajectories of the twelve largest changes are shown in Supplementary Fig. 20A–L. Hepatic stellate and plasmacytoid dendritic cells increased in liver with both abundance methods agreeing.

**Figure 3.**
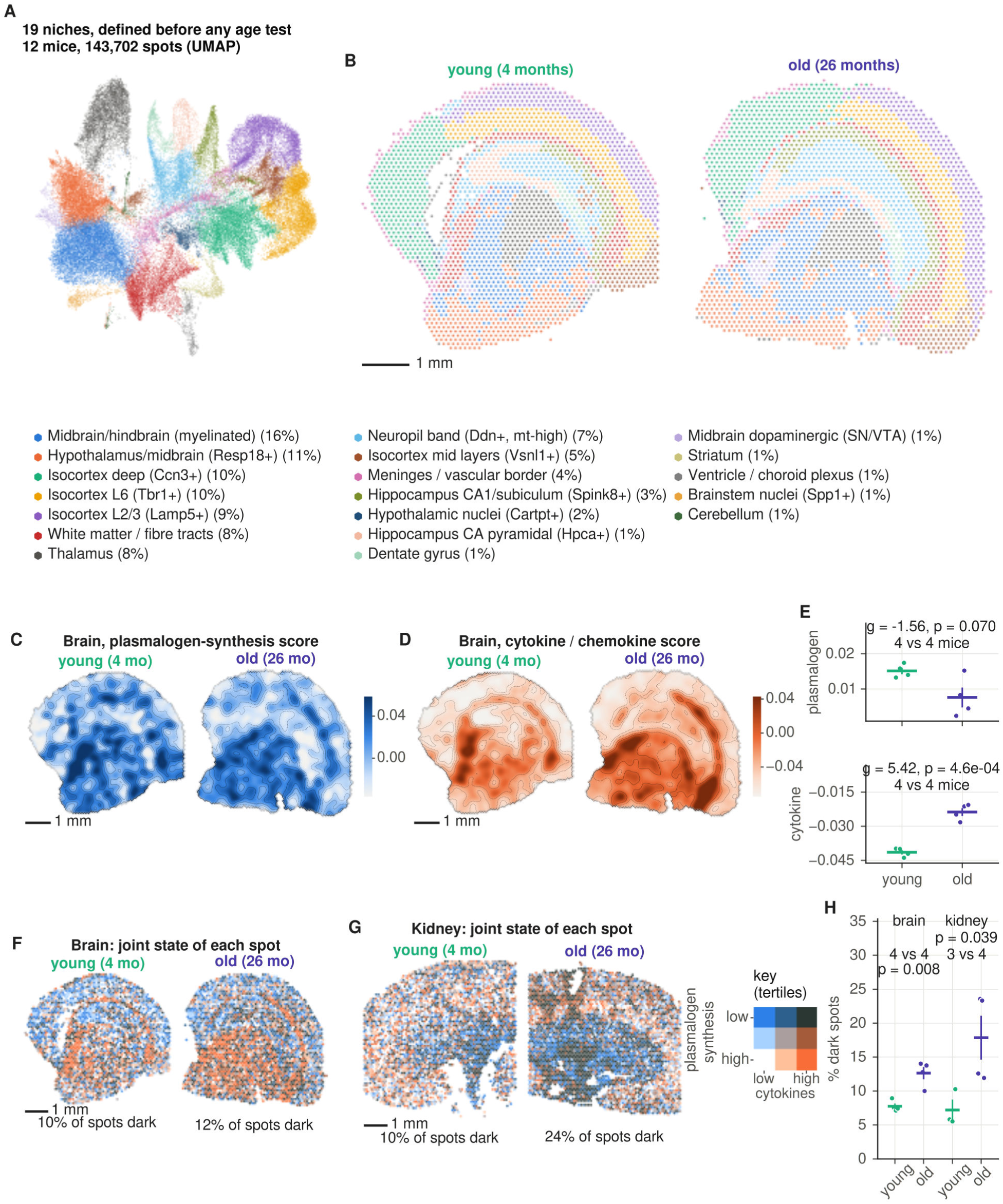
A spatial atlas of the ageing mouse brain and kidney. Visium discovery cohort GSE283283 (male mice). **A,** UMAP of 143,702 brain spots from 12 mice (4, 17 and 26 months) coloured by the 19 niches defined from marker genes before any age test (Supplementary Data 5, sheet niche_markers). **B,** The niches on one representative section of a young (4 months) and an old (26 months) mouse; legend, niche and share of spots. **C,D,** Plasmalogen-synthesis (16-gene focus set; blue) and cytokine/chemokine (orange) spot scores, each relative to expression-matched control genes, on the same young and old sections (Gaussian smoothing over 130 µm; contours, score quintiles; colour scale shared between ages). **E,** Mean score per mouse, youngest (3.9 months) versus oldest (25.8 months), 4 versus 4 mice; line, mean; whisker, SEM; Hedges’ g and Welch p. **F,G,** Joint state of every spot in brain (**F**) and kidney (**G**): each 55-µm spot coloured by its tertile of plasmalogen-synthesis score (blue, low) and cytokine score (orange, high), tertiles over all young and old spots of the tissue; dark, low plasmalogen synthesis and high cytokines; text, share of dark spots on the section. **H,** Per-mouse percentage of dark spots (brain 4 versus 4, kidney 3 versus 4 mice); Welch p. Sections are representative (the section with most spots per age, the same coronal level for brain); all tests are per mouse. Panels C–H are secondary analyses (not pre-registered). Niche maps of the other tissues: Supplementary Figs. 16 and 17. Source data: Supplementary Data 5.

**Figure 4.**
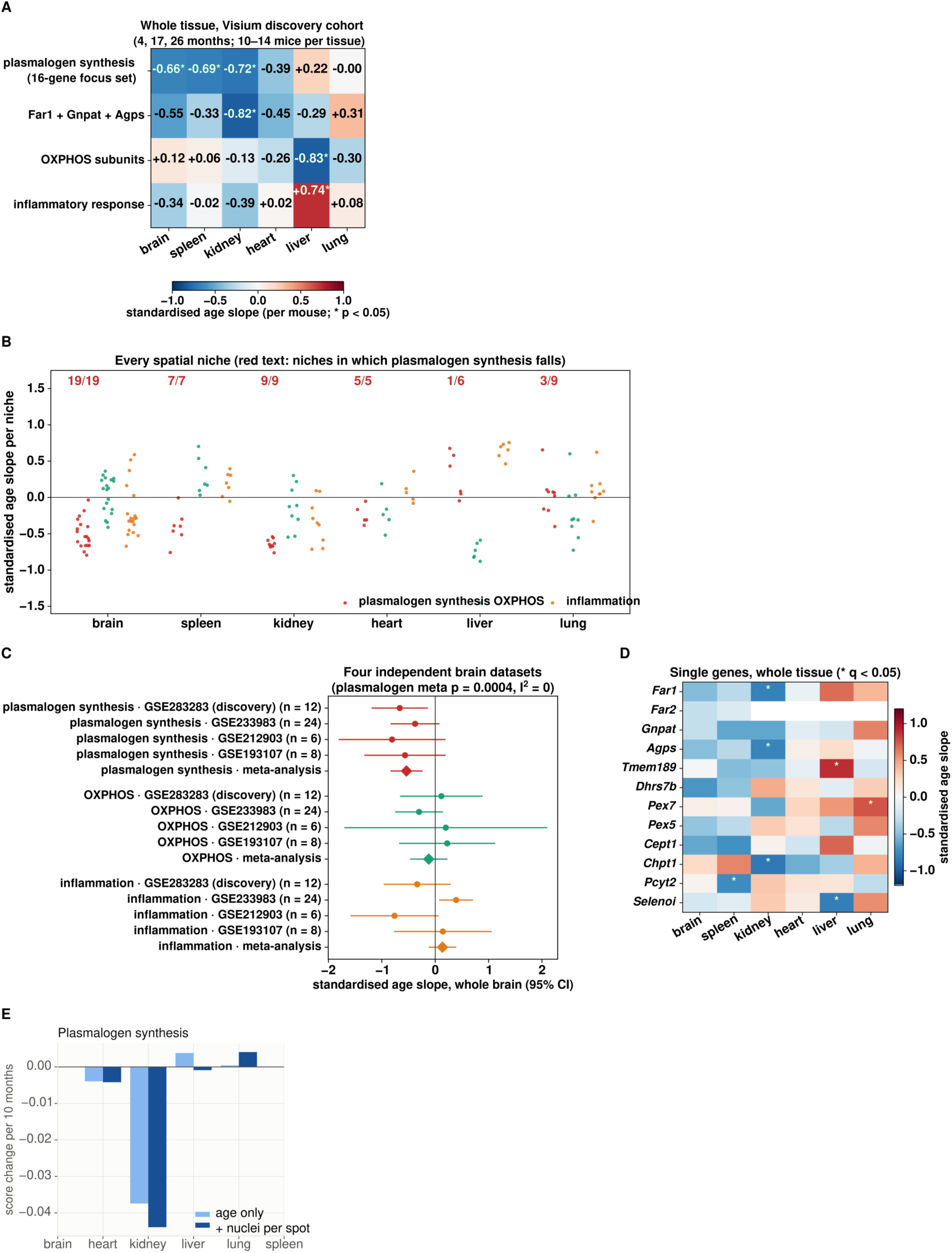
The decline is spatially pervasive within ageing tissues and not explained by cellularity. Spatial transcriptomics (10x Visium), discovery cohort GSE283283 (male mice aged 4, 17 and 26 months; 10–14 mice per tissue), analysed under a pre-registered plan with the mouse as statistical unit. **A,** Whole-tissue standardised age slope of the mean spot score (relative to expression-matched control genes) for plasmalogen synthesis (16-gene focus set), the committed trio Far1 + Gnpat + Agps, OXPHOS subunits and inflammatory response; * p < 0.05. **B,** Standardised age slopes of every niche (57 niches defined from marker genes before any age test); red text, niches in which plasmalogen synthesis falls. **C,** Whole-brain age slopes (95% confidence intervals) in the discovery cohort and three independent Visium datasets (GSE233983, 24 mice of both sexes, sex-adjusted; GSE212903; GSE193107), with inverse-variance meta-analysis (diamonds). **D,** Single-gene standardised age slopes per tissue; * q < 0.05 (Benjamini–Hochberg). **E,** Mouse-level age slopes of the programme scores with and without mean nuclei per spot (segmented from the H&E images) as covariate, and a spot-level mixed model (score ∼ age + log nuclei + niche + (1 | mouse)); the kidney plasmalogen-synthesis slope is unchanged after adjustment (p = 0.026 → 0.007); pre-registered addendum. Source data: Supplementary Data 4 and 5.

In representative sections of the oldest brains the plasmalogen-synthesis score was lower across the section rather than in a few regions, while the cytokine/chemokine score rose (**Fig. 3C,D**); per mouse, the plasmalogen-synthesis score fell (Hedges’ g = −1.56, p = 0.07) and the cytokine score rose (g = +5.42, p = 4.6 × 10⁻⁴; 4 versus 4 mice; **Fig. 3E**). Classifying every spot by both scores, spots with low plasmalogen-synthesis and high cytokine scores became more frequent with age in brain (7.8% to 12.6% of spots per mouse, p = 0.008) and kidney (7.2% to 17.9%, p = 0.039), where they formed contiguous territories (**Fig. 3F–H**; descriptive), and in spleen (about 8% to 18%, p = 0.037), but not significantly in heart (about 8% to 13%, p = 0.14) (Supplementary Fig. 21M–O and P–R). In the oldest versus youngest mice, plasmalogen-synthesis effect sizes were g = −2.14 in spleen (p = 0.009), −1.80 in kidney (p = 0.08) and −1.10 in heart (p = 0.22), against +0.45 in liver and +0.05 in lung, and every tissue with a falling score also raised its cytokine score, brain most (g = +5.4; Supplementary Fig. 22A). The spot clouds of brain and spleen moved towards low plasmalogen-synthesis and high cytokine scores, unlike those of liver and lung (Supplementary Fig. 22B–G), and nearly every spleen, kidney and heart niche moved the same way (Supplementary Fig. 23B–D), as did most brain cell types at cell resolution (Supplementary Fig. 23A).

Plasmalogen-synthesis expression fell at whole-tissue level in brain (standardised slope −0.67, p = 0.013), spleen (−0.69, q = 0.021) and kidney (−0.72, p = 0.026), in direction in heart (−0.39), and not in liver (+0.22) or lung (0.00) (**Fig. 4A**). The decline was not confined to vulnerable regions: it was negative in all 19 brain niches, all 9 kidney niches, all 5 heart niches and all 7 spleen niches, but in only 1 of 6 liver and 3 of 9 lung niches (**Fig. 4B**; brain and heart, Supplementary Fig. 24A,B; kidney, liver, lung and spleen, Supplementary Fig. 24C–F); in liver the periportal niche rose, with OXPHOS falling and inflammatory response rising across liver niches (Supplementary Fig. 24D). Grouped by dominant deconvolved cell type, spots of brain oligodendrocytes, OPCs and vascular cells, heart cardiomyocytes, kidney proximal-tubule cells and spleen B cells and erythroblasts declined, in direction, whereas brain choroid plexus and lung type II pneumocytes did not (Supplementary Fig. 25A–C and D–F); whole-tissue slopes are shown in **Fig. 4A**. Against the pre-registered criteria, the plasmalogen-synthesis hypothesis was supported in brain (replicated) and spleen, directional in heart and kidney, and not supported in liver or lung. The committed trio Far1, Gnpat and Agps alone showed the same pattern, most strongly in kidney (−0.82; **Fig. 4A**). The brain result replicated in all three independent Visium datasets (GSE233983, 24 mice of both sexes, −0.38; GSE212903, −0.81; GSE193107, −0.57), with an inverse-variance meta-analysis slope of −0.54 (p = 4.4 × 10⁻⁴; I² = 0; **Fig. 4C**); in the largest replication cohort, 15 of 17 transferred niches declined, significantly in deep cortical layer 6, the neuropil band, white matter, myelinated midbrain and hindbrain, and meninges. At the gene level, the rate-limiting enzymes *Far1* (q = 0.048) and *Agps* (q = 0.014) and the downstream *Chpt1* (q = 0.005) fell in kidney, and *Pcyt2* in spleen (q = 0.001), whereas in brain *Far1*, *Agps*, *Gnpat*, *Dhrs7b* and *Cept1* fell in direction (**Fig. 4D**). Per niche, *Cept1* fell in brainstem and mid-cortical niches and *Far1* in white matter while *Pex7* rose in hippocampal pyramidal cells, and *Far2* fell in stressed heart myocardium (Supplementary Fig. 26A,B), and *Agps*, *Chpt1* and *Far1* across kidney tubule niches and *Pcyt2* across spleen niches, whereas *Pex7* rose in lung (Supplementary Fig. 27A–D). Maps of the plasmalogen-synthesis score in two young and two old sections of each tissue, with per-mouse niche means, show whole-section shifts rather than changes confined to single niches in brain and heart (Supplementary Fig. 28A–E and F–J), kidney and liver (Supplementary Fig. 29A–E and F–J) and lung and spleen (Supplementary Fig. 30A–E and F–J), as do maps of the most-decreased pathway genes in brain and kidney (Supplementary Fig. 21A–F and G–L); in the oldest-versus-youngest contrast no single pathway gene passed q < 0.05.

Two controls address what the spot score measures. Nuclei segmented from the H&E images under a pre-registered addendum validate the niches and show little change in cellularity with age, and the kidney decline is unchanged after adjustment for nuclei per spot (p = 0.026 before, 0.007 after; **Fig. 4E**). Segmentation quality differed between tissues, and spleen sections failed registration and were excluded (nuclear density, area and stain saturation, Supplementary Fig. 31A–C). Nuclei per spot correlated only weakly with UMIs per spot within sections (median ρ −0.06 to +0.15; Supplementary Fig. 32A–E) but differed between niches as expected, dentate gyrus, choroid plexus and cerebellum carrying the most nuclei in brain and tubular niches in kidney (Supplementary Fig. 32F–J). Cellularity barely changed with age: in heart, kidney and lung only the lung pulmonary-vein myocardium lost nuclei (q = 0.008; Supplementary Fig. 31D–F), and brain, liver and spleen gave no estimate. Lymphocyte-like nuclei were most frequent in the kidney immune-infiltrate, lung pleural and lymphoid, and brainstem and white-matter niches (Supplementary Fig. 33A–E), and nuclear morphology changed little with age in heart and kidney (Supplementary Fig. 34A,B) or liver (whole tissue only; Supplementary Data 5, sheet image_morphology_age), whereas elongated nuclei became more frequent in four lung niches (q < 0.05; Supplementary Fig. 34C). The decline is relative to genes of similar expression level; absolute pseudobulk levels move less (brain −0.47, p = 0.18; spleen −0.64, p = 0.013). In spleen the decline is partly compositional, as B cells (35% to 20%, p = 0.07) and T cells (9.1% to 3.8%, p = 0.013) become fewer and plasma cells more numerous (Supplementary Fig. 35A). Attributing expression to cell types by regression, which places marker genes in their own cell type (Supplementary Fig. 35B), the spleen decline is not carried by one cell type: in spots dominated by erythroblasts or B cells the score falls without reaching significance (g = −1.1 and −0.7; Supplementary Fig. 35C), and gene changes per cell type are mixed, the largest being *Pcyt2* in plasma cells (p = 0.010; Supplementary Fig. 35D,E). In such dominated spots *Pcyt2* and *Chpt1* change in erythroblasts and *Gnpat* falls in B cells, each nominally (Supplementary Fig. 35F).

The inflammatory signature behaved differently. The Hallmark inflammatory response rose only in liver (+0.74, q = 0.009) and, in brain, in white matter in every dataset and in the ventricle/choroid plexus niche, whereas narrower inflammatory gene sets — complement (Hedges’ g = +10.3), interferon-α (+6.7), cytokine/chemokine (+5.4) and MHC (+3.7) — rose across the whole brain in the oldest mice (each p ≤ 0.005), whereas the Hallmark TNF-α/NF-κB set fell (−2.1, p = 0.02); liver raised the Hallmark inflammatory (+2.1), interferon-α (+3.0), complement and MHC sets, and spleen the cytokine (+2.0) and SASP (+1.8) sets (**Fig. 6A**). Thus the plasmalogen-synthesis decline, unlike the inflammatory signature concentrated in brain white matter^21,22^, is spread through every niche.

The pre-registered mitochondrial hypothesis was supported in liver only (OXPHOS −0.83, q = 0.009; Supplementary Note 1); how the programmes relate to one another spot by spot, between individuals and across tissues is analysed in a companion study.

### In the ageing brain, the decline is glial, not neuronal

Spots mix cells, so we asked which brain cells carry the decline using single-cell transcriptomes of the same strain, with the mouse as the unit (Tabula Muris Senis FACS brain, myeloid and non-myeloid cells sorted from the same mice, 3 versus 18–24 months, both sexes^27^; **Fig. 5**; Methods), because Stereo-seq cell bins (median 326 UMIs) do not resolve microglial states (results summarised schematically in **Fig. 1B**). The plasmalogen-synthesis score fell within oligodendrocytes (g = −2.72, q = 0.03), oligodendrocyte precursor cells (OPCs; −2.50, q = 0.008), microglia (−1.46, q = 0.03), astrocytes (−1.25, q = 0.05) and endothelial cells (−1.34, q = 0.05), but not within neurons (+0.16) or interneurons (−0.39) (**Fig. 5A**), while an inflammation composite rose most in microglia (**Fig. 5A**). Per mouse, astrocytes, oligodendrocytes, OPCs and endothelial cells lost plasmalogen-synthesis score (p = 0.035, 0.011, 0.001 and 0.035), oligodendrocytes and OPCs raised reactive and complement scores, and neurons raised reactive, complement and cytokine scores (Supplementary Fig. 36A). The committed-enzyme score (Far1 + Gnpat + Agps) fell in oligodendrocytes (−2.97, q = 0.008) and microglia (−1.85, q = 0.04).

**Figure 5.**
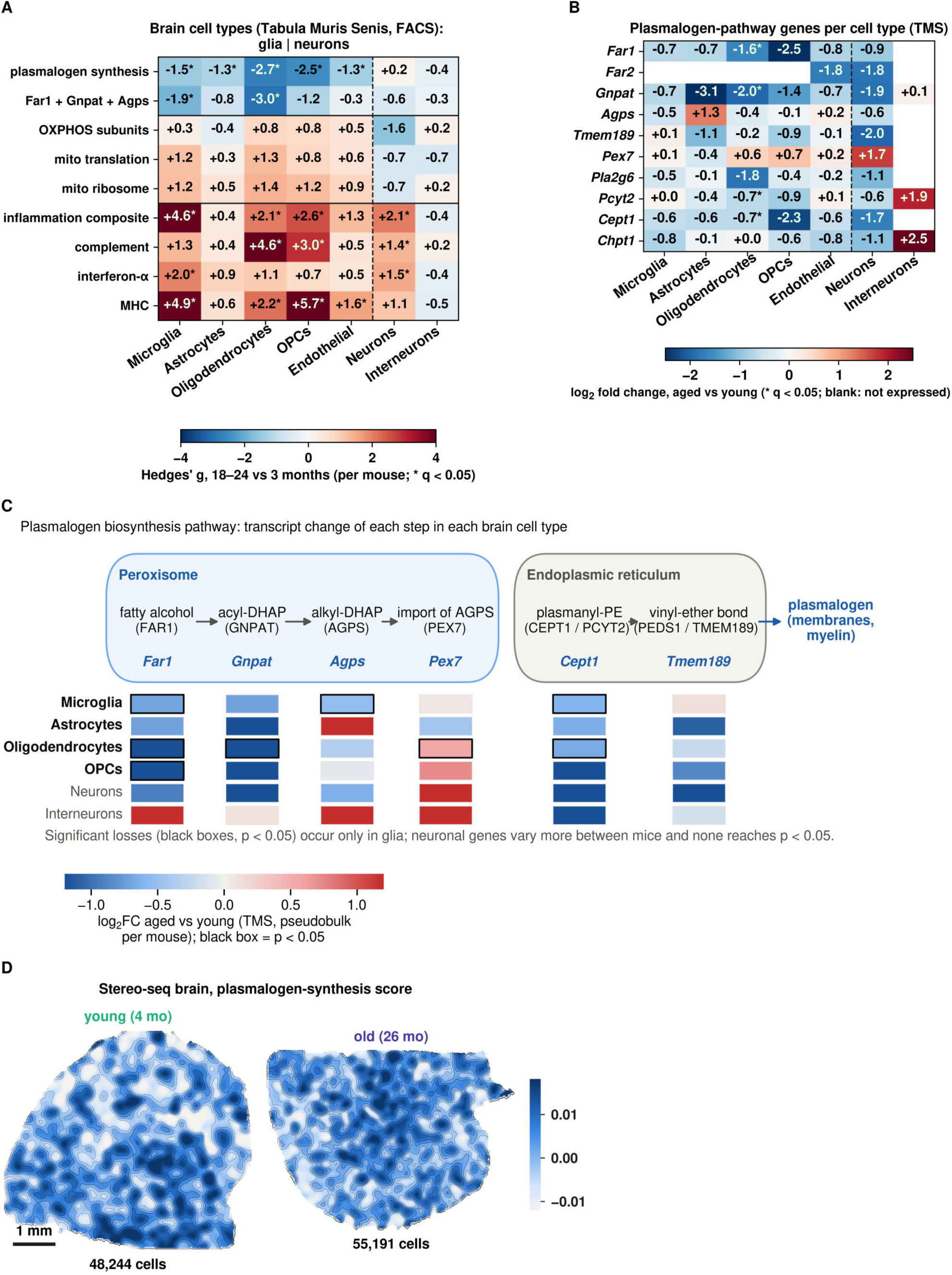
In the ageing brain the decline is glial, not neuronal. **A,** Brain cell types (Tabula Muris Senis FACS, Smart-seq2; 3 versus 18–24 months; 4–8 mice per group): Hedges’ g of programme scores per mouse; inflammation composite, mean z of reactive, complement and cytokine/chemokine scores; * BH q < 0.05 within programme; dashed line separates glial and vascular from neuronal types. **B,** log₂ fold change of plasmalogen-pathway genes per cell type (pseudobulk per mouse); blank, mean young log₂ CPM < 2; * q < 0.05. **C,** The plasmalogen biosynthesis pathway (peroxisomal FAR1, GNPAT, AGPS and PEX7 import; endoplasmic-reticulum CEPT1/PCYT2 and PEDS1/TMEM189) with, at each step, tiles giving the log₂ fold change of the gene in each cell type; black box, p < 0.05 (significant losses occur only in glia). **D,** Stereo-seq brain sections at cell resolution (GSE283283; one mouse per age, so descriptive): plasmalogen-synthesis cell score of a young (4 months) and an old (26 months) section (Gaussian smoothing over 90 µm; colour scale shared between ages); text, cells per section. Panels A–D are secondary analyses (not pre-registered). Source data: Supplementary Data 5.

Mapped onto the pathway, the losses lie at its committed steps. In oligodendrocytes the committed enzymes themselves lost expression — *Far1* (−1.63 log₂, q = 0.003) and *Gnpat* (−1.97, q = 0.035) — together with the downstream endoplasmic-reticulum enzymes *Pcyt2* and *Cept1* (q = 0.046), whereas neuronal changes were neither consistent nor significant (**Fig. 5B,C**). Mitochondrial programmes did not follow the same pattern within brain cells: OXPHOS and mitochondrial-translation scores did not fall within any glial type (g between −0.4 and +1.4, all q > 0.1) and fell only in direction in neurons (OXPHOS −1.6, q = 0.06) (**Fig. 5A**).

Two tissue-resolved views agree in direction but are descriptive. In Visium spots dominated by one deconvolved cell type, deconvolved proportions did not change and no comparison reached p < 0.1, reflecting the few dominated spots and mice (Supplementary Fig. 36B,C). In Stereo-seq brain sections at single-cell resolution (one mouse per age; 11 cell types resolved from 48,000–55,000 cells per section, Supplementary Fig. 37A–C), the plasmalogen-synthesis score was lower at 26 than at 4 months across the section (**Fig. 5D**), in 10 of 11 resolved cell types when cells were pooled and in 8 of 11 when sections were averaged, most in astrocytes (g = −1.89), oligodendrocytes (−1.13) and microglia (−0.79) (Supplementary Fig. 38A; pathway genes per cell type, Supplementary Fig. 38B). Neurons are the one cell class in which data types disagree: unchanged in sorted cells but lower in Stereo-seq cells of one mouse per age and nominally lower in neuron-rich Visium territory (**Fig. 6C**).

**Figure 6.**
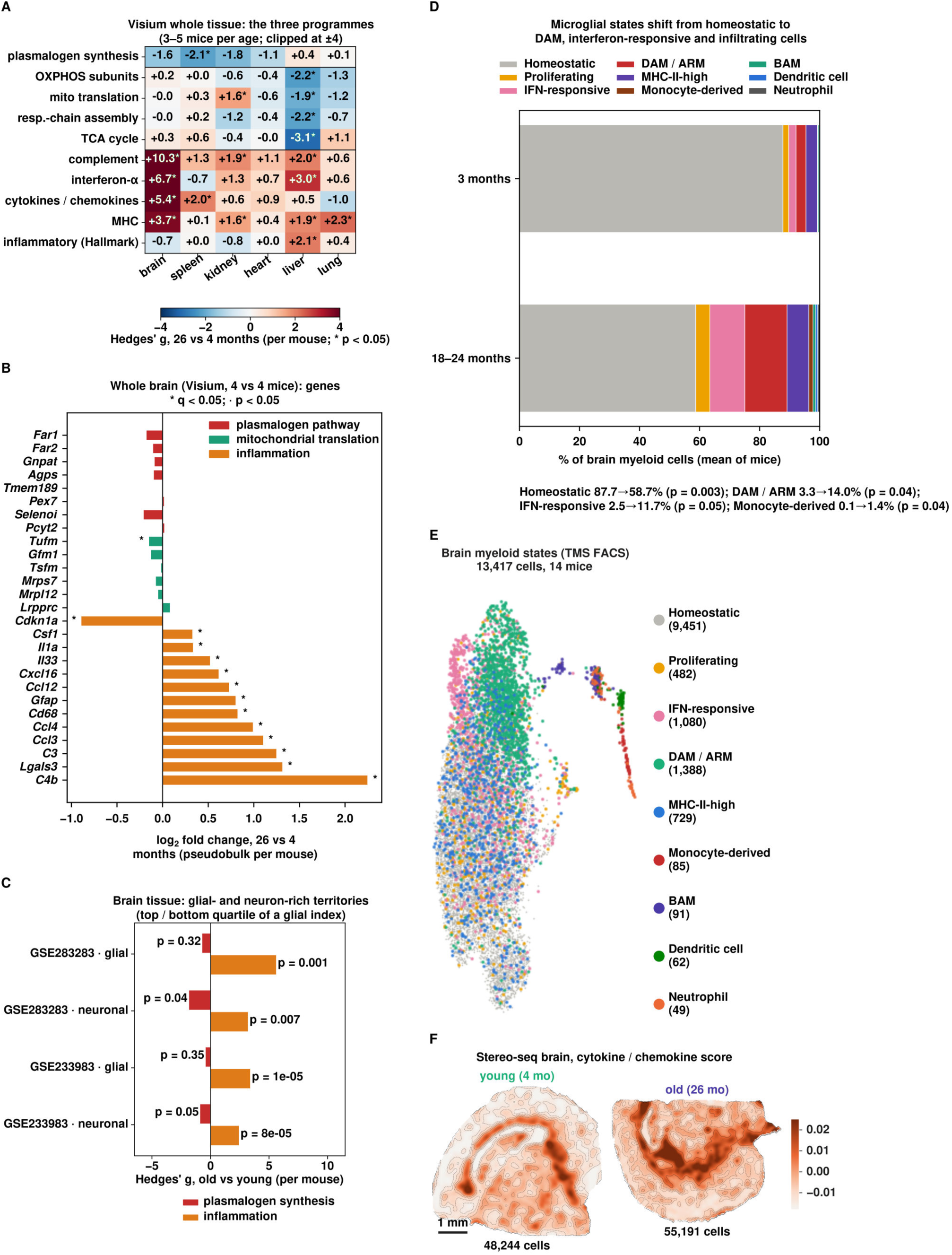
The same glia become inflamed and microglia change state. **A,** Visium discovery cohort (GSE283283), oldest (25.8 months) versus youngest (3.9 months) mice, 3–5 per age: Hedges’ g (per mouse) of the plasmalogen-synthesis score, five mitochondrial gene sets and five inflammatory gene sets per tissue; * Welch p < 0.05; colour clipped at ±4. **B,** Whole brain (Visium, 4 versus 4 mice): log₂ fold change (pseudobulk per mouse) of plasmalogen-pathway genes (red), mitochondrial-translation genes (green) and the inflammatory genes with q < 0.05 (orange); * BH q < 0.05, · p < 0.05. **C,** Visium brain territories in two cohorts (GSE283283, 4 versus 4; GSE233983, 10 versus 10 mice of both sexes): Hedges’ g of the plasmalogen-synthesis and inflammation scores in glia-rich and neuron-rich territories (top and bottom quartile of a glial index per section); text, Welch p. **D,** Composition of brain myeloid cells by state (Tabula Muris Senis FACS; mean of mice; states assigned from marker scores with thresholds fixed before age was examined); text, Welch p. **E,** UMAP of 13,417 brain myeloid cells (Tabula Muris Senis FACS, 14 mice) coloured by state; legend, cells per state. **F,** Stereo-seq brain sections (one mouse per age, descriptive): cytokine/chemokine cell score of a young (4 months) and an old (26 months) section (Gaussian smoothing over 90 µm; colour scale shared between ages). Source data: Supplementary Data 5.

### The same glia become inflamed, and microglia change state

The glia that lose plasmalogen-synthesis expression became inflamed. An inflammation composite of reactive, complement and cytokine genes rose in microglia (g = +4.57, q < 10⁻⁵), OPCs (+2.58) and oligodendrocytes (+2.09), with MHC transcripts rising in microglia (+4.9), OPCs (+5.7) and oligodendrocytes (+2.2) and complement in oligodendrocytes (+4.6) and OPCs (+3.0); neurons also raised the composite (+2.06) (**Fig. 5A**). Across the whole brain, C4b, Lgals3, C3, Ccl3, Ccl4, Cd68, Gfap, Ccl12, Cxcl16, Il33, Il1a and Csf1 transcripts rose (each q < 0.05; **Fig. 6B**), and in the spatial discovery cohort complement, interferon and MHC programmes rose most in brain among six tissues (**Fig. 6A**).

Among brain myeloid cells, homeostatic microglia fell from 88% to 59% (q = 0.03), whereas disease-associated (DAM/ARM; 3% to 14%), interferon-responsive (2.5% to 11.7%) and Ccr2⁺ monocyte-derived cells (0.14% to 1.37%) expanded (**Fig. 6D,E**). In the full state atlas of 13,417 myeloid cells (**Fig. 6E**; marker genes, Supplementary Fig. 39A), monocyte-derived cells (p = 0.038; Supplementary Fig. 39B), border-associated macrophages (p = 0.046), DAM/ARM (p = 0.045) and proliferating microglia (p = 0.011) rose from 3 to 24 months, interferon-responsive (p = 0.054) and MHC-II-high microglia in direction (Supplementary Fig. 39C–G). Monocyte origin is inferred from *Ccr2* and *Ly6c2* and is not lineage-traced. The plasmalogen-synthesis score fell within homeostatic and DAM/ARM microglia (g = −2.14 and −2.07, q = 0.03), and interferon-responsive microglia raised *Il1b* (+2.37 log₂, q < 10⁻⁴) and *Ccl5* (+7.0, q = 0.025) (Supplementary Fig. 39J); with age every microglial state moved towards lower plasmalogen-synthesis and higher cytokine scores, and the aged-only border-associated and monocyte-derived cells had the lowest plasmalogen-synthesis scores (Supplementary Fig. 39H,I). The Hallmark inflammatory score falls within states, so the tissue-level inflammatory rise reflects the shift between states. In brain tissue, monocyte-signature hotspots carried steeper cytokine gradients with age (p = 0.05 and 0.026) without a local dip in the plasmalogen-synthesis score (p = 0.39 and 0.075): the non-microglial macrophage signature rose with age (p = 0.003 and 2 × 10⁻⁵; Supplementary Fig. 40A–C), cytokines rose near hotspots in old mice (Supplementary Fig. 40F–H), and plasmalogen-synthesis scores did not dip there (Supplementary Fig. 40D,E,I). In tissue, inflammation rose most in glia-rich territories (g = +5.6 and +3.4 in two Visium cohorts), whereas the plasmalogen-synthesis score fell in both glia- and neuron-rich territories, nominally in the latter (−1.83, p = 0.04; replication −0.88, p = 0.054) (**Fig. 6C**; Supplementary Fig. 41A–F), and along the neuronal-to-glial axis inflammation rose most at the glial end (Supplementary Fig. 41G–I), as expected when 55-µm spots mix glia with neurons. At cell resolution the cytokine score was higher across the old section (**Fig. 6F**), with more microglia in the old section (1,377 versus 494) and similar astrocyte numbers (Supplementary Fig. 37D; one mouse per age, descriptive).

Exploratory analyses of the hypothesis that glia supply lipids to neurons (Supplementary Fig. 42A; Supplementary Note 1) found no glia-to-neuron signal that transcripts could resolve. Among lipid-carrier genes, *Apoe* rose in aged microglia (+2.2 log₂) and neurons (+5.7) (Supplementary Fig. 42B,C); no glia-to-neuron ligand–receptor pair passed q < 0.1 in single cells, the strongest being microglial *Apoe* to neuronal *Lrp1* (+1.93 log₂, p = 0.004, q = 0.15; Supplementary Fig. 43A–F). In tissue, *C3*–*C3ar1*, *Spp1*, *C1qa*–*Lrp1*, *Apoe*, *Il1a*, *Il1b* and *Csf1* pairs rose in co-expression with age, significantly in the larger replication cohort (Supplementary Fig. 44A–C), and the plasmalogen-synthesis score of the surrounding tissue was negatively, not positively, associated with synaptic genes, without an age interaction or microglial modification (Supplementary Fig. 42D–F). Transcripts cannot measure lipid transfer.

### Plasmalogen lipids and the rate-limiting enzyme fall in ageing macaque tissue

Transcripts are not lipid or enzyme levels^26,28^, so we examined lipids and proteins measured in the same 17 macaques as the transcriptome, in up to 30 tissues^24^. The ether-PE (plasmalogen) fraction — ether-linked minus diacyl PE, a measure independent of membrane content — fell with age in 20 of 29 tissues (p = 0.06; median t = −0.36), most strongly in cecum, pancreas, skin, hypothalamus and temporal and parietal cortex, whereas diacyl-PE did not (13/29, p = 0.71) (**Fig. 7A**; Methods). The rate-limiting reductase FAR1 fell at the protein level in 8 of 9 tissues where it was detected (p = 0.04), although its transcript did not fall across cohorts (**Fig. 1E**), consistent with control of FAR1 at the protein level^26^. OXPHOS proteins fell in 21 of 29 tissues (p = 0.02) and inflammatory and complement proteins rose in 22 of 28 (p = 0.004). In macaque cortex, plasmalogen-synthesis transcripts are flat but plasmalogen lipids fall in hypothalamus and temporal and parietal cortex; the youngest animals of the atlas are still maturing. In mouse brain, by contrast, the ether-PE fraction rose strongly during development and did not fall from 16 to 59 weeks within one batch (6/10 regions falling; p = 0.75), so the adult brain transcript decline is not mirrored by lipids by middle age (**Fig. 7B**).

**Figure 7.**
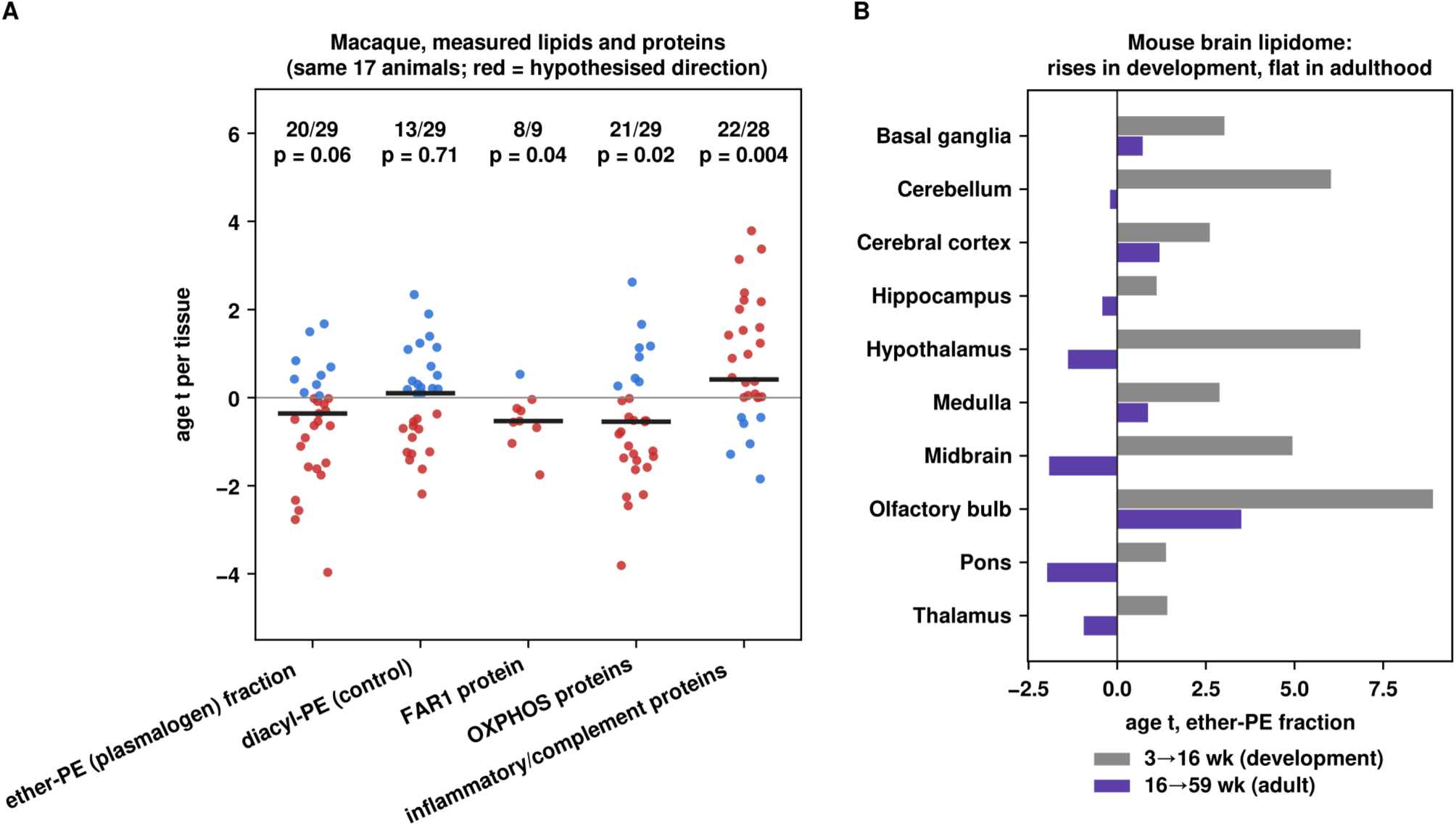
Plasmalogen lipids and FAR1 protein fall in ageing macaque tissue. **A,** Macaque (17 animals, same as the transcriptome; up to 30 tissues): age t per tissue for the ether-PE (plasmalogen) fraction, diacyl-PE (control), FAR1 protein, an OXPHOS protein score and an inflammatory/complement protein score; red, hypothesised direction; bar, median; text, tissues in the hypothesised direction / tested and sign-test p. **B,** Mouse brain lipidome: age t of the ether-PE fraction per region from 3 to 16 weeks (development) and from 16 to 59 weeks (adult; one batch). Source data: Supplementary Data 2 and 3.

Pre-specified brain arms mark the limits of the transcript decline within brain cells (Supplementary Data 3, sheet arms_vs_criteria; Methods). It is not seen within adult human prefrontal cortex cell types (PsychAD; 192 adults aged 20–89 years), although it is seen there in development. In the Allen mouse brain atlas it is seen within cell classes and, with the committed-enzyme score, within 80% of subclasses, but not robustly with the broader pre-specified focus set (15 of 22 subclasses, p = 0.13). Together with the absence of a fall within human blood cell types (OneK1K; **Fig. 2E**), these boundaries place the decline in mammalian tissue and in mouse and macaque cells.

### Plasmalogen deficiency fragments brain mitochondria, and plasmalogen protects astrocyte mitochondria from inflammation

Transcripts cannot show whether losing plasmalogens disturbs mitochondria, so we examined mice lacking GNPAT, the first committed enzyme of the pathway. *Gnpat*-knockout (KO) mice cannot make ether lipids, including plasmalogens, and develop male infertility, eye defects and myelination defects^29,30^. We compared the brains of male KO and wild-type (WT) mice from heterozygous intercrosses on a C57BL/6J background at 14 weeks of age (**Fig. 8**; Methods). Astrocytes, one of the glial types in which plasmalogen-synthesis expression falls with age (**Fig. 5A**), were sparser in KO brain: GFAP-positive astrocyte density was 24% lower (432 ± 20 versus 328 ± 11 per mm², P = 0.002) and the GFAP-positive area 25% lower (P = 0.043), with unchanged nuclear density (P = 0.33; 6 versus 6 images; **Fig. 8A,D**). Individual KO astrocytes had fewer, longer and more bipolar processes, a nominal difference (**Fig. 8B,C**; not significant after correction across 41 descriptors).

**Figure 8.**
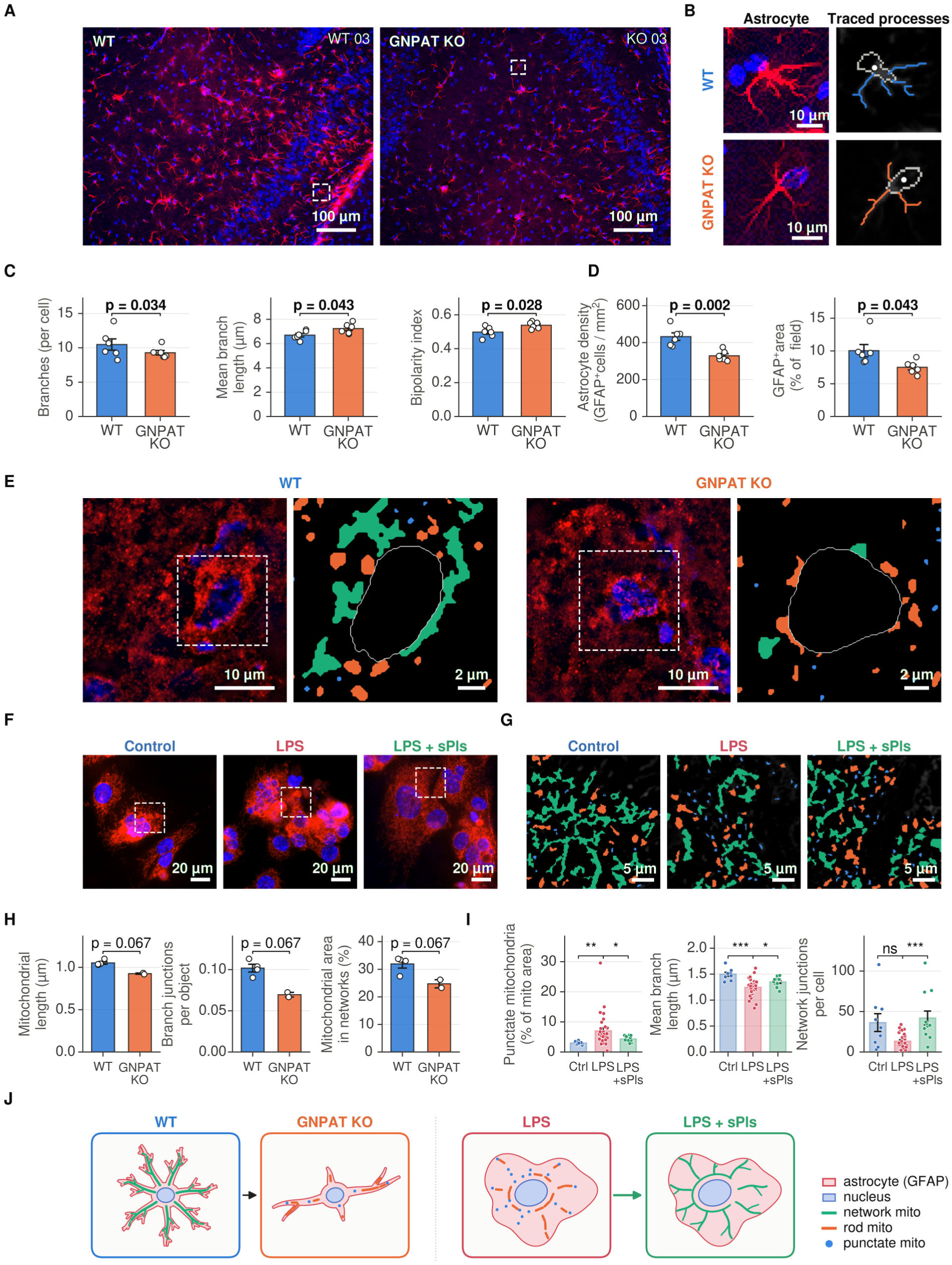
Plasmalogen deficiency fragments brain mitochondria, and plasmalogen protects astrocyte mitochondria from inflammation. **A–E,H,** Brains of male *Gnpat*-knockout (KO) and wild-type (WT) mice, C57BL/6J background, 14 weeks old. **A,** GFAP (red) and DAPI (blue), WT and KO sections (contrast set per image because GFAP exposure differed between genotypes; brightness is not compared); boxes, astrocytes enlarged in **B**. Scale bars, 100 µm. **B,** Enlarged astrocytes and their traced processes rooted at the soma (blue, WT; orange, KO). Scale bars, 10 µm. **C,** Per-astrocyte morphology: branches per cell, mean branch length and bipolarity index (0, radial; 1, aligned on one axis); mixed model with image as random intercept (618 WT, 651 KO astrocytes); P unadjusted (not significant after Benjamini–Hochberg correction across 41 descriptors). **D,** GFAP-positive astrocyte density and GFAP-positive area fraction; Welch’s t-test on image values. In **C, D**, bars, mean of image means ± SEM; circles, images (6 WT, 6 KO). **E,** VDAC1 (red) and DAPI (blue) in WT and KO tissue: a field with a perinuclear cell boxed (scale bar, 10 µm) and that cell’s mitochondria coloured by shape (blue, punctate; orange, rod; green, network; white outline, nucleus; scale bar, 2 µm). **F,** A1 astrocytes stained with MitoTracker Orange (red) and DAPI (blue) after control, LPS or LPS + scallop plasmalogen (sPls) treatment; boxes, regions enlarged in **G**. Scale bars, 20 µm. **G,** Mitochondria of the boxed regions coloured by shape as in **E**. Scale bars, 5 µm. **H,** Tissue mitochondrial length, branch junctions per object and network share of mitochondrial area; bars, mean ± SEM; circles, fields (4 WT, 2 KO); exact two-sided permutation test, for which 0.067 is the smallest attainable P with 4 versus 2 fields. **I,** A1 astrocytes: punctate share of mitochondrial area, mean network branch length and network junctions per cell; bars, mean ± SEM; dots, cells (control 9, LPS 30, LPS + sPls 12; two images per group); two-sided Mann–Whitney U tests, Benjamini–Hochberg corrected (* q < 0.05, ** q < 0.01, *** q < 0.001; ns, not significant). **J,** Summary of **A–I** (astrocyte shape was not measured in culture).

Mitochondria were fragmented in KO tissue. VDAC1-labelled mitochondria formed continuous perinuclear networks in WT cells but short, separate rods and puncta in KO cells (**Fig. 8E**): mitochondrial length was 12% lower, branch junctions per object 32% lower and the share of mitochondrial area in networks 23% lower (**Fig. 8H**; every WT field above both KO fields, P = 0.067, the minimum attainable with 4 versus 2 fields).

Inflammation produced the same mitochondrial phenotype, and plasmalogen counteracted it. In cultured A1 astrocytes, LPS raised the punctate share of mitochondrial area from 3.0% to 7.0% (q < 0.01) and shortened network branches from 1.50 to 1.25 µm (q < 0.001); with scallop-derived plasmalogen (sPls) the punctate share fell to 4.3% and branches lengthened to 1.35 µm (each q < 0.05), and network junctions per cell rose from 13 to 42 (q < 0.001; LPS alone did not change them significantly) (**Fig. 8F,G,I**). Plasmalogen deficiency is therefore sufficient to fragment mitochondria in young brain tissue, and plasmalogen supplementation protects astrocyte mitochondria from inflammatory fragmentation. Because inflammation itself lowers glial plasmalogens^10^, the two routes may converge on mitochondria (**Fig. 8J**; **Fig. 1B**).

## Discussion

Across species, tissues and cells, the decline of plasmalogen-synthesis gene expression emerges as a feature of mammalian ageing with three properties, and our experiments add a fourth (**Fig. 1B**). It is conserved: the committed enzymes FAR2, GNPAT and AGPS lose expression in human, macaque and mouse tissues under every composition control, in rat in direction, in an independent human lifespan resource and within purified and directly converted human cells, with matching falls in macaque plasmalogen lipids and FAR1 protein. It is spatially pervasive: within ageing mouse brain, kidney, heart and spleen it is present in every niche rather than in a few vulnerable regions, it replicates in three independent brain cohorts and it is not explained by cellularity. In the brain it is glial: it occurs within oligodendrocytes, OPCs, microglia and astrocytes, where the committed enzymes lose expression, but not within neurons. And its loss matters for mitochondria: plasmalogen deficiency fragments brain mitochondria in young *Gnpat*-knockout mice, and plasmalogen supplementation rescues the mitochondrial networks of inflamed astrocytes. These properties fit the falling serum plasmalogens of older people^14,15,31^ and the early plasmalogen deficiency of Alzheimer’s and Parkinson’s disease^16–18^.

The decline is mammalian. It is absent or reversed in worm, fly, zebrafish and mouse retina, each with a plausible reason: the worm has no immune cells to repress GNPAT, and ether lipids are required for worm longevity; fly enzymes are too lowly expressed to score in most cell types; zebrafish brain regenerates throughout life and carries a duplicated GNPAT; and plasmalogen-rich photoreceptors are renewed daily from local synthesis. Within the mammals, the decline is clearest in tissue and in mouse glia but not in adult human cortical or blood cell types, so part of the bulk brain signal may reflect shifts between and within cell states that pseudobulk does not resolve.

The glial result connects the decline to brain inflammation. Oligodendrocytes and their precursors carry much of the brain’s plasmalogen, which is enriched in myelin^1,3^, and they are the cells in which the committed enzymes fall most; the same cells, together with microglia, raise complement, MHC and cytokine transcripts with age, and microglia shift from homeostatic to disease-associated and interferon-responsive states. This matches the white-matter focus of inflammatory brain ageing^21,22^. Yet the plasmalogen-synthesis decline is broader than inflammation in tissue — present in cortex, hippocampus, thalamus, striatum and white matter alike — and persists under composition control and within purified and converted human cells, so it is not a by-product of immune infiltration. Kidney is the peripheral tissue where the two meet: there the rate-limiting enzymes fall, an immune-infiltrate niche expands and low-plasmalogen, high-cytokine spots form contiguous territories. Glia and kidney are therefore the most promising settings for testing whether loss of plasmalogen synthesis permits local inflammation.

Mitochondrial gene-set transcripts did not fall within mouse glia, yet plasmalogen-deficient brain tissue showed fragmented mitochondria, which suggests that plasmalogen loss alters mitochondrial structure before mitochondrial gene expression. Plasmalogen deficiency impairs respiration and mitochondrial fission^7–9;^ in our experiments it fragmented brain mitochondria in young mice, and plasmalogen supplementation rescued the fragmentation that inflammation caused in astrocytes. Its relation to inflammation is reciprocal and less direct: inflammation represses GNPAT^10^ and plasmalogens restrain microglial NF-κB and TLR4 signalling^11,12,19,32^, yet *Gnpat*-knockout mice lack the innate-immune activation of peroxisome-deficient mice^33^. The timing of the decline relative to mitochondrial and inflammatory programmes and epigenetic ageing, and its response to interventions, are analysed in a companion study. Together, these results make plasmalogen replacement^19,32^ or targeted activation of the committed enzymes a candidate route to brain mitochondria, and the rescue of astrocyte mitochondrial networks by scallop plasmalogen (**Fig. 8F,G,I**) shows that this route can reach mitochondria directly.

This study has limitations. Except in macaque, the plasmalogen findings rest on transcripts, which do not measure enzyme activity or lipid content, the mouse brain lipidome does not fall by middle age, and plasmalogen levels were not measured in any immune population. The replication in purified and converted human cells is one of direction, with few donors per unit. Spatial spots mix cells, the spatial discovery cohort is male, and the spatial decline is relative to genes of similar expression. The brain cell-type analyses are secondary and were not pre-registered; they rest on few mice per age (4–8), pool 18- and 24-month animals, and disagree for neurons. The *Gnpat*-knockout experiment models lifelong, congenital plasmalogen deficiency in young male mice rather than gradual loss in old animals; its tissue mitochondrial result rests on 4 versus 2 imaging fields, the per-astrocyte morphology differences are nominal, GFAP exposure differed between genotypes, the cell type of the VDAC1-imaged cells was not identified, and plasmalogen levels were not measured in the LPS-treated astrocytes.

Further limitations of design are set out in Supplementary Note 2. The decisive experiments are lipidomic time series of the same tissues, conditional or adult-onset perturbation of FAR1, GNPAT or AGPS in glia with mitochondrial and inflammatory read-outs in more animals, and plasmalogen supplementation in aged mice.

## Methods

### Study design and data

Apart from the *Gnpat*-knockout and astrocyte experiments described below, this study re-analyses public data; all human data were de-identified at source. The public data comprise: (i) a three-mammal bulk atlas of 105 tissue age models (human GTEx v10^23^, rhesus macaque 30-tissue atlas^24^, mouse GSE132040^25^); (ii) single-cell ageing atlases of mouse^27,34^, human prefrontal cortex^35^ and blood^36^, macaque brain^37^, rat^38^, worm^39,40^, fly^41,42^ and zebrafish and mouse retina^43^; (iii) an independent human lifespan resource of eight datasets^23,44–50;^ (iv) spatial transcriptomes (GSE283283, GSE233983^22^, GSE212903^21^, GSE193107^51^); and (v) macaque metabolomes and proteomes from the same animals^24^ and a mouse brain lipidome^52^ (Supplementary Fig. 1). Raw downloads were verified and deleted after all features had been extracted.

### Data processing and quality control

Bulk libraries with fewer than 1,000 reads or zero total counts were removed before normalisation. Single-cell data were pseudobulked per donor or animal and cell type, requiring at least 5 cells per unit for the brain cell-type analyses; cells were never treated as replicates.

### Programme definitions

Plasmalogen synthesis was measured in three nested ways: the Reactome wax and plasmalogen biosynthesis pathway^53^ (bulk atlas); the committed enzymes FAR1, FAR2, GNPAT, AGPS and PEDS1/TMEM189 (pseudobulk landscape); and a 16-gene focus set (committed enzymes plus PEX5, PEX7, SELENOI, CEPT1, CHPT1, PCYT2, PLA2G6, LPCAT1, DHRS7B and ABCD1; single-cell, spatial). Macaque PEDS1 was excluded because its one-to-one ortholog resolves to the UBE2V1 read-through locus. Peroxisomal import (Reactome peroxisomal protein import, including PEX5 and PEX7) was scored separately. In the worm, the ether-lipid pathway comprised *fard-1*, *acl-7* and *ads-1*. Mitochondrial programmes were Hallmark OXPHOS and Reactome mitochondrial translation; inflammation was Hallmark inflammatory response, interferon-α/γ response and TNF-α/NF-κB, with complement, cytokine/chemokine and MHC sets in the spatial and brain cell-type analyses^53–55^.

### Age models and gene-set scoring

Bulk counts were normalised with DESeq2 size factors^56^, log-transformed and modelled by least squares on age as a fraction of maximum lifespan (human 122.5, macaque 40, mouse 4 years) with available covariates (GTEx: sex, RNA integrity, ischaemic time, and circumstances of death in a sensitivity analysis). Gene-level age statistics were converted to signed z values, detrended on mean abundance (LOWESS), and gene sets were scored against random sets matched for the set’s abundance-decile profile (analytic null for the bulk atlas; 2,000 draws elsewhere). Sample-level scores were the set’s mean standardised expression minus the expectation for expression-decile-matched random sets. The detrending bandwidth is not load-bearing (frac = 0.1, 0.3 or 0.6 and a decile-mean alternative change no signs).

Because a single continuous predictor explains about 1/(n − 1) of a gene’s variance by chance, variance partitioning was judged against that sample-size null. Power in the small macaque cohorts was tested by injecting human- or mouse-sized age effects into each cohort’s own data (dispersion, library sizes, covariates and n).

### Composition control

Cell-class proportions were estimated per sample by non-negative least squares against a Tabula Muris Senis reference (11 classes, 440 markers)^27^, and every age model was refitted with centred-log-ratio myeloid and lymphoid fractions and, additionally, the first principal component of the other classes (Supplementary Figs. 6 and 7). Brain cohorts were also refitted against a primate multi-region brain reference (9 classes, 360 markers)^37^. This per-sample deconvolution superseded a between-cohort adjustment, in which, for key terms, cohort z was regressed on each cohort’s immune-marker shift across the 105 cohorts and the median value at zero shift is reported (Supplementary Data 1, sheet S10_composition).

### Pre-specified arms beyond the bulk atlas

Rat (tissue × cell class, old versus young, 2 versus 2 libraries), zebrafish, fly, mouse retina and worm single-cell cohorts, human prefrontal cortex and the Allen mouse brain atlas were analysed under plans fixed before the data were examined, and every cohort is reported whatever its outcome (Fig. 1D; Supplementary Data 3). Human prefrontal cortex (PsychAD) was analysed as donor × subclass pseudobulk with edgeR with age, sex and brain bank: 192 adults aged 20–89 years (primary), the same without the nine donors top-coded at 80 years and over (sensitivity), and 57 donors under 20 years (development, separate). The Allen mouse brain atlas (64 adult, about 2 months, and 44 aged, 18 months, mice; donor × subclass) was analysed with a region-adjusted fit on unbiased-sampled cells (brain division as covariate) and a class-level fit. Human blood cell types were taken from OneK1K (981 donors aged 19–97 years).

### Single-cell visualisation

Single-cell embeddings (Harmony^57^ and UMAP^58^; Supplementary Figs. 2 and 3) and per-cell scores of the committed enzymes (scanpy^59^ score_genes) are shown for illustration; every age comparison averages scores per animal or donor and cell label and compares old with young by Welch’s t-test (Fig. 2C–E).

### Human lifespan resource

Eight public bulk RNA-seq datasets (4,513 samples) were combined: brain tissue (BrainSeq DLPFC, 320 controls, prenatal to 97 years^47^; BrainSpan, 16 regions, prenatal to 40 years^48^; post-mortem prefrontal cortex^50^; cortical tissue^46^); purified brain cells (HepaCAM-purified astrocytes, fetal to 65 years^45^; CD11b-purified cortical microglia, 34–102 years^46^); directly converted cells (induced neurons and their donor fibroblasts, 0–89 years^50^; induced astrocytes from young and old donors^49^), in which direct conversion bypasses pluripotency and keeps donor age^49,50^; cultured fibroblasts (a 133-donor lifespan series^44^); and GTEx v8 (13 brain regions and cultured fibroblasts^23^). Each dataset × cell type was a unit (25 units). Counts were modelled with limma-voom with sex, platform or batch, region, RNA integrity, ischaemic time and Hardy scale where available; contrasts were the adult age trend per decade (adults 20–59 and older 60+), older versus adult, and developmental contrasts (prenatal, newborn 0–2, young 2–19). Gene sets were tested by pre-ranked GSEA (fgsea) on the moderated t, with Benjamini–Hochberg correction within unit (Supplementary Fig. 12). Per-sample module scores were the mean z of member genes across samples of a unit and were related to donor age in adults (≥ 20 years) by Spearman correlation with a LOWESS line for display (Fig. 2H–K; Supplementary Figs. 14 and 15); with prenatal and child samples included, the astrocyte score appears to rise (ρ = +0.41), driven by the six prenatal samples.

### Spatial transcriptomics

#### Design and pre-registration

The spatial arm was planned after download and extraction but before any gene-set scoring or age test; the plan was frozen by recording its SHA-256 checksum, and three amendments, all made before any age test, are logged with their reasons (brain abundance computed per section and averaged per coronal level; a two-level Stereo-seq annotation rule; an RCTD-style platform correction for deconvolution with a marker-score cross-check). The biological unit is the mouse throughout: spots and sections are never replicates. Hypotheses H1–H3 predicted that plasmalogen synthesis (16-gene focus set) and OXPHOS subunits decline and the inflammatory response rises; support in a tissue required at least half of its populations in the expected direction, at least one population at FDR < 0.05 and a tissue-level slope in the expected direction, with same-sign replication in GSE233983 for brain claims. H4 concerned abundance changes and H5 the relation between overall transcriptional ageing and the programmes; the pre-registered coupling and attenuation hypotheses (H6, H7) are reported in a companion study.

#### *Data.* Discovery

GSE283283 Visium ageing cohort (male C57BL/6JN; 4, 17 and 26 months; brain at five coronal levels, heart, kidney, liver, lung and spleen; 10–14 mice per tissue). Replication (brain): GSE233983 (2.5, 13 and 29 months, both sexes, 24 mice, 118,573 spots; resequencing libraries summed into their sections)^22^; GSE212903 (6, 18 and 21 months; six sections)^21^; GSE193107 (2 versus 28 months; eight sections; age confounded with slide)^51^. Single-cell resolution: GSE283283 Stereo-seq cellbin (one mouse per age, five sections each; 658,243 cells, 60% assigned a type; descriptive). The GSE283283 infection cohort was excluded as pre-registered.

#### Processing

Spots with ≥ 500 UMIs and ≥ 250 genes were kept, excluding spots above 30% mitochondrial UMIs (50% in heart); sections with < 300 spots were dropped; genes detected in ≥ 0.5% of spots were kept. Counts were normalised to 10,000 per spot and log-transformed; pseudobulk counts were summed per mouse and population and expressed as log₂ CPM. Niches: 3,000 highly variable genes, 50 principal components, Harmony on section^57^, k-nearest-neighbour graph (k = 15), Leiden clustering (resolution 0.6), giving 57 niches (19 brain, 5 heart, 9 kidney, 7 liver, 9 lung, 8 spleen) labelled from marker genes without reference to age (Fig. 3A,B; Supplementary Figs. 16 and 17). Marker genes were ranked by a Wilcoxon test of each niche against all other spots of the tissue (brain, random subsample of 60,000 spots); the top 30 genes with positive fold change per niche and the label of every niche are given in Supplementary Data 5 (sheets niche_markers and niche_labels). Cell-type proportions were estimated per spot by corrected non-negative least squares (RCTD-style^60^) against Tabula Muris Senis (peripheral tissues) or broad Stereo-seq^61^ classes (brain) (Supplementary Fig. 19). The deconvolution does not resolve brain neuronal subtypes or lung, which are therefore interpreted only through niches and marker scores.

#### Statistics

Programme scores per spot: scanpy score_genes (set mean minus 50 expression-matched control genes per gene, 25 bins), averaged per mouse and population. Age models per mouse: y ∼ age (+ sex in GSE233983), OLS with HC3 errors; Benjamini–Hochberg within tissue over populations and the three primary sets. Replication niches were transferred from the discovery brain by centroid correlation on shared highly variable genes; meta-analysis used inverse-variance weighting of standardised slopes across the four datasets. GSE212903 and GSE193107 have no mouse identifiers, so their sections were treated as animals.

#### Images (pre-registered Addenda A and B)

Written before any image-derived measure was related to age, niche or transcriptome, the addenda specified that nuclei be segmented in the full-resolution H&E images and assigned to spots; sections failing a registration check were dropped (all spleen sections) (segmentation quality control in Supplementary Fig. 31). Cellularity was modelled as score ∼ age + log nuclei + niche + (1 | mouse) at spot level, and by mouse-level age models with and without mean nuclei per spot (Fig. 4E).

### Brain cell types and secondary spatial analyses

These analyses were added after the pre-registered spatial analysis was complete; they are secondary and descriptive, were not pre-registered, and are reported in full whatever their outcome. Tabula Muris Senis FACS brain cells (Smart-seq2; myeloid and non-myeloid sorts)^27^ were log-normalised and scored per cell with the spatial gene sets (scanpy score_genes, 50 control genes); scores were averaged per mouse and cell type (≥ 5 cells), and 3-month mice were compared with 18–24-month mice by Welch’s t-test with Hedges’ g and Benjamini–Hochberg correction within programme; 18- and 24-month mice were pooled because 24-month non-myeloid cells are few. Gene-level changes used pseudobulk counts per mouse and cell type (log₂ CPM; genes with mean young log₂ CPM < 2 not tested). Microglial states (homeostatic, proliferating, IFN-responsive, DAM/ARM, MHC-II-high, monocyte-derived, border-associated macrophages, dendritic cells, neutrophils) were assigned from marker scores with thresholds fixed before age was examined.

In the spatial discovery cohort, the 3.9-month and 25.8-month mice (3–5 per age) were compared per tissue and per mouse (Welch t-test, Hedges’ g, Benjamini–Hochberg within tissue) for the plasmalogen-synthesis focus set and seven mitochondrial and seven inflammatory gene sets (Fig. 6A). Plasmalogen-synthesis (blue) and cytokine (orange) scores were drawn as smoothed landscapes, 3 × 3 bivariate maps and niche-change maps for the four tissues where the plasmalogen-synthesis score falls, with every section image paired with a per-mouse quantification (Fig. 3; Supplementary Figs. 21M–R, 22 and 23). For tissue territories, cell-type signatures (genes ≥ 2.7-fold higher in one Tabula Muris Senis brain cell type than in every other, detected in ≥ 30% of its cells; up to 30 genes) were scored on Visium brain (GSE283283, 4 versus 4 mice; GSE233983, 10 versus 10). A glial index (mean z of oligodendrocyte, astrocyte and microglia signatures minus the neuron signature) defined glia-rich (top quartile) and neuron-rich (bottom quartile) territories per section (Fig. 6C; Supplementary Fig. 41). For exploratory communication analyses, ligand–receptor pairs from the OmniPath intercell network were scored per mouse as √(mean ligand in sender × mean receptor in receiver) in Tabula Muris Senis brain cells and, on Visium, as ligand(spot) × mean receptor of the spot and its six neighbours; neuronal synaptic and activity-gene scores were regressed per section on the plasmalogen-synthesis score of the surrounding 100–300 µm ring (Supplementary Figs. 42–44).

### Lipids and proteins

The 17 macaques of the transcriptome atlas^24^ were profiled by untargeted metabolomics and proteomics in up to 30 tissues. The ether-PE fraction was the mean log₂ ether-linked minus diacyl PE per animal and tissue; the ether-PE fraction, FAR1 protein, an OXPHOS protein score and an inflammatory/complement protein score were regressed on age per tissue, and the share of tissues in the hypothesised direction was tested by an exact sign test. The mouse brain lipidome (ten regions^52^) was analysed per region with linear models of age and sex, from 3 to 16 weeks (development) and from 16 to 59 weeks within one batch (adult; 8 male and 8 female mice per age), with Benjamini–Hochberg correction across regions; the ether-PC fraction was analysed in the same way (Supplementary Data 3).

### Gnpat-knockout mice and astrocyte experiments

Heterozygous *Gnpat*-knockout mice (*Gnpat*^+/−^) on a C57BL/6J background were generated and supplied by Cyagen Biosciences (contract GEM-KOAIP-230630-CAL-03; knockout line EK5264, founder line 2-1766) and intercrossed to obtain wild-type (*Gnpat*^+/+^) and knockout (*Gnpat*^−/−^) offspring; the same colony was used for our analysis of the *Gnpat*-knockout testis^62^. Male WT and KO mice [littermates: confirm; number per genotype] were analysed at 14 weeks of age [genotyping method; housing conditions]. All animal experiments were approved by the Ethics Committee for Animal Experiments of the Institute of Rheological Functions of Food [approval number], were carried out in accordance with the institutional guidelines and are reported in line with the ARRIVE guidelines. Brains were fixed and sectioned [fixation, section thickness, brain region], im-munostained for GFAP or VDAC1 [antibodies, dilutions] with DAPI, and imaged on a Keyence BZ-X microscope (full-focus images, 0.8 µm per pixel). GFAP-positive astrocytes were segmented and their processes traced from the soma; 41 per-cell morphology descriptors (618 WT and 651 KO astrocytes) were compared with a linear mixed model with image as random intercept, and field-level measures (astrocyte density, GFAP-positive area, nuclear density; 6 versus 6 images) with Welch’s t-test. Mitochondria in perinuclear VDAC1 fields were segmented and classified as punctate, rod or network/branched, and length, branch junctions per object and network area were compared between WT (4) and KO (2) fields by an exact two-sided permutation test, for which 0.067 is the smallest attainable P. A1 astrocytes [cell source; induction] were treated with LPS [concentration, duration] with or without scallop-derived plasmalogen (sPls) [source, concentration, timing], stained with MitoTracker Orange and DAPI, and mitochondria were classified as above (control 9, LPS 30, LPS + sPls 12 cells from two images per group); the punctate share of mitochondrial area, mean network branch length and network junctions per cell were compared by two-sided Mann–Whitney U tests with Benjamini–Hochberg correction. Image analysis used custom Python code (BZX_image_analysis).

### Statistics and reproducibility

Tests were two-sided unless stated; agreement across cohorts was tested with exact binomial sign tests rather than by the significance of single correlations; multiple testing was controlled by the Benjamini–Hochberg procedure^63^. Random-set nulls used fixed seeds. Analyses marked as pre-specified followed plans fixed before the data were examined, and every cohort is reported whatever its outcome. Every main and supplementary figure passed an automated text-overlap test, and an automated dry test checked that every figure exists, is newer than its script and renders, that source tables have values in range and correct multiple-testing relations, that every Supplementary Figure is cited in the main text and that every citation resolves. Main figures were assembled from panels of these figures with a table mapping each panel letter to its source figure.

## Data availability

All public data analysed are listed here. Human: GTEx v10 and v8, BrainSeq Phase 1, BrainSpan, GEO GSE113957, GSE73721 and GSE99074, ArrayExpress E-MTAB-3037, OneK1K, and the PsychAD Aging cohort (CELLxGENE). Macaque: figshare 10.6084/m9.figshare.26963386 and GEO GSE307184. Mouse: GEO GSE132040, GSE283283, GSE233983, GSE212903, GSE193107 and GSE325479, Tabula Muris Senis, the Allen Brain Cell Atlas and Metabolomics Workbench ST001637/ST001888. Rat: GEO GSE137869. Zebrafish: GSE325479, GSE197673, GSE205601. Fly: GSE218661, GSE107451. *C. elegans*: GSE229022, GSE208154. Source data for Figs. 1–7 are provided as Supplementary Data 1–5. The cohort list, the spatial quality-control summary and the human lifespan unit definitions, which a companion study also uses, are deposited at Zenodo [DOI]. The *Gnpat*-knockout and astrocyte images and per-cell measurements underlying Fig. 8 are available from the corresponding authors [or: deposited at repository, accession].

## Code availability

All code used for data analysis can be provided upon reasonable request to the corresponding authors.

## Author contributions

N.S. designed the experiments, performed the analyses and wrote the manuscript. M.H., S.M. and T.F. helped design the experiments and revised the manuscript.

## Supplementary Information

Supplementary Notes 1 and 2 follow, then Supplementary Figures 1–44, each with its legend. Supplementary Data 1–5 (Excel workbooks) are provided as separate files.

## Supplementary Notes

### Supplementary Note 1. Additional analyses and controls

These passages extend the Results of the main text (moved here for length).

#### Analysis controls

Several controls shaped these comparisons (Methods). Detrending on mean expression attenuates but does not remove the abundance artefact (the product of the two datasets’ trends still predicts the detrended cross-species correlation, ρ = +0.384, p = 0.012), so gene-level correlations are upper bounds and cross-species conservation is mapped at pathway resolution (Supplementary Figs. 4 and 5). Significance is not agreement (of 1,365 worm × mammal gene-level pairings, 503 pass FDR < 0.05 at a median correlation of +0.009), so agreement is tested by sign tests over cohorts. A flat macaque signal is not a lack of power: injected human- or mouse-sized age effects are recovered in five of the six macaque cohorts that returned almost none, with a false-positive floor of zero; only macaque heart is genuinely underpowered. As a positive control, canonical tissue markers fall in the top 5% of tissue specificity (τ), which is conserved across species (ρ = +0.69 to +0.82) and does not confound age association (median ρ = +0.041). Pooling tissues can reverse an age effect (CDKN2B appears to fall across all human samples, ρ = −0.09, but rises within 9 of 20 tissue concepts), so age effects are estimated within tissue. Against a sample-size null, the donor explains more variance than age for 96–100% of genes in every species.

#### Species outside the mammals

The rat result rests on 2 versus 2 libraries per cohort. In mouse and zebrafish retina, plasmalogen synthesis does not fall while OXPHOS falls and inflammation rises; zebrafish cohorts are dominated by retina, and in zebrafish whole brain inflammation rises while OXPHOS and plasmalogen synthesis are flat. In fly, plasmalogen synthesis falls only in trend, although a decline has been reported in ageing oenocytes. In the worm the ether-lipid transcripts (*fard-1*, *acl-7*, *ads-1*) rise with age, whereas worm ether-lipid proteins are reported to fall^3^.

#### Inflammation and development in the human lifespan resource

Inflammation in this resource is tissue-only, as the composition analysis predicts: inflammatory response and complement rise in 13 and 11 of 14 GTEx brain units but in 1 and 0 of 4 cell-intrinsic units, purified microglia being the one cell-intrinsic unit with a significant inflammatory rise. OXPHOS falls in GTEx brain (3 of 14 rising; 7 significant falls), in induced neurons (NES −3.2) and in the 133-donor fibroblast series (−2.0), but not in independent bulk brain or purified astrocytes (BrainSeq DLPFC +0.7, prefrontal cortex +1.8, astrocytes +2.0).

Development runs differently. From prenatal to adult life, OXPHOS rises in BrainSeq, BrainSpan and purified astrocytes (NES +2.3, +2.4 and +3.0), as do mitochondrial translation and the lysosome, whereas plasmalogen-enzyme transcripts split: *GNPAT* (t = −15.9 and −6.5) and *AGPS* (−10.2 and −3.5) fall in BrainSeq and BrainSpan whereas *FAR1*, *FAR2* and *PEX7* rise, and in purified astrocytes the pathway runs opposite in development and adult ageing (**Fig. 2G**), which is why the adult age trend is the primary contrast.

#### Mitochondrial programmes in the spatial cohort

The pre-registered mitochondrial hypothesis was supported in liver only (OXPHOS −0.83, q = 0.009). In the oldest-versus-youngest contrast, mitochondrial sets fell in liver (OXPHOS −2.24, mitochondrial translation −1.94, respiratory-chain assembly −2.15, TCA −3.10, fatty-acid oxidation −2.67; all p < 0.03) and in direction in lung and heart, whereas mitochondrial translation and ribosome sets rose in kidney (+1.6, +2.0). Across 68 populations, the fraction of age-associated genes correlates with the plasmalogen-synthesis slope (ρ = −0.34, p = 0.004, q = 0.13), a suggestive relation. How the programmes relate to one another spot by spot, between individuals and across tissues is analysed in a companion study.

#### Glia-to-neuron lipid supply (exploratory)

Exploratory analyses asked what transcripts can test of the hypothesis that glia supply lipids to neurons (Supplementary Fig. 42A), using the expression of synthesis, lipid-carrier and uptake-receptor genes by brain cell type (Supplementary Fig. 42B,C). No glia-to-neuron ligand–receptor pair passed q < 0.1 in single cells (Supplementary Fig. 43). In tissue, Spp1, Apoe, Il1a, Il1b and Csf1 pairs rose in GSE233983 (q < 0.05) and C1qa → Lrp1 in GSE283283 (q = 0.02) (Supplementary Fig. 44), and neighbourhood plasmalogen-synthesis scores did not predict neuronal synaptic and activity genes in an age-dependent way (Supplementary Fig. 42D–F). Transcripts cannot measure lipid transfer.

### Supplementary Note 2. Further limitations of design

Further limitations concern design. The composition control is partial: broad cell classes miss shifts within compartments^64^, adjustment may remove signal that is itself downstream of ageing, and the whole-body reference is murine and cross-mapped. No human brain single-cell reference was available, so the brain control uses macaque snRNA-seq cross-mapped by gene symbol, and marker-based estimates cannot distinguish neuron loss from reduced neuronal transcript output, although normal brain ageing largely preserves neuron numbers^65,66^. The independent human set is small (nine units) and heterogeneous; no deconvolution was applied to its bulk units, for which the cell-intrinsic units stand in; BrainSpan stops at 40 years and public GTEx ages are 10-year bins; and the induced-astrocyte data are a pre-normalised array table with three donors per group. In the spatial arm, brain neuronal subtypes are not resolved by deconvolution, two replication datasets lack mouse identifiers and one confounds age with slide (Methods), and some niches are partly defined by immune genes, so their inflammatory scores are partly circular. FACS sorting can alter transcripts and under-samples neurons, and ligand–receptor scores are co-expression, not signalling. No cohort observes the last third of its species’ lifespan: the oldest animals reach 0.61 (human), 0.68 (macaque), 0.56 (mouse) and 0.47 (worm) of maximum lifespan, so nothing here speaks to late life. The macaque atlas is female-only, so sex cannot be modelled there and human–macaque comparisons are not sex-balanced, and macaque batch 1 is exactly the three 14-year-old animals, so batch is partly confounded with age (the batch term is retained). Species is completely confounded with study, laboratory and platform, because no matched cross-species ageing design exists in public data; every cross-species claim therefore compares age coefficients estimated within species, never expression, and between single-cell-type worm cohorts and bulk mammalian tissues only directions are comparable. GTEx brain regions share donors, so brain cohorts are not independent tests. In the knockout, lower GFAP-positive density may reflect fewer astrocytes or less GFAP per astrocyte. Finally, these are transcriptomic signatures, not biomarkers validated against outcomes in the sense proposed by the Biomarkers of Aging Consortium^67,68^. The decisive experiments are lipidomic time series of the same tissues, conditional or adult-onset perturbation of FAR1, GNPAT or AGPS in glia with mitochondrial and inflammatory read-outs in more animals, and plasmalogen supplementation in aged mice.

## Supplementary Figures

**Supplementary Fig. 1.**
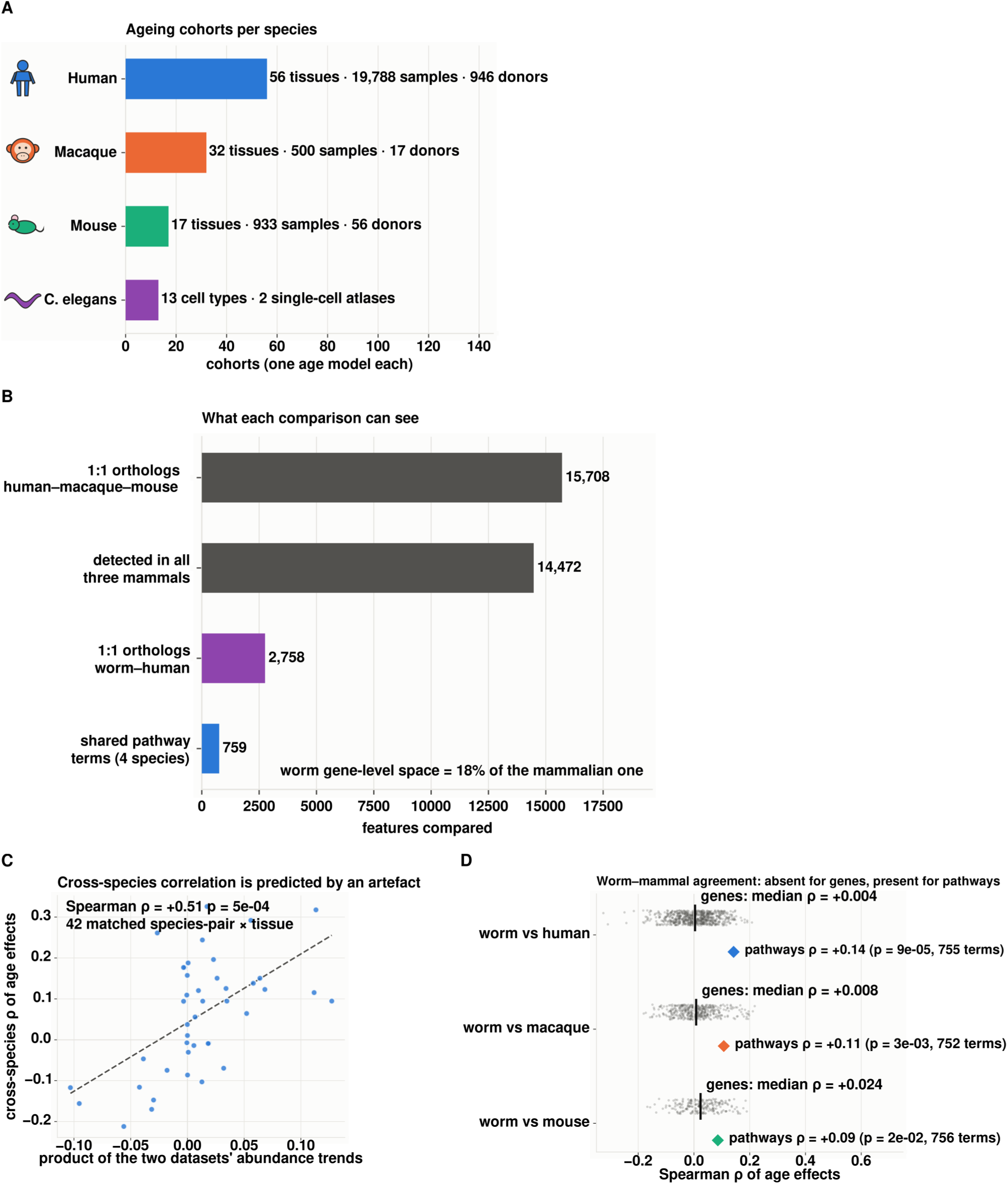
Resource and why comparisons are made at pathway resolution. **A,** Ageing cohorts per species (one age model each): human GTEx v10, 56 tissues (19,788 samples, 946 donors); rhesus macaque 30-tissue atlas, 32 tissues (500 samples, 17 female animals aged 3–27 years); mouse GSE132040, 17 tissues (933 samples, 56 animals); and *C. elegans*, 13 single-cell-type cohorts from two independent atlases (GSE229022, wild-type N2 single-nucleus, days 1–14; GSE208154, germline-less *gon-2* single-cell, days 1–11). Age is modelled as a fraction of each species’ maximum lifespan. **B,** Features available to each comparison: 15,708 one-to-one human–macaque–mouse orthologues, 14,472 of them detected in all three mammals; 2,758 worm–human one-to-one orthologues (18% of the mammalian set); and 759 pathway terms annotated in all four species’ own genes. **C,** Why gene-level correlations mislead. Each point is one of 42 matched species-pair × tissue comparisons. x, the product of the two datasets’ abundance trends (within each dataset, the Spearman ρ between per-gene age slope and mean expression); y, the cross-species Spearman ρ of per-gene age effects over orthologues. Dashed line, least-squares fit; Spearman ρ = +0.51, p = 5 × 10⁻⁴. **D,** Worm–mammal agreement. Grey points, gene-level Spearman ρ for each worm × mammal cohort pairing over one-to-one orthologues (1,365 pairings; black bar, median, +0.004 to +0.024). Diamonds, pathway-level Spearman ρ over 752–756 shared terms, with p values. Per-cohort statistics: Supplementary Data 1; the gene-level analyses are detailed in Methods.

**Supplementary Fig. 2.**
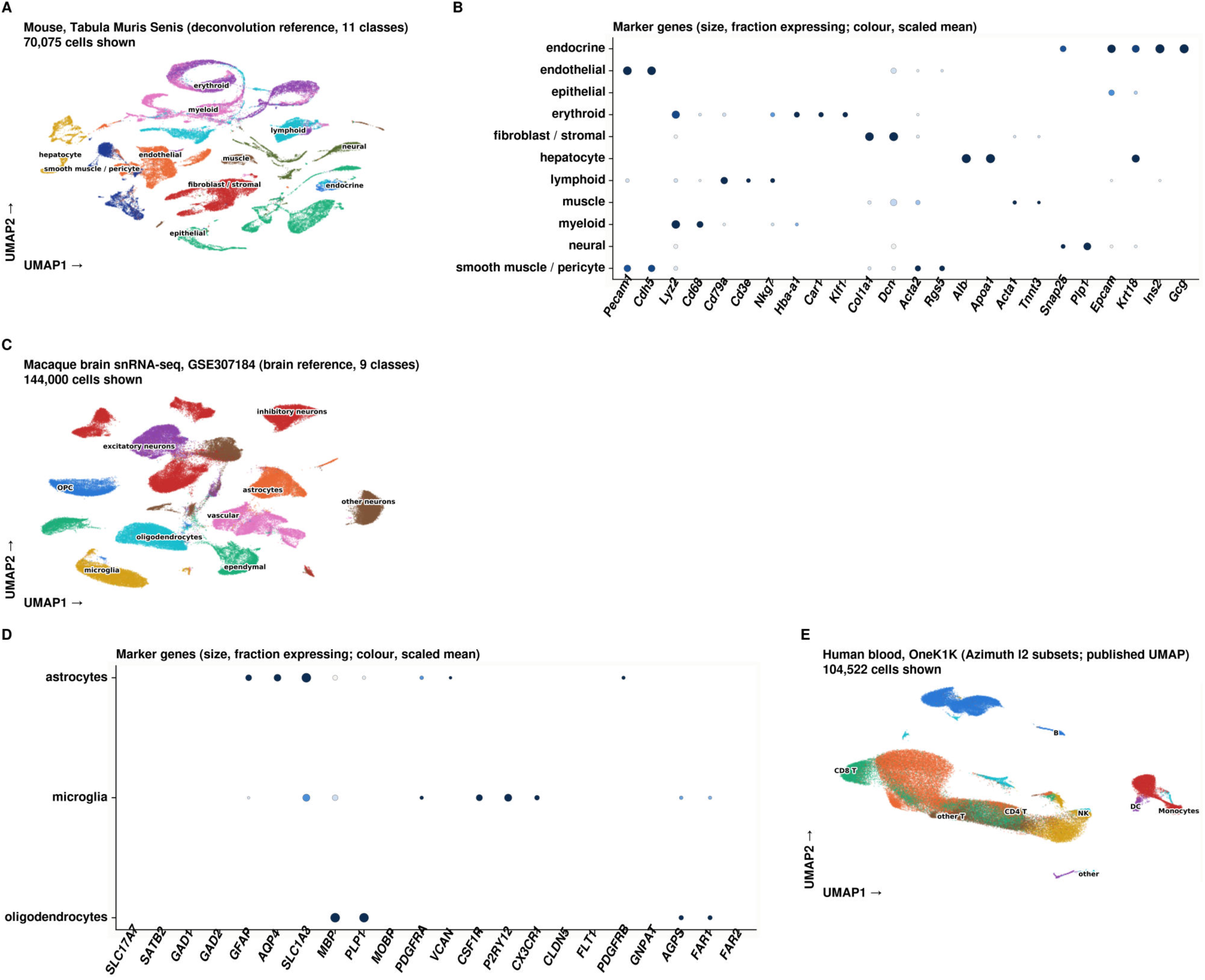
Single-cell datasets used in this study: cell identities and marker genes (part 1 of 2). UMAP (left; stratified subsample of up to 6,000–12,000 cells per label; Harmony-corrected; Methods) and marker-gene dot plot (right; dot size, fraction of cells expressing; colour, mean log expression scaled per gene). **A,B,** Tabula Muris Senis droplet and FACS, 70,075 cells: the 11 broad classes of the whole-body deconvolution reference. **C,D,** Macaque brain snRNA-seq (GSE307184), 144,000 nuclei: the 9 classes of the brain composition control (dot plot for the glial classes astrocytes, microglia and oligodendrocytes). **E,** Human blood (OneK1K), 104,522 cells: published UMAP coloured by lineage (dot plot in Supplementary Fig. 3A).

**Supplementary Fig. 3.**
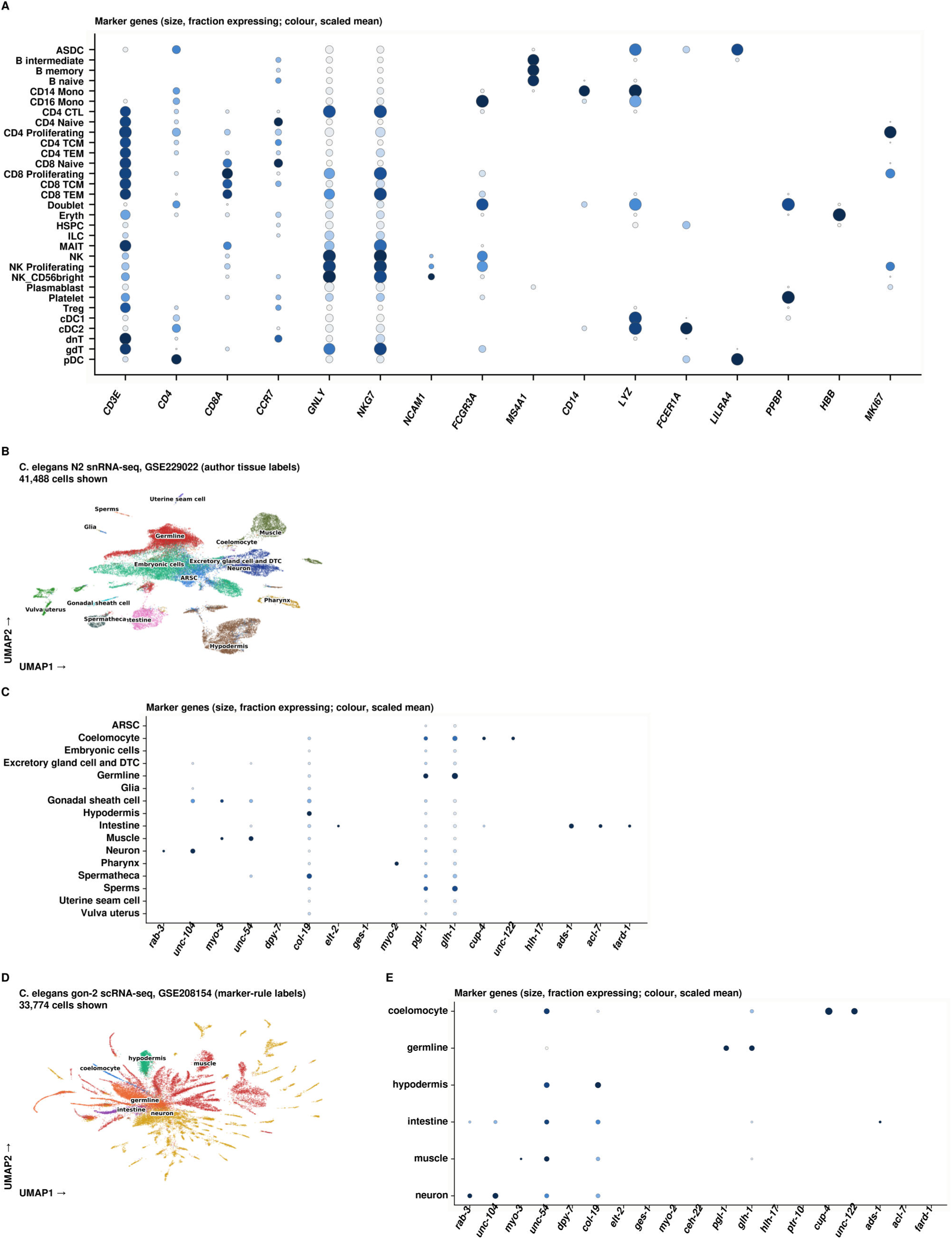
Single-cell datasets used in this study: cell identities and marker genes (part 2 of 2). Continued from Supplementary Fig. 2. **A,** Human blood (OneK1K): marker-gene dot plot at the 31 Azimuth l2 subsets. **B,C,** *C. elegans* N2 snRNA-seq (GSE229022), 41,488 nuclei from the 12 N2 samples analysed, author tissue labels. **D,E,** *C. elegans* gon-2 scRNA-seq (GSE208154), 33,774 of 77,128 cells, labels from the highest of eight marker-set scores per Leiden cluster.

**Supplementary Fig. 4.**
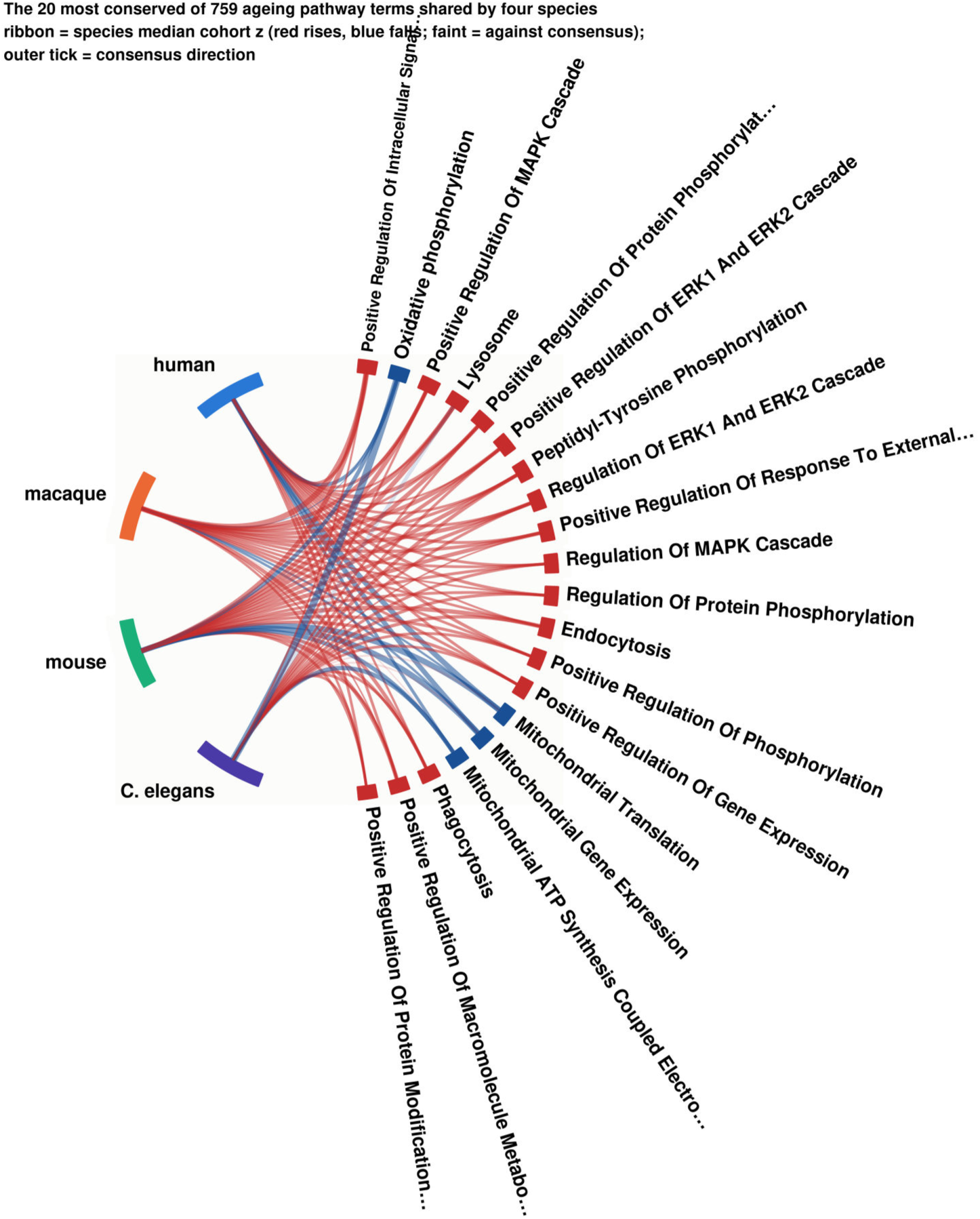
Cross-species conservation map (part 1 of 2). Chord diagram linking four species (left) with the 20 pathway terms of highest conservation score (right); ribbon colour = direction of the species’ median cohort z (red rises, blue falls), width = |z|, faint ribbons disagree with the consensus direction (outer tick).

**Supplementary Fig. 5.**
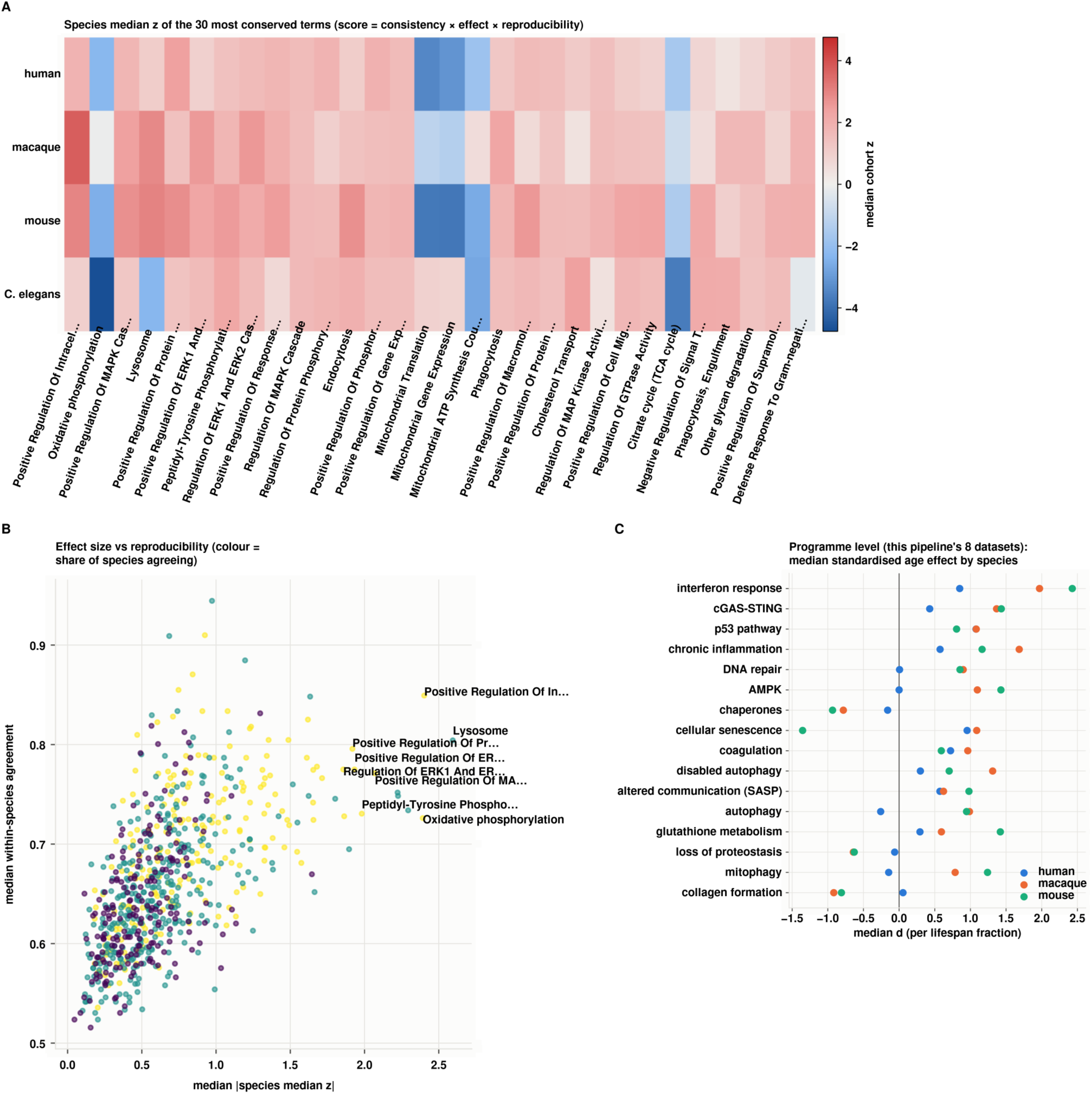
Cross-species conservation map (part 2 of 2). Continued from Supplementary Fig. 4. **A,** Species median z for the 30 most conserved terms. **B,** Effect size against reproducibility for all shared terms (colour = share of species agreeing). **C,** Programme-level effects from this pipeline’s eight datasets: median standardised age effect *d* per species.

**Supplementary Fig. 6.**
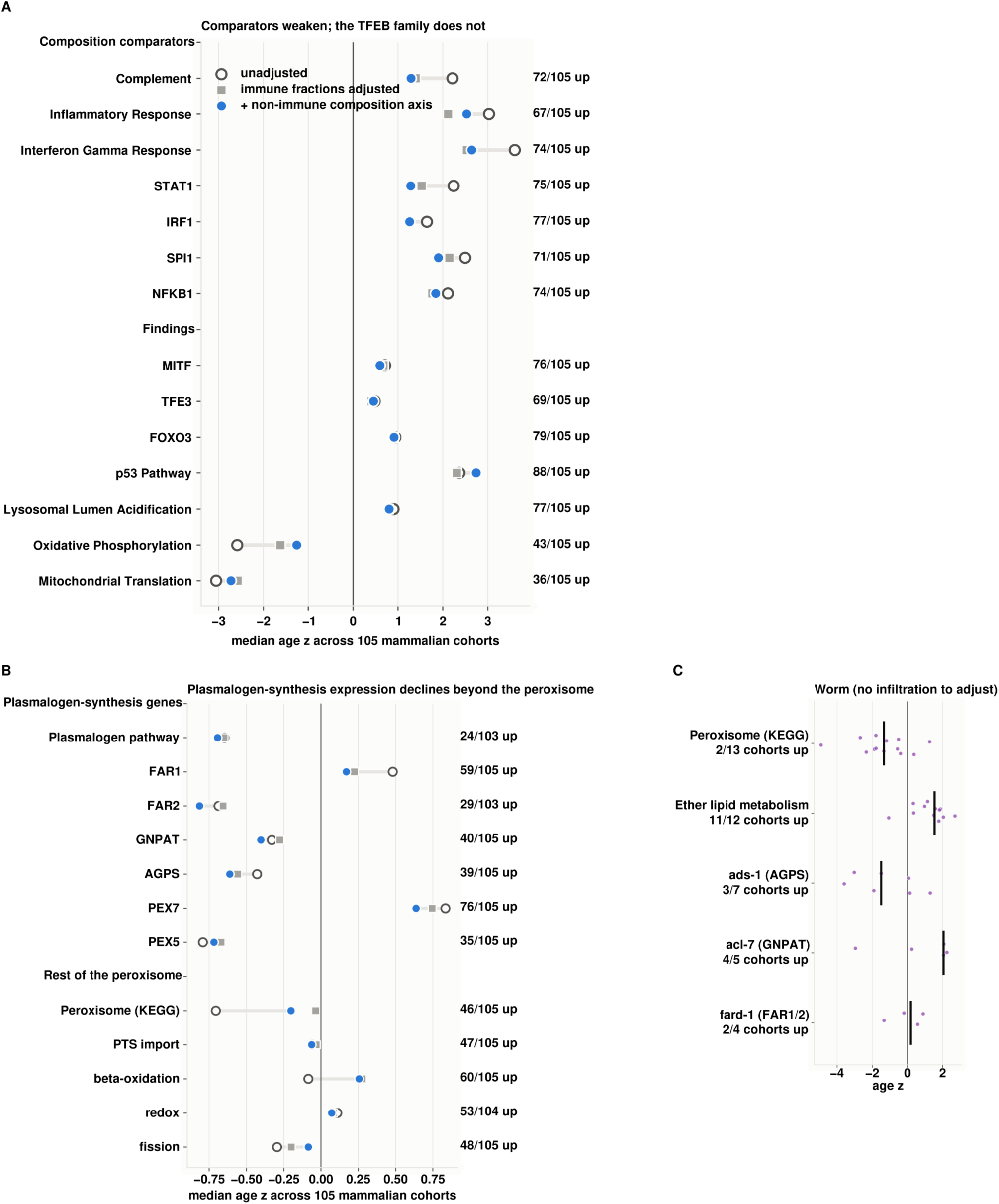
Plasmalogen-synthesis expression declines beyond cell composition (part 1 of 2). Cell-class proportions were estimated for each of 21,221 mammalian samples by non-negative least squares against a Tabula Muris Senis reference (11 broad classes, 440 marker genes; no plasmalogen enzyme among the markers). Every age model was refitted with the centred-log-ratio myeloid and lymphoid fractions (immune-adjusted) and additionally with the first principal component of the other classes (fully adjusted). Values are median age z across cohorts, each gene set scored against 2,000 abundance-matched random sets. **A,** Composition comparators (complement, inflammatory and interferon-γ responses, STAT1, IRF1, SPI1, NF-κB1) and other findings (TFEB-family and FOXO3 regulon output, p53 pathway, lysosomal acidification, OXPHOS, mitochondrial translation) across 105 cohorts: open circles, unadjusted; grey squares, immune-adjusted; blue circles, fully adjusted. Text, cohorts rising after full adjustment. **B,** The same for plasmalogen-synthesis genes (Reactome wax and plasmalogen biosynthesis; *FAR1*, *FAR2*, *GNPAT*, *AGPS*; *PEX7* and *PEX5*) and for the rest of the peroxisome (KEGG peroxisome; PTS import, β-oxidation, redox and fission modules); 103–105 cohorts with enough expressed genes. **C,** *C. elegans*, whose single-cell-type cohorts need no composition adjustment: KEGG peroxisome, KEGG ether-lipid metabolism and the plasmalogen-enzyme orthologues *ads-1* (AGPS), *acl-7* (GNPAT) and *fard-1* (FAR1/2). Points, cohorts; bar, median; text, cohorts rising.

**Supplementary Fig. 7.**
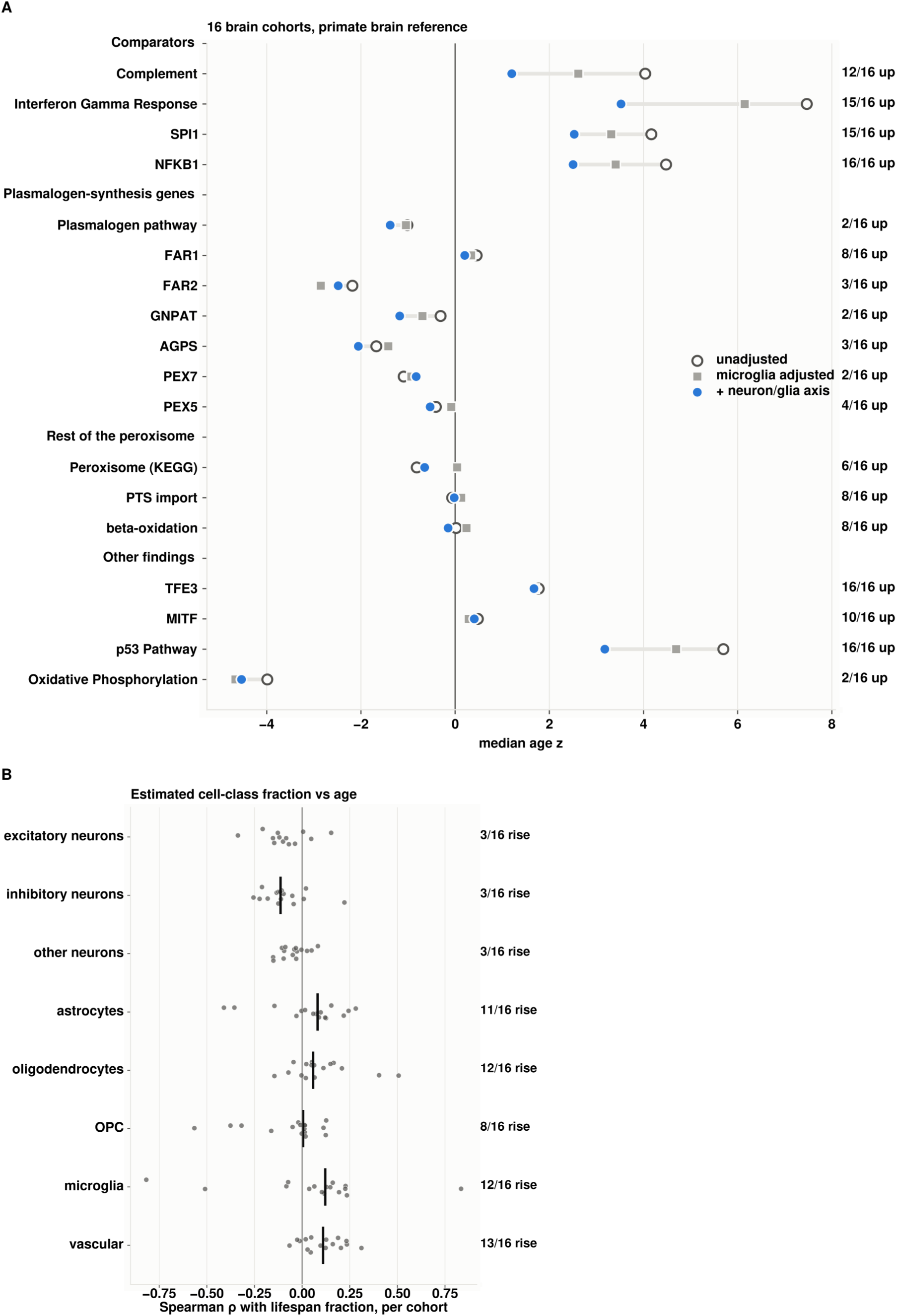
Plasmalogen-synthesis expression declines beyond cell composition (part 2 of 2). Continued from Supplementary Fig. 6. **A,** The 16 mammalian brain cohorts refitted against a primate brain reference (macaque multi-region snRNA-seq, GSE307184; 55 donors, 9 classes, 360 markers): open, unadjusted; grey, adjusted for the centred-log-ratio microglial fraction; blue, additionally for the first principal component of the other classes (neuron–glia axis). Text, brain cohorts rising after full adjustment. **B,** Spearman ρ between each estimated brain cell-class fraction and lifespan fraction within each brain cohort; points, cohorts; bar, median; text, cohorts in which the fraction rises. Per-cohort values: Supplementary Data 1 (sheets S13–S15).

**Supplementary Fig. 8.**
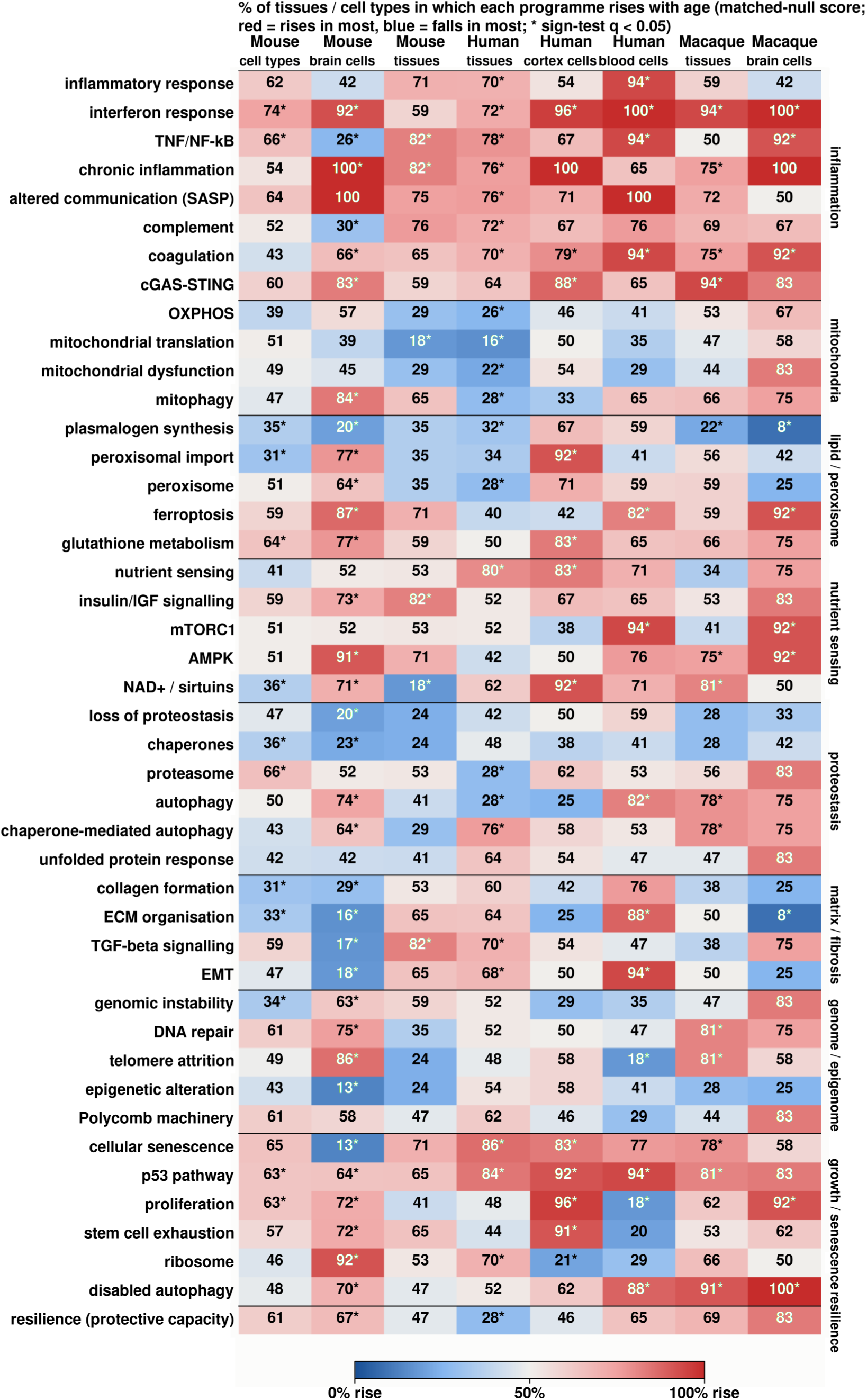
Programme atlas with matched-null scoring (part 1 of 3). For 44 programmes (rows, grouped by theme) in eight datasets (columns), the percentage of tissues or cell types in which the programme’s matched-null score rises with age (red = rises in most, blue = falls in most; * sign-test q < 0.05).

**Supplementary Fig. 9.**
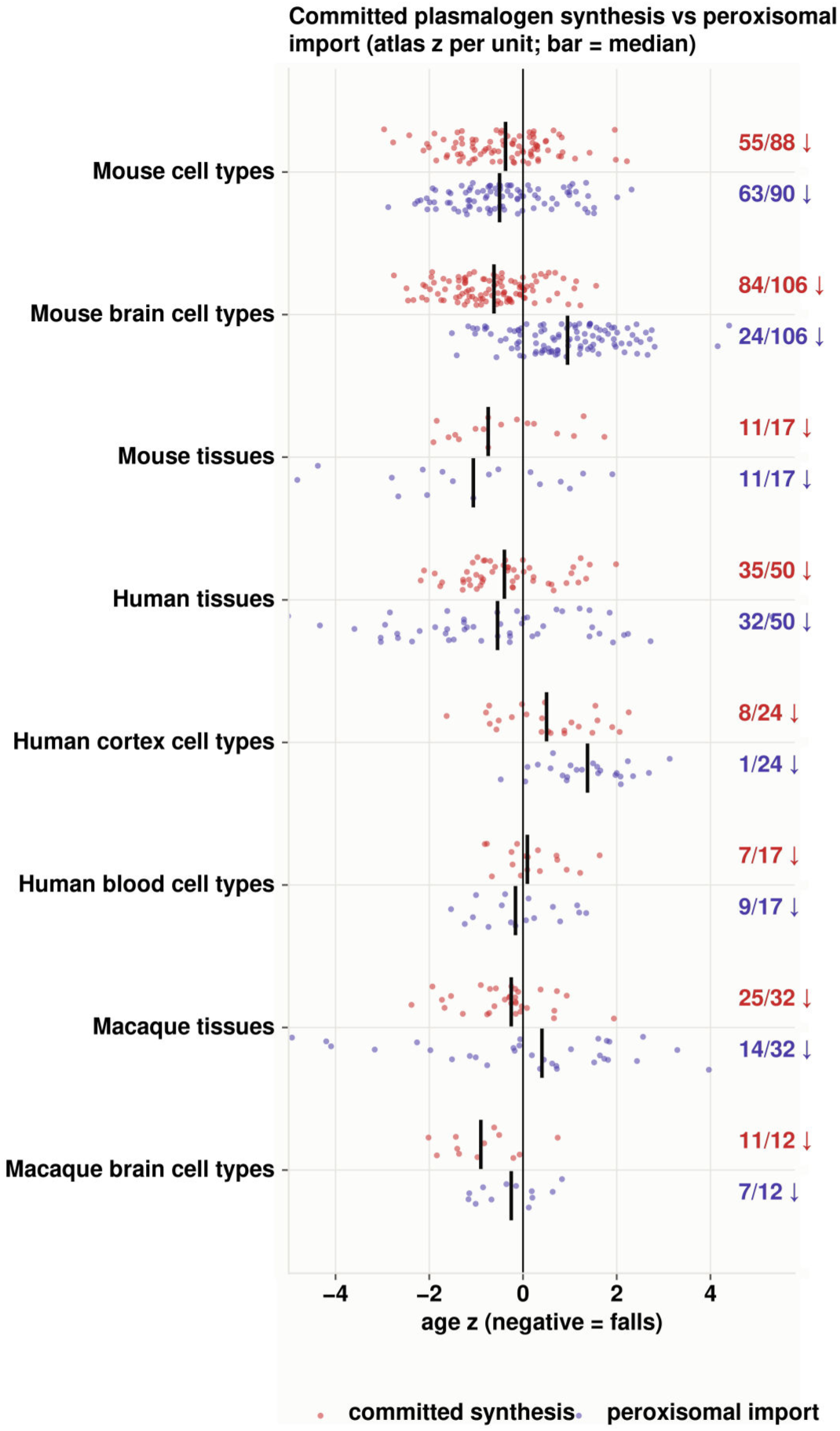
Programme atlas with matched-null scoring (part 2 of 3). Continued from Supplementary Fig. 8. Committed plasmalogen synthesis (red) and peroxisomal import (purple): per-unit age z (gene-level matched-null atlas z) for each dataset; bars = medians; text = units falling / tested.

**Supplementary Fig. 10.**
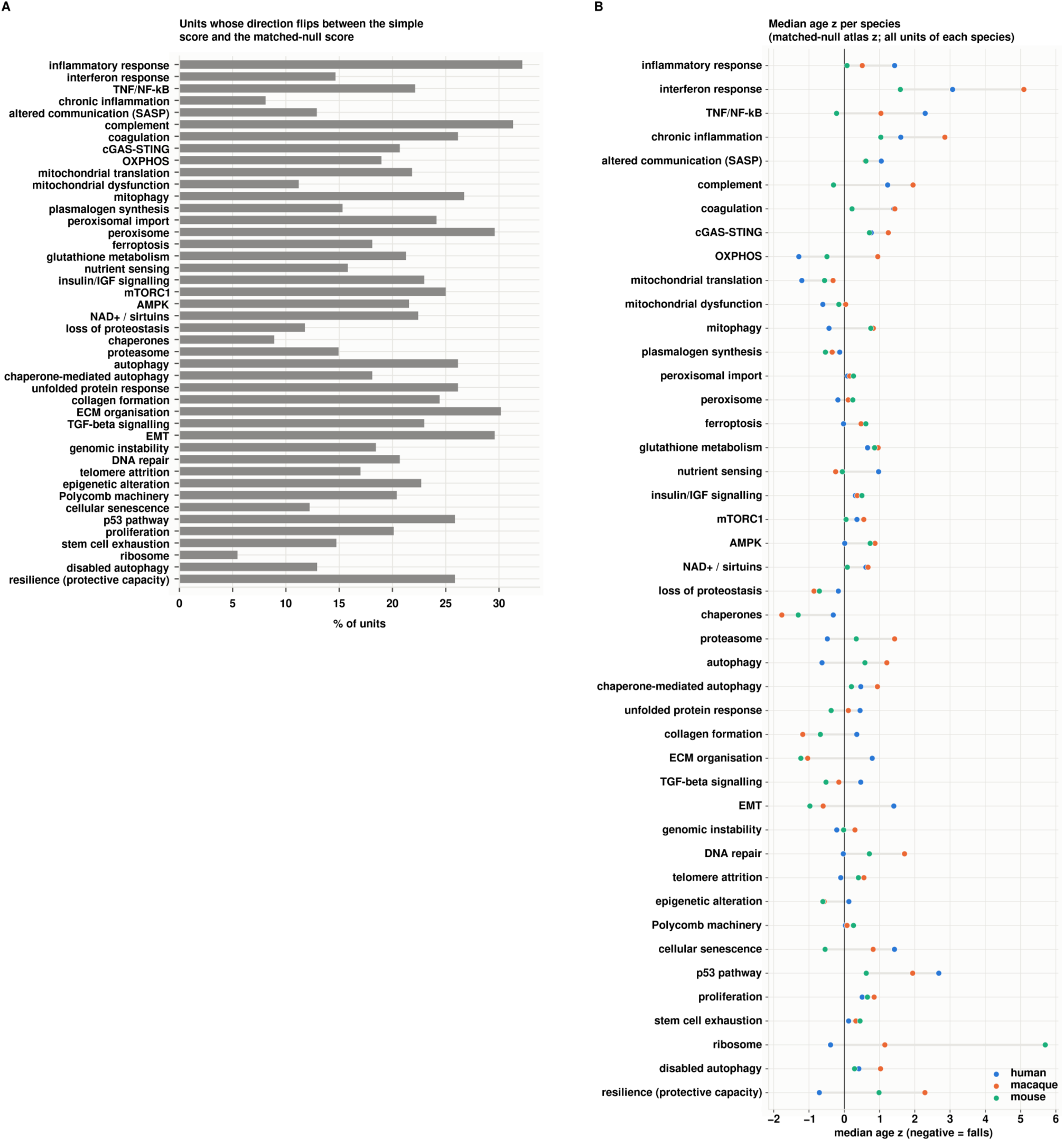
Programme atlas with matched-null scoring (part 3 of 3). Continued from Supplementary Fig. 8. **A,** Share of units whose direction differs between the earlier simple score and the matched-null score, per programme. **B,** Median age z per species (human, macaque, mouse) for each programme.

**Supplementary Fig. 11.**
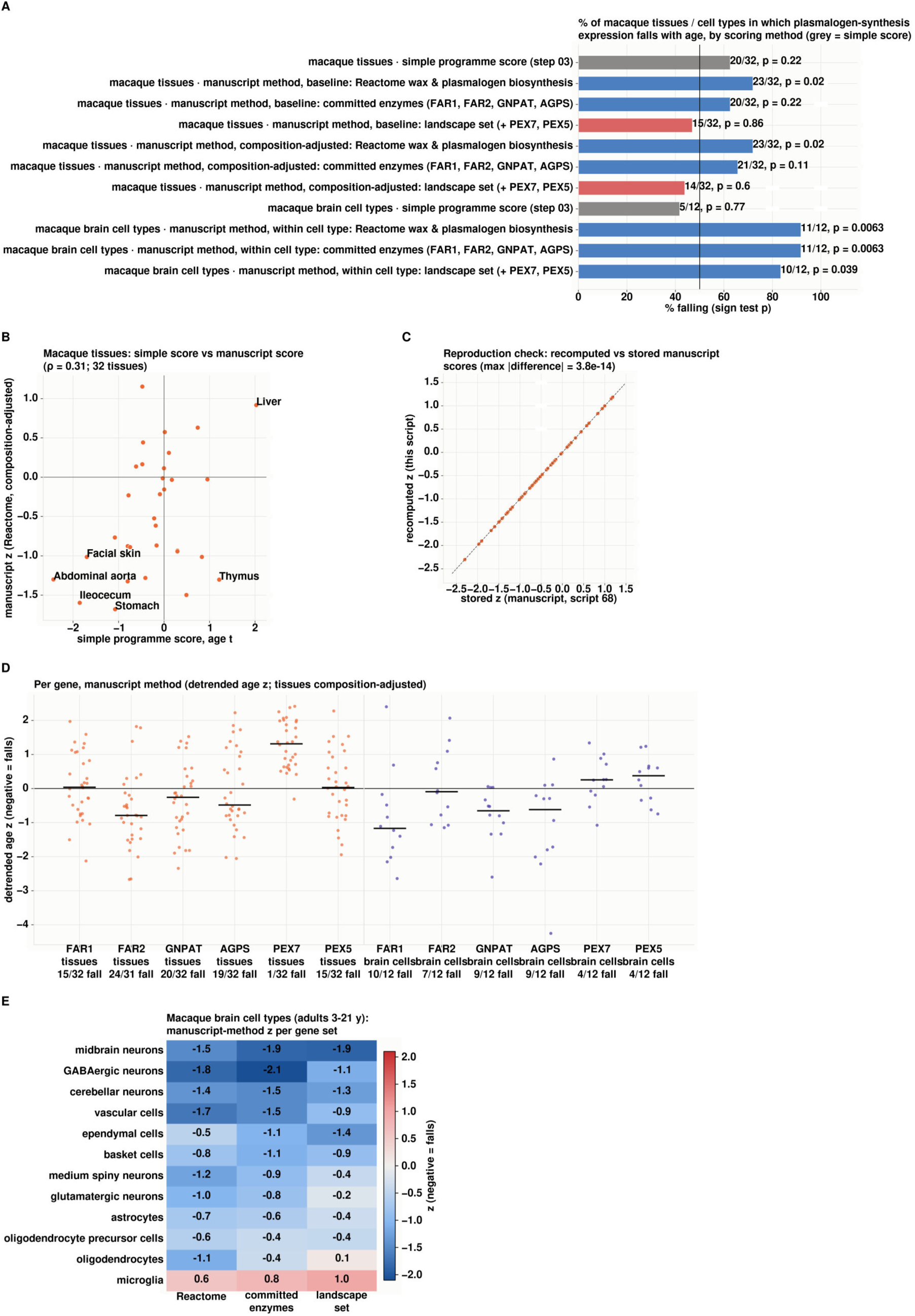
Macaque plasmalogen-synthesis expression with the manuscript’s method. **A,** Percentage of macaque tissues and brain cell types in which plasmalogen-synthesis expression falls with age, for the simple programme score on the committed enzymes (grey) and for the manuscript’s method with three gene sets (blue = falls in the majority; red = does not), baseline and composition-adjusted for tissues (sign-test p in text). **B,** Per tissue: simple score (x) against the manuscript’s composition-adjusted Reactome z (y). **C,** Reproduction check: z recomputed by this script against the stored manuscript values. **D,** Per gene, manuscript method (tissues composition-adjusted). **E,** Brain cell types: manuscript-method z per gene set.

**Supplementary Fig. 12.**
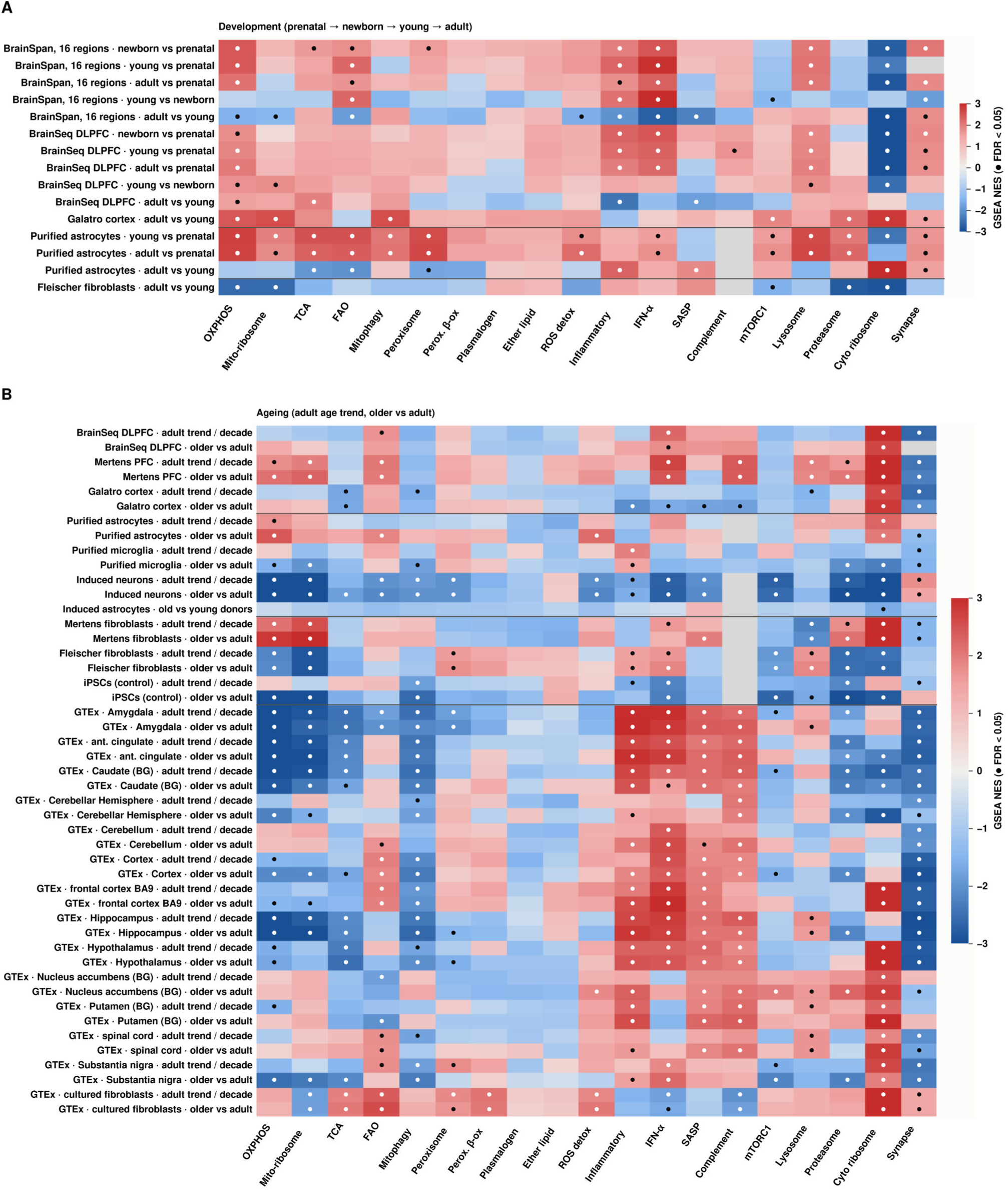
Human lifespan resource: focus modules in every unit and contrast. Normalised enrichment scores (NES; fgsea on the limma moderated t) of 19 curated modules: OXPHOS, mitochondrial ribosome, TCA cycle, fatty-acid oxidation, mitophagy, peroxisome, peroxisomal β-oxidation, plasmalogen synthesis, ether lipid, ROS detoxification, inflammatory response, interferon-α, SASP, complement, mTORC1, lysosome, proteasome, cytoplasmic ribosome and synapse. **A,** Development contrasts. **B,** Ageing contrasts (adult age trend per decade; older versus adult). Rows, the units and contrasts of the human lifespan resource; horizontal lines separate bulk brain, purified or converted cells, cultured cells and GTEx units. Colour, NES (red, higher with age); ●, FDR < 0.05; grey, not testable.

**Supplementary Fig. 13.**
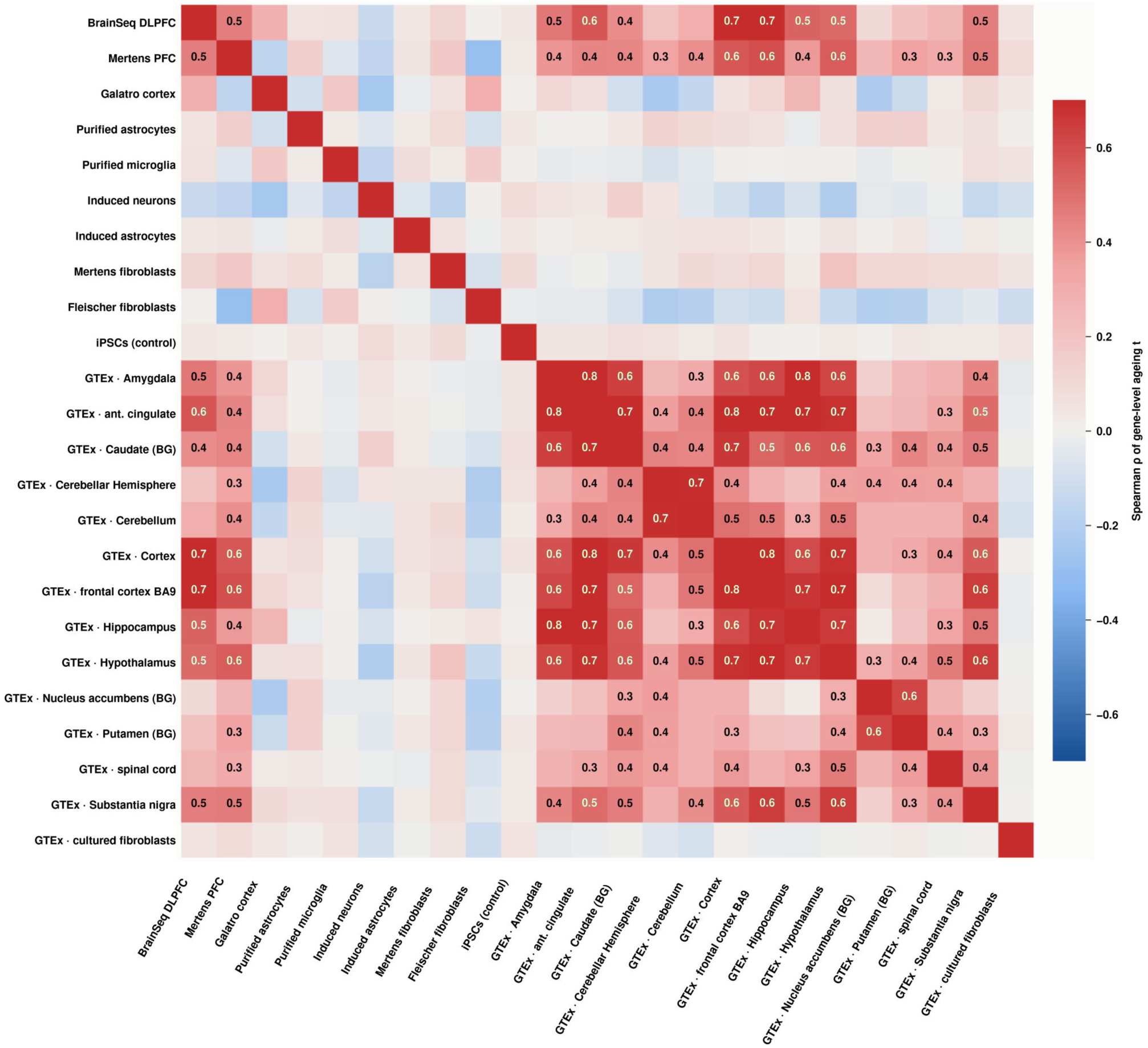
Similarity of gene-level ageing signatures between human units. Spearman ρ between the gene-level moderated t statistics of the adult age trend (or older versus adult) of every pair of units in the human lifespan resource and GTEx v8; values with |ρ| ≥ 0.3 are printed. Bulk brain series agree with each other (BrainSeq DLPFC versus GTEx frontal cortex, ρ = 0.70; GTEx brain regions 0.3–0.8), whereas purified cells, converted cells, the two fibroblast series and iPSCs share little with any other unit, as expected when their ageing signals are small or cell-type specific.

**Supplementary Fig. 14.**
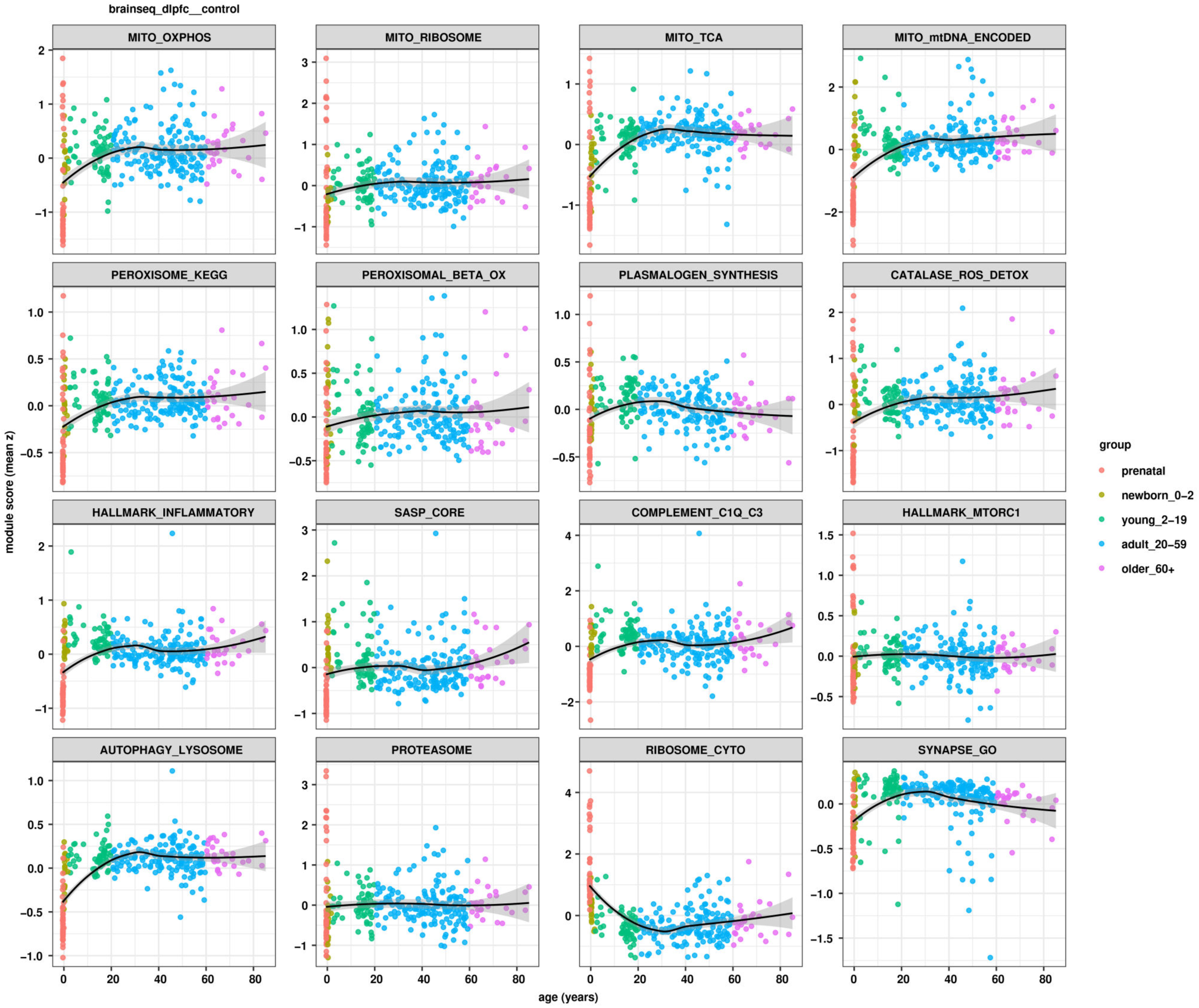
Module trajectories across the human lifespan in BrainSeq DLPFC. Per-sample module score (mean z of member genes across samples) against age for 16 focus modules (mitochondrial OXPHOS, ribosome, TCA and mtDNA-encoded genes; KEGG peroxisome, peroxisomal β-oxidation, plasmalogen synthesis, catalase/ROS detoxification; Hallmark inflammatory response, SASP core, complement C1Q/C3, Hallmark mTORC1; autophagy–lysosome, proteasome, cytoplasmic ribosome and GO synapse) in 320 control donors from prenatal life to 97 years. Points, samples coloured by age group (prenatal, newborn 0–2, young 2–19, adult 20–59, older 60+ years); line, loess fit with 95% confidence band. Mitochondrial and lysosomal modules rise steeply during development and then plateau or decline; plasmalogen synthesis peaks in adolescence and drifts down through adulthood.

**Supplementary Fig. 15.**
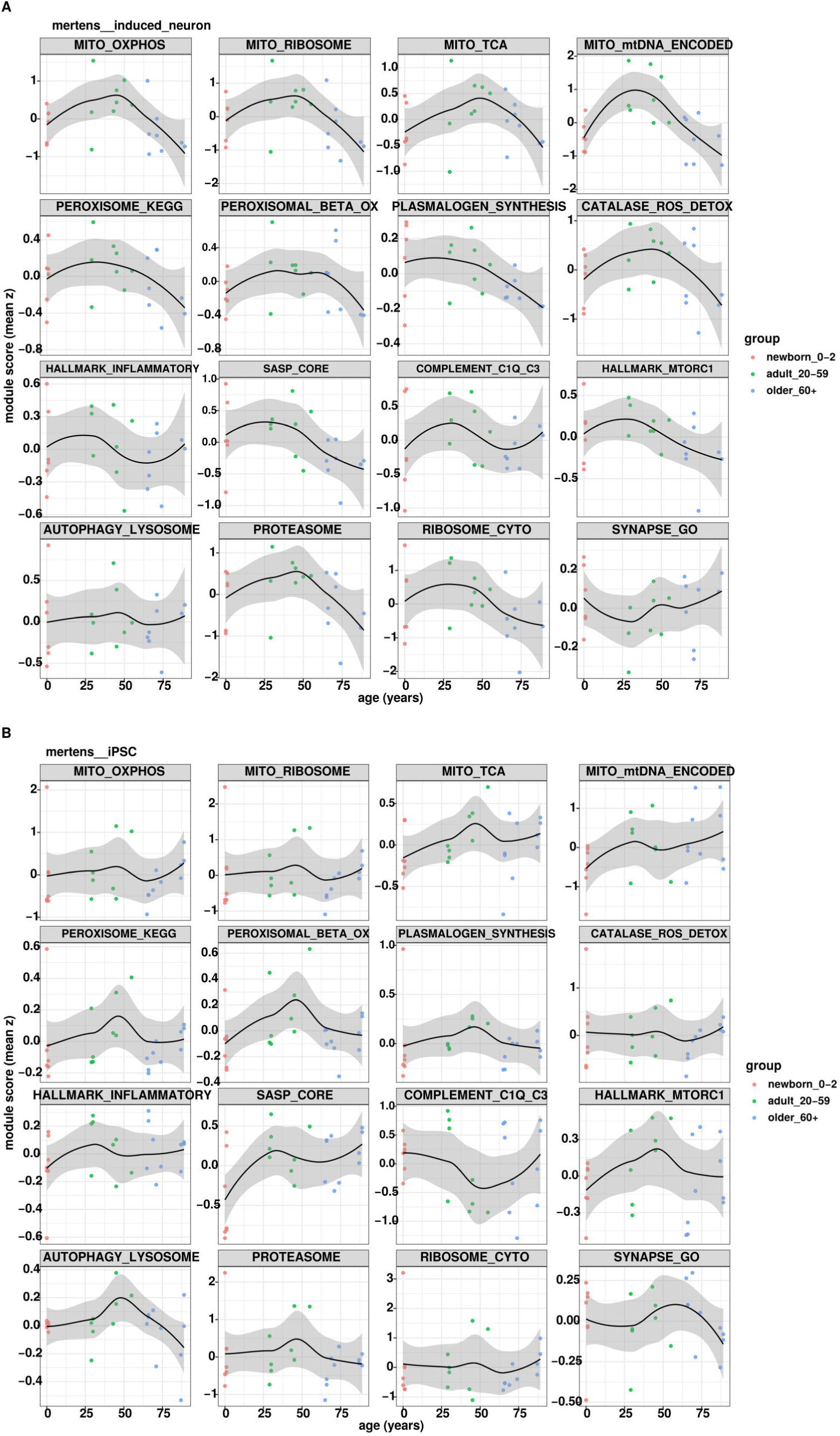
Module trajectories in directly induced neurons and in iPSCs from the same donors. As Supplementary Fig. 14. **A,** Neurons converted directly from the fibroblasts of donors aged 0–89 years (E-MTAB-3037). Direct conversion bypasses pluripotency and keeps the donor’s age signature, so trajectories follow donor age in cells whose composition is fixed. **B,** Induced pluripotent stem cells reprogrammed from the same donors’ fibroblasts. Reprogramming erases the age signature: no module follows donor age, the negative control for the induced-neuron trajectories in **A**. Points, samples; line, loess fit with 95% confidence band.

**Supplementary Fig. 16.**
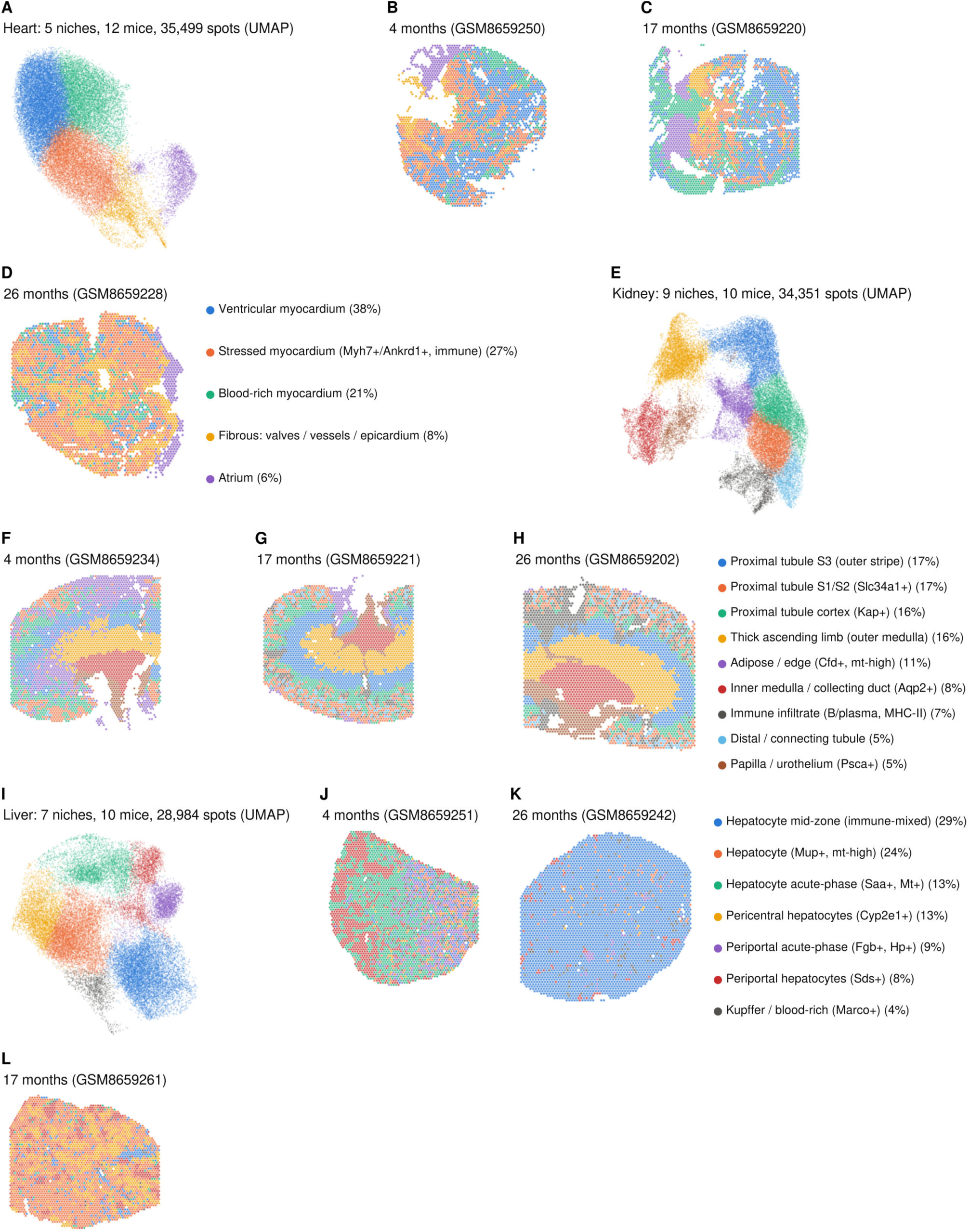
Spatial niches of the ageing mouse heart, kidney and liver. UMAP of Visium spots coloured by niche, and the niche map of one section per age (4, 17 and 26 months), for heart (**A–D**), kidney (**E–H**) and liver (**I–L**). Visium discovery cohort GSE283283 (male C57BL/6JN; 4, 17 and 26 months; 10–14 mice per tissue): spots passing QC were log-normalised, integrated with Harmony and clustered (Leiden) into niches, which were labelled from their marker genes before any age test, as set out in the pre-registered plan (Methods); niche labels and marker genes are given in Supplementary Data 5 (sheets niche_labels and niche_markers).

**Supplementary Fig. 17.**
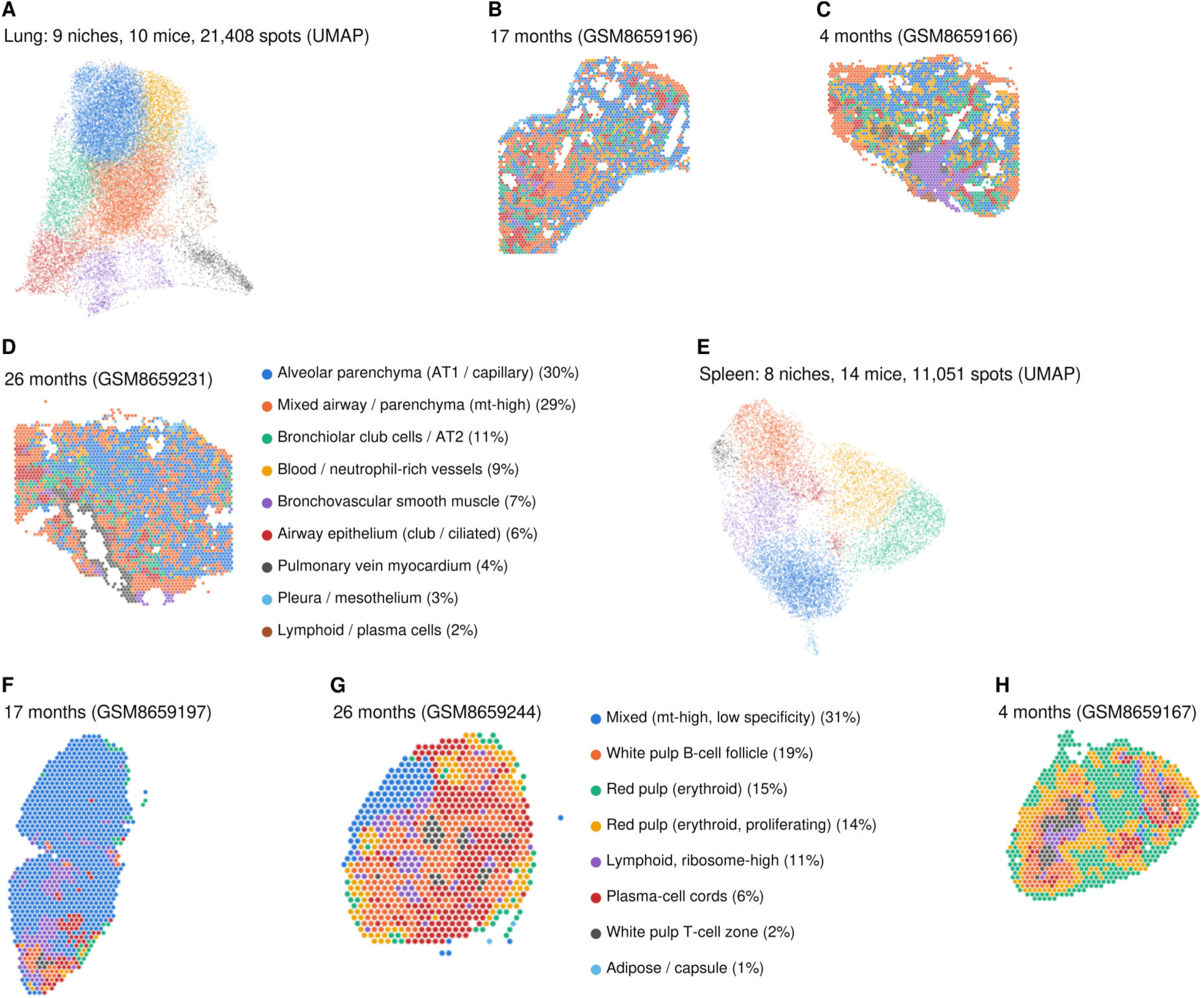
Spatial niches of the ageing mouse lung and spleen. UMAP of Visium spots coloured by niche, and the niche map of one section per age (4, 17 and 26 months), for lung (**A–D**) and spleen (**E–H**); the percentage after each niche name is its share of spots. Visium discovery cohort GSE283283 (male C57BL/6JN; 4, 17 and 26 months; 10–14 mice per tissue): spots passing QC were log-normalised, integrated with Harmony and clustered (Leiden) into niches, which were labelled from their marker genes before any age test, as set out in the pre-registered plan (Methods); niche labels and marker genes are given in Supplementary Data 5 (sheets niche_labels and niche_markers).

**Supplementary Fig. 18.**
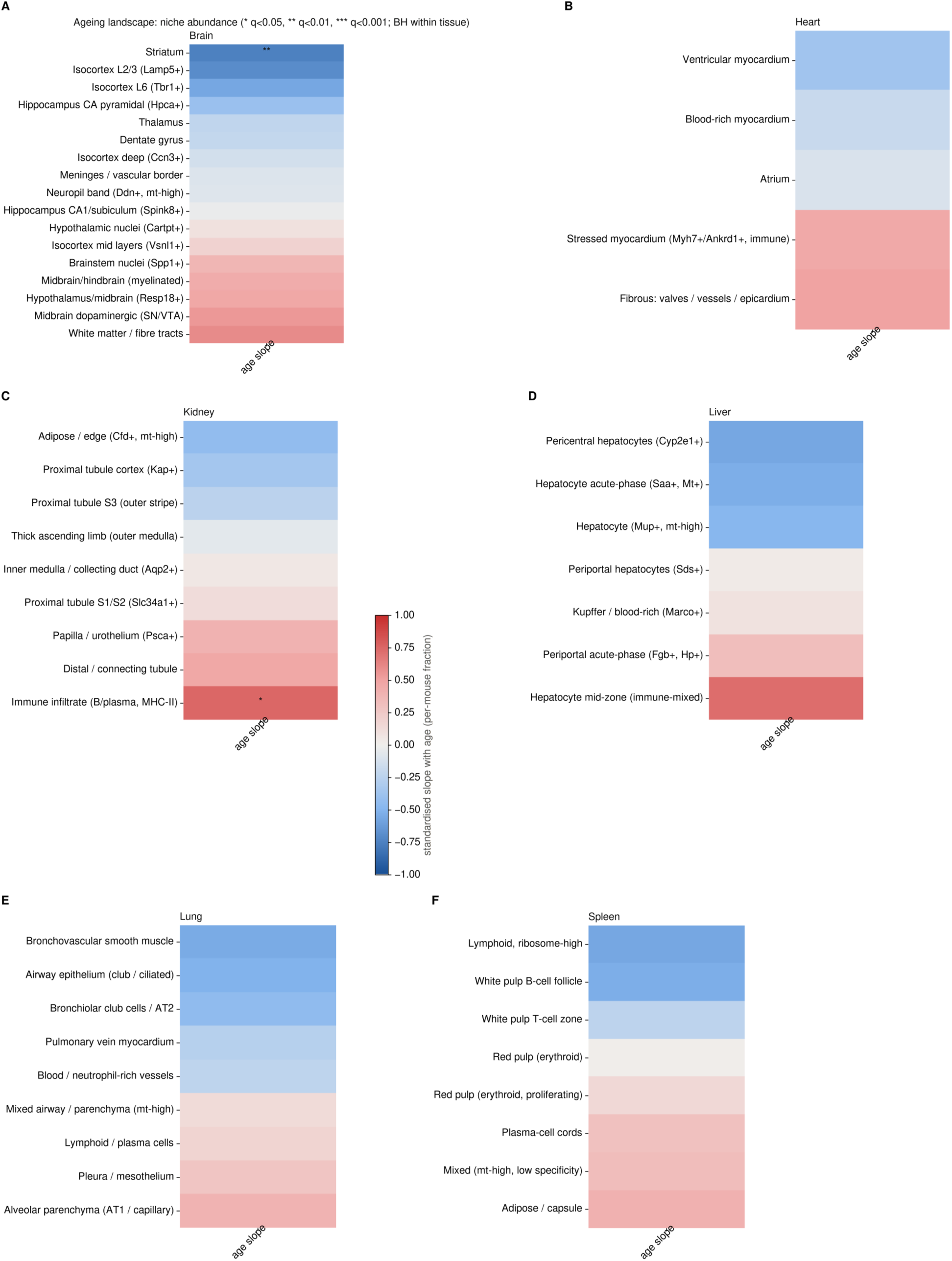
Niche abundance with age. Standardised age slope of each niche’s share of spots per mouse (linear model on age; mouse is the unit; * q < 0.05, ** q < 0.01, BH within tissue). **A,** Brain: the striatum fraction falls (q < 0.01; likely section level). **B,** Heart. **C,** Kidney: the immune-infiltrate niche (B/plasma, MHC-II) expands (q < 0.05). **D,** Liver. **E,** Lung. **F,** Spleen. GSE283283 Visium (male C57BL/6JN; 4, 17 and 26 months; 10–14 mice per tissue); niches labelled from marker genes before any age test (pre-registered plan).

**Supplementary Fig. 19.**
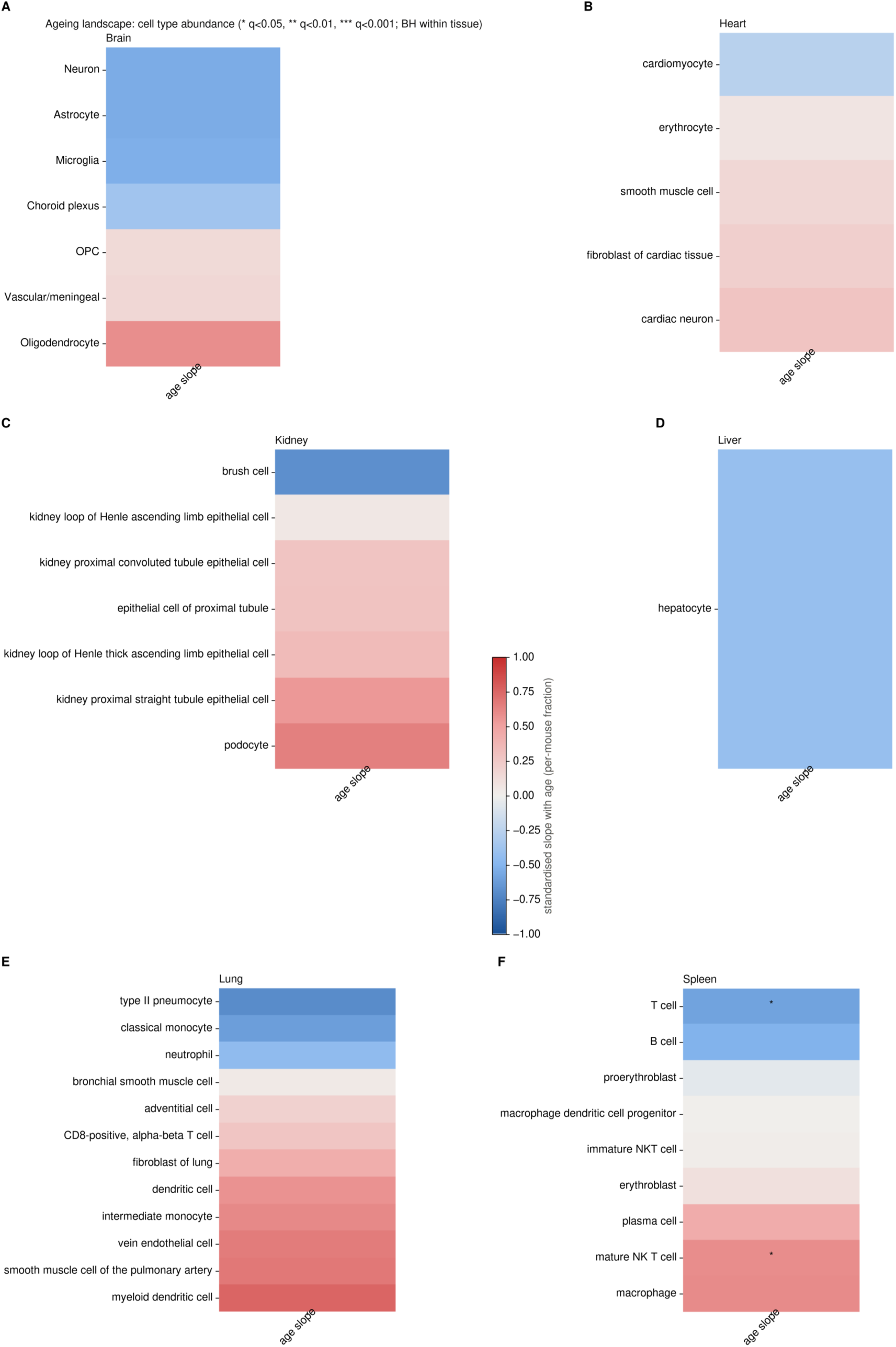
Cell-type proportions with age. Age slope of deconvolved cell-type proportions per mouse (corrected NNLS on Tabula Muris Senis for peripheral tissues, broad Stereo-seq classes for brain; * q < 0.05). **A,** Brain: oligodendrocytes rise, neurons, astrocytes and microglia fall in direction. **B,** Heart. **C,** Kidney. **D,** Liver (hepatocytes only resolved). **E,** Lung. **F,** Spleen: T cells fall and mature NK T cells rise (q < 0.05). A change is called robust only when deconvolution and marker scores agree (both q < 0.1; pre-registered amendment 3): robustly, liver hepatic stellate cells and plasmacytoid dendritic cells increase. GSE283283 Visium (male C57BL/6JN; 4, 17 and 26 months; 10–14 mice per tissue); niches labelled from marker genes before any age test (pre-registered plan).

**Supplementary Fig. 20.**
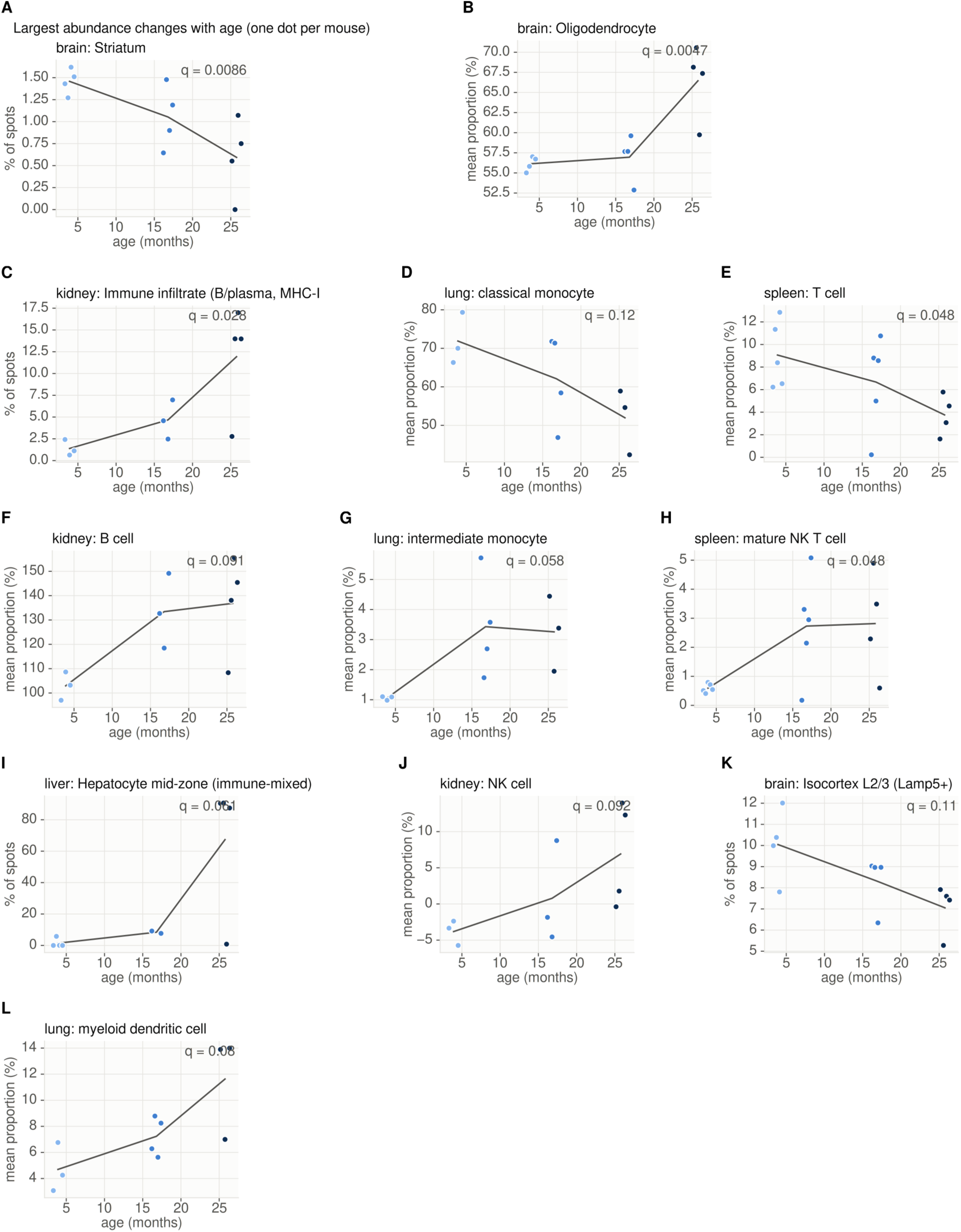
Largest abundance changes, per mouse. Per-mouse trajectories of the niches and cell types with the largest age slopes (points, mice coloured by age; line, mean per age; q from the age model): **A,** brain striatum niche (q = 0.0086); **B,** brain oligodendrocytes (q = 0.0047); **C,** kidney immune-infiltrate niche (q = 0.028); **D,** lung classical monocytes (q = 0.12); **E,** spleen T cells (q = 0.048); **F,** kidney B cells (q = 0.091); **G,** lung intermediate monocytes (q = 0.058); **H,** spleen mature NK T cells (q = 0.048); **I,** liver mid-zone hepatocyte niche (q = 0.061); **J,** kidney NK cells (q = 0.092); **K,** brain isocortex L2/3 niche (q = 0.11); **L,** lung myeloid dendritic cells (q = 0.08). Deconvolved values are unnormalised NNLS proportions and can exceed 100% (**F**). GSE283283 Visium (male C57BL/6JN; 4, 17 and 26 months; 10–14 mice per tissue); niches labelled from marker genes before any age test (pre-registered plan).

**Supplementary Fig. 21.**
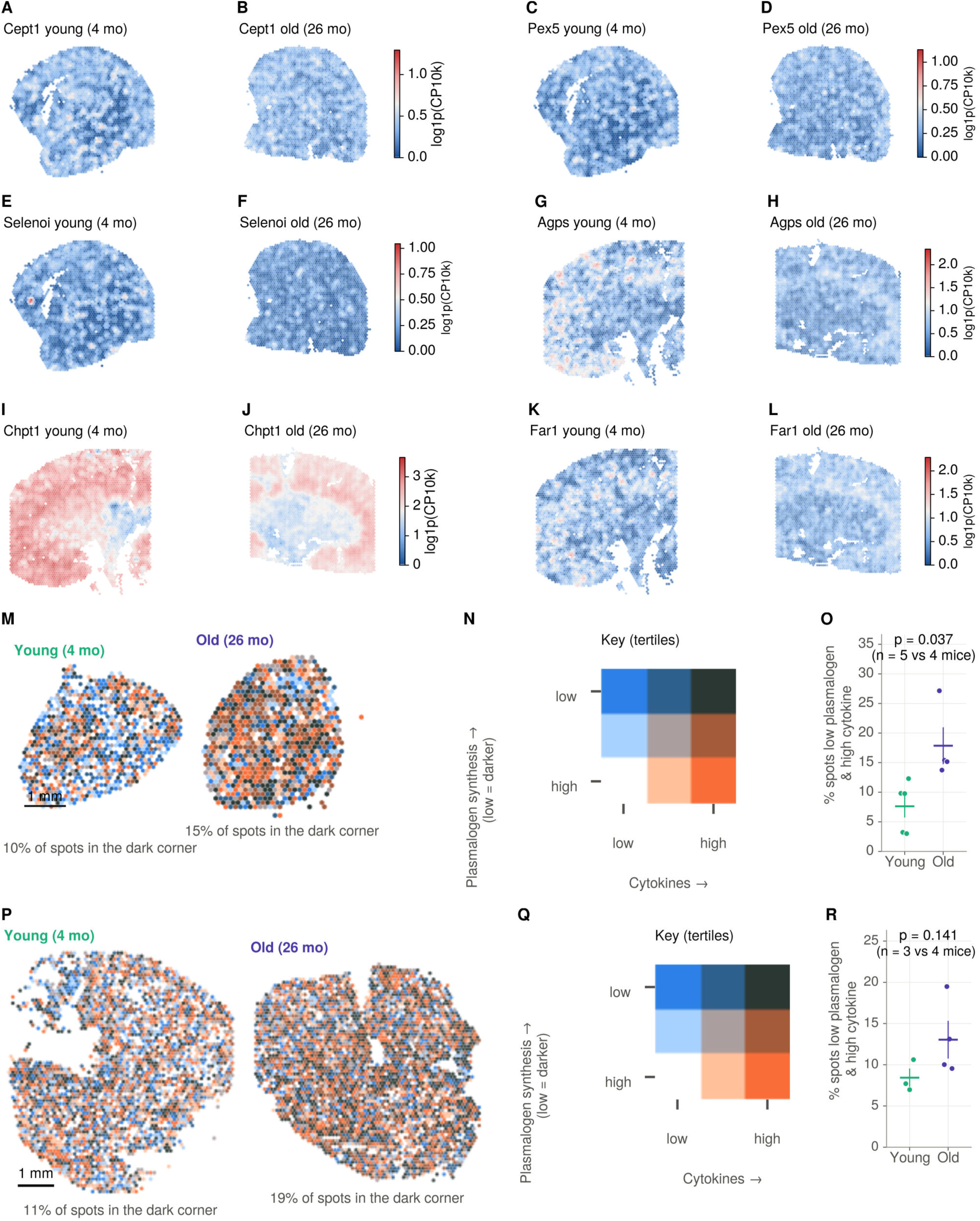
Spatial maps of single plasmalogen-pathway genes (brain, kidney) and of low plasmalogen-synthesis with high cytokine scores (spleen, heart). **A–L,** Expression maps (log-normalised, smoothed) of the plasmalogen-synthesis genes with the largest old-minus-young decrease, young versus old sections: **A–F,** brain; **G–L,** kidney. **M–O,** Bivariate map of low plasmalogen-synthesis and high cytokine scores, spleen. 3 × 3 bivariate colour per spot (dark corner: low plasmalogen-synthesis AND high cytokine score) in young and old spleen sections, with the per-mouse fraction of spots in the dark corner. **P–R,** The same for heart. GSE283283 Visium, young 3.9 versus old 25.8 months (3–5 mice per age and tissue; 17-month mice drawn for context, not tested); unit = mouse; Welch t-test and Hedges’ g, Benjamini–Hochberg (BH) within tissue. Secondary, descriptive analysis (not pre-registered).

**Supplementary Fig. 22.**
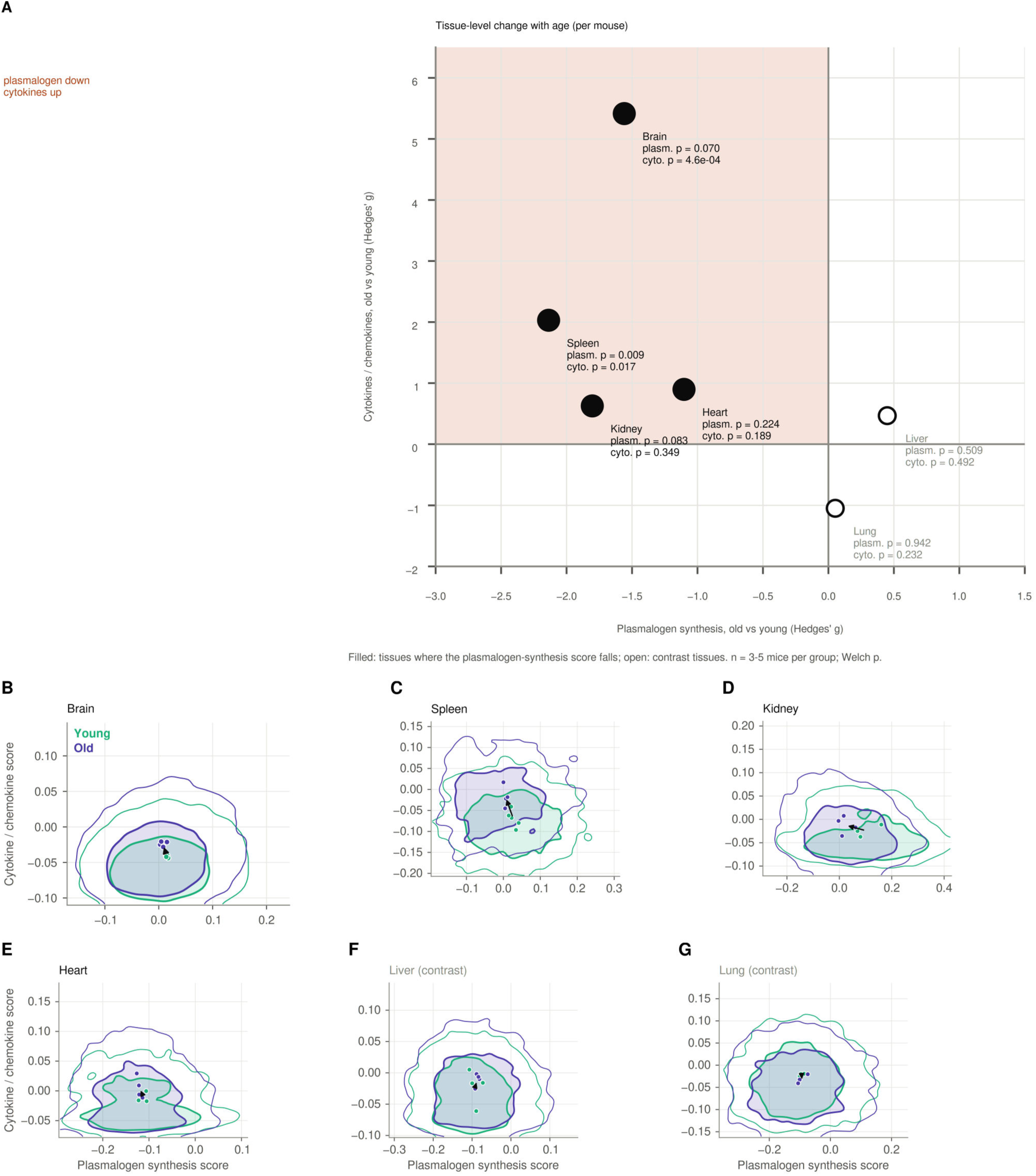
Tissue-level plasmalogen-synthesis and cytokine effect sizes; spots in plasmalogen-synthesis × cytokine score space. **A,** Tissue-level plasmalogen-synthesis and cytokine effect sizes. Hedges’ g (old versus young, per mouse) of the plasmalogen-synthesis score against that of the cytokine/chemokine score per tissue; filled, tissues where the plasmalogen-synthesis score falls (brain, spleen, kidney, heart); open, contrast tissues (liver, lung). Brain combines the largest cytokine rise (g = +5.4) with a falling plasmalogen-synthesis score. GSE283283 Visium, young 3.9 versus old 25.8 months (3–5 mice per age and tissue; 17-month mice drawn for context, not tested); unit = mouse; Welch t-test and Hedges’ g, Benjamini–Hochberg (BH) within tissue. Secondary, descriptive analysis (not pre-registered). **B–G,** Spots in plasmalogen-synthesis × cytokine score space. Two-dimensional density of all spots of each tissue in plasmalogen-synthesis × cytokine score space, young versus old; with age the cloud moves towards lower plasmalogen-synthesis and higher cytokine scores in brain and spleen. GSE283283 Visium, young 3.9 versus old 25.8 months (3–5 mice per age and tissue; 17-month mice drawn for context, not tested); unit = mouse; Welch t-test and Hedges’ g, Benjamini–Hochberg (BH) within tissue. Secondary, descriptive analysis (not pre-registered).

**Supplementary Fig. 23.**
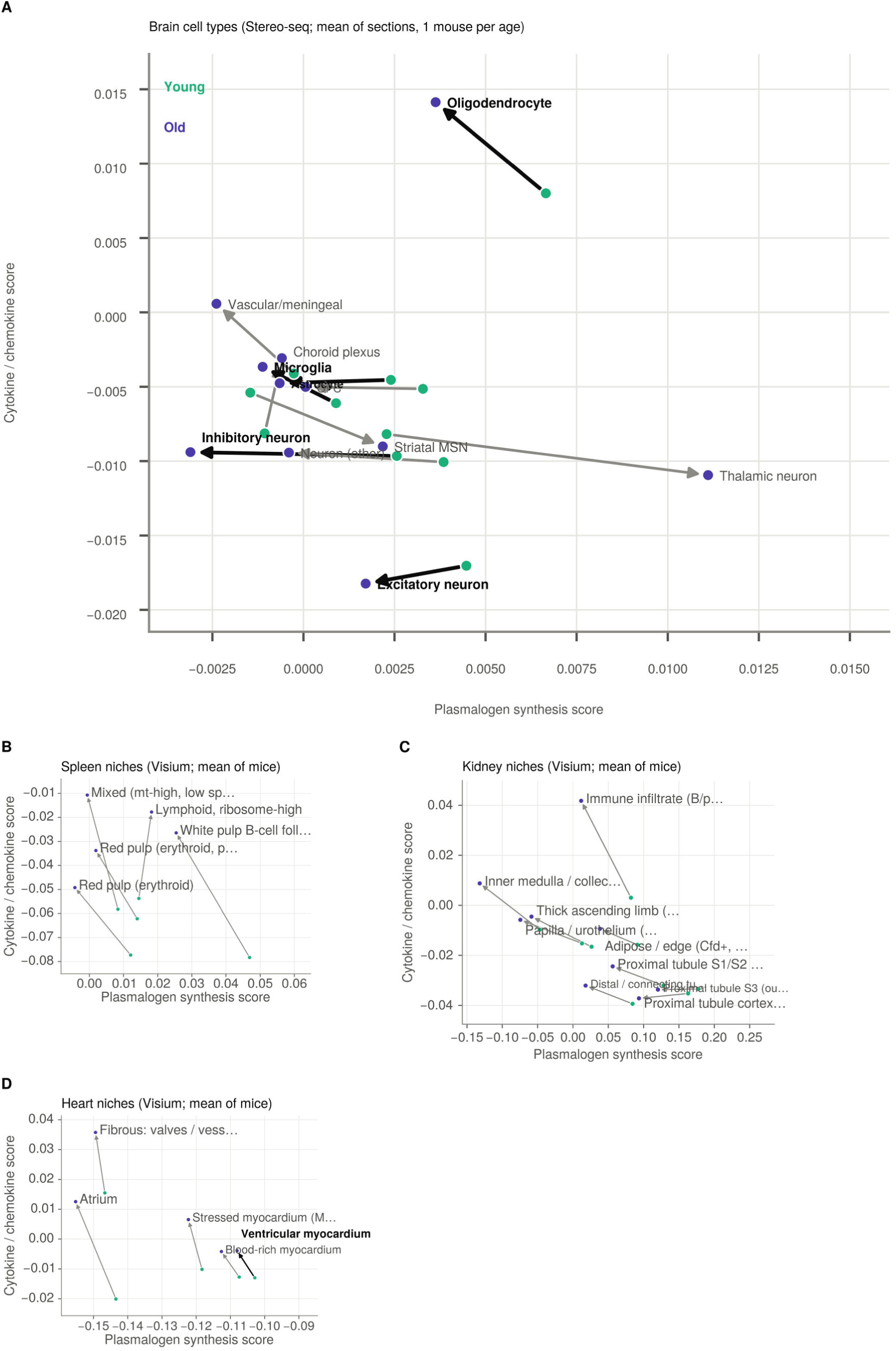
Cell types and niches move towards low plasmalogen-synthesis and high cytokine scores. Each cell type or niche drawn as an arrow from its young (green) to its old (purple) position in plasmalogen-synthesis × cytokine/chemokine score space. **A,** Stereo-seq brain cell types (mean of sections; one mouse per age, 4 and 26 months; descriptive). **B–D,** Visium niches (mean of mice) of **B,** spleen, **C,** kidney and **D,** heart: nearly every niche moves towards lower plasmalogen-synthesis and higher cytokine scores. Visium: GSE283283, young 3.9 versus old 25.8 months (3–5 mice per age and tissue); unit = mouse. Secondary, descriptive analysis (not pre-registered).

**Supplementary Fig. 24.**
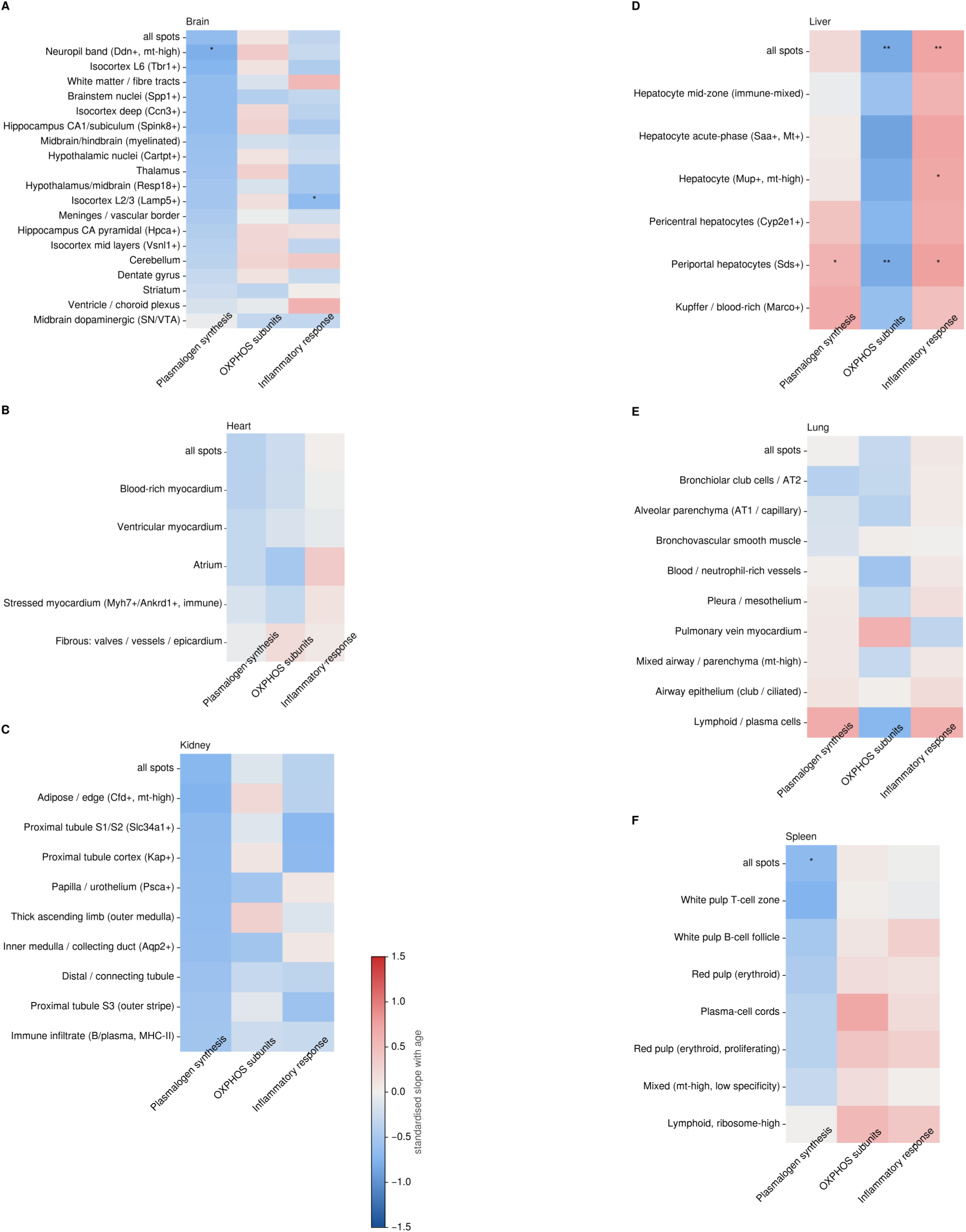
Programme age slopes per niche. Standardised age slope (per mouse) of the primary programmes — plasmalogen synthesis (16-gene focus set), OXPHOS subunits and inflammatory response — in all spots and in every niche; mean spot score relative to expression-matched control genes (scanpy score_genes); * q < 0.05 (BH within tissue). **A,** Brain: the plasmalogen-synthesis slope is negative in every niche (neuropil band q < 0.05). **B,** Heart: negative in every niche. **C,** Kidney: negative in every niche. **D,** Liver: plasmalogen synthesis rises in the periportal niche (q < 0.05), OXPHOS falls and inflammatory response rises across niches (all spots q < 0.01). **E,** Lung: no consistent change. **F,** Spleen: plasmalogen synthesis falls in all spots (q < 0.05) and in every niche. GSE283283 Visium (male C57BL/6JN; 4, 17 and 26 months; 10–14 mice per tissue); niches labelled from marker genes before any age test (pre-registered plan).

**Supplementary Fig. 25.**
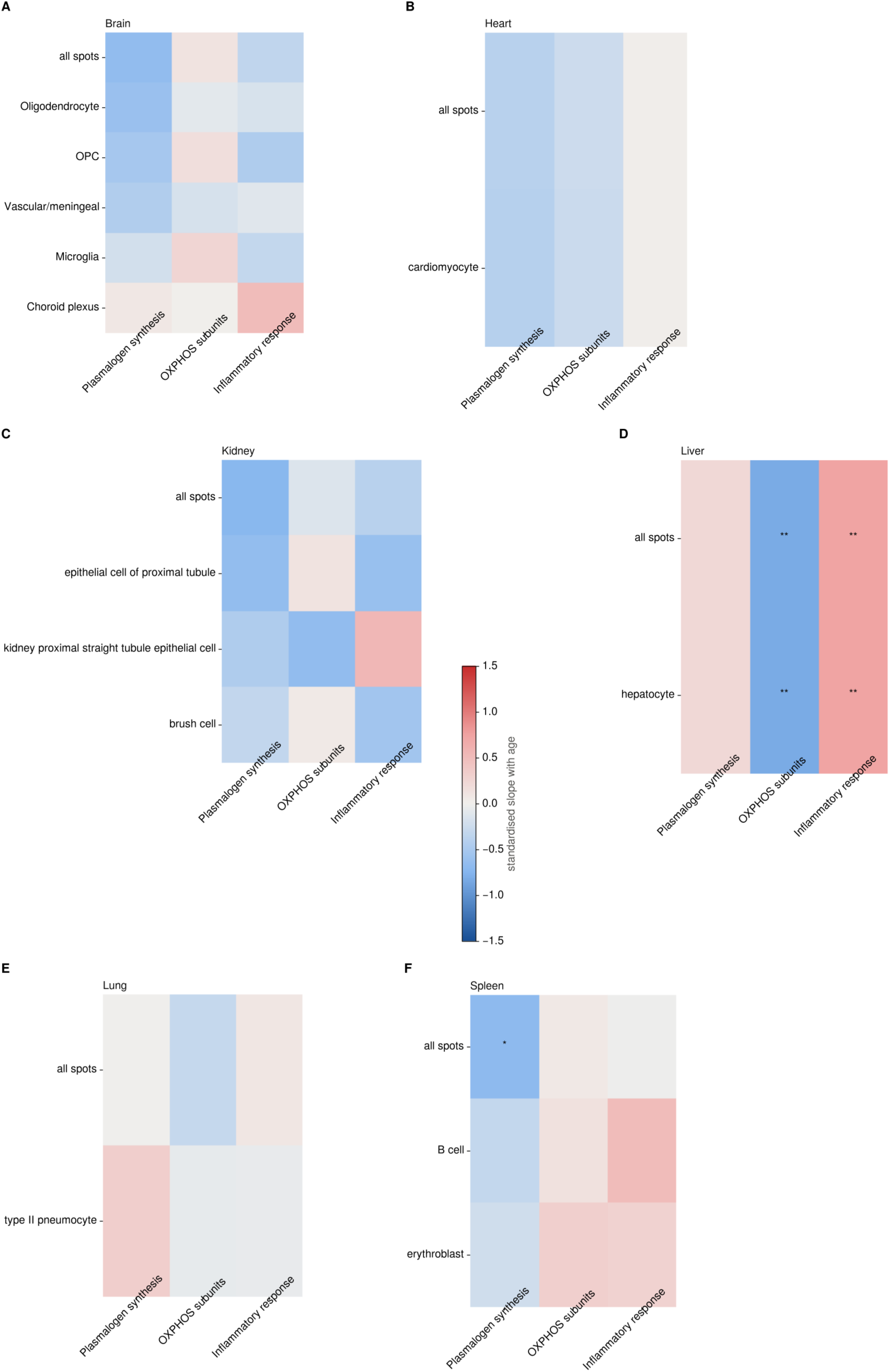
Programme age slopes per dominant cell type. As Supplementary Fig. 24, for spots grouped by their dominant deconvolved cell type. **A,** Brain: oligodendrocyte-, OPC- and vascular-dominated spots decline, choroid plexus does not. **B,** Heart (cardiomyocytes). **C,** Kidney (proximal-tubule epithelial and brush cells). **D,** Liver (hepatocytes): OXPHOS falls and inflammatory response rises (q < 0.01). **E,** Lung (type II pneumocytes): no decline. **F,** Spleen (B cells, erythroblasts): plasmalogen synthesis falls in all spots (q < 0.05). GSE283283 Visium (male C57BL/6JN; 4, 17 and 26 months; 10–14 mice per tissue); niches labelled from marker genes before any age test (pre-registered plan).

**Supplementary Fig. 26.**
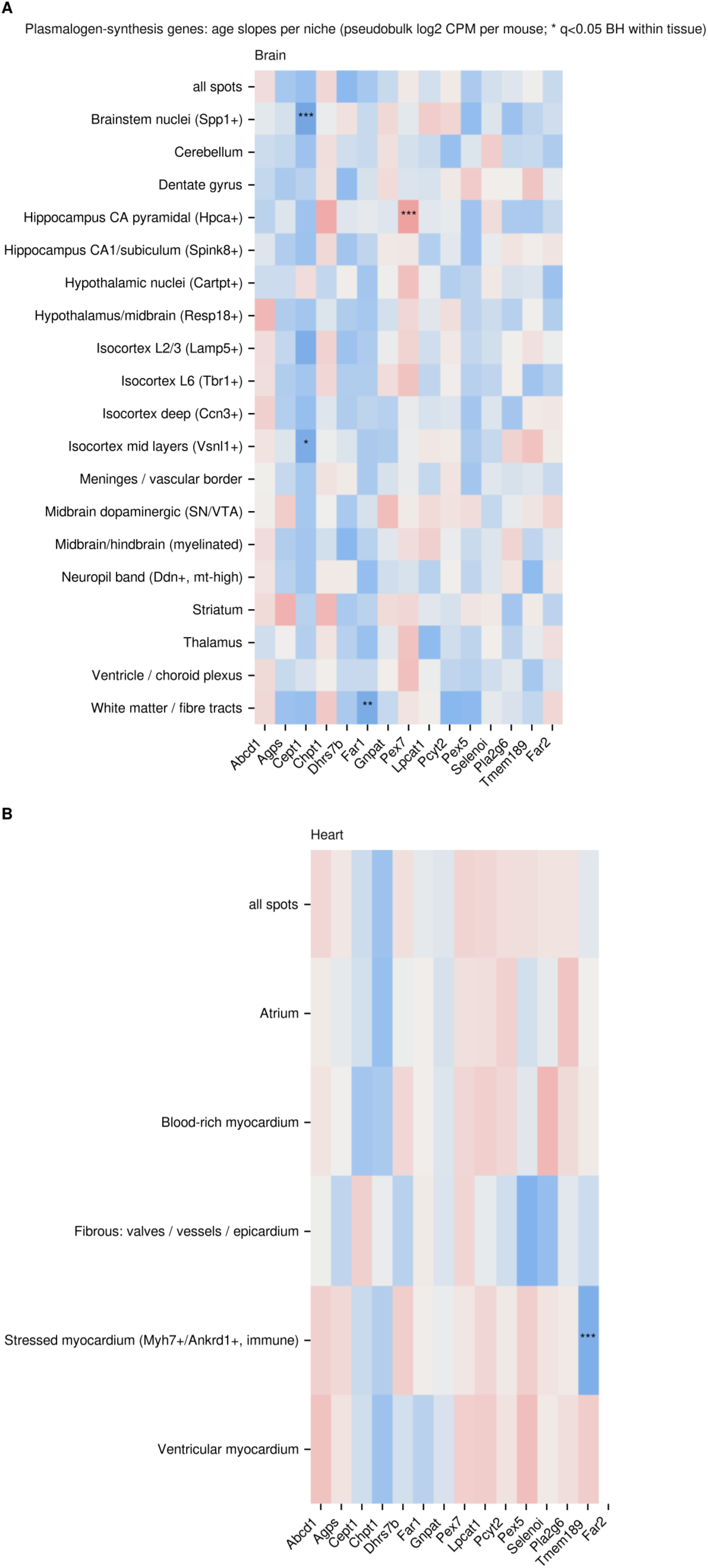
Single plasmalogen-pathway genes per niche (part 1 of 2). Standardised age slope of each plasmalogen-pathway gene (log2 CPM pseudobulk per mouse and niche; * q < 0.05, ** q < 0.01, *** q < 0.001, BH within tissue). **A,** Brain: *Cept1* falls in brainstem and mid-cortical niches and *Far1* in white matter; *Pex7* rises in the hippocampal pyramidal niche. **B,** Heart: *Far2* falls in stressed myocardium. GSE283283 Visium (male C57BL/6JN; 4, 17 and 26 months; 10–14 mice per tissue); niches labelled from marker genes before any age test (pre-registered plan).

**Supplementary Fig. 27.**
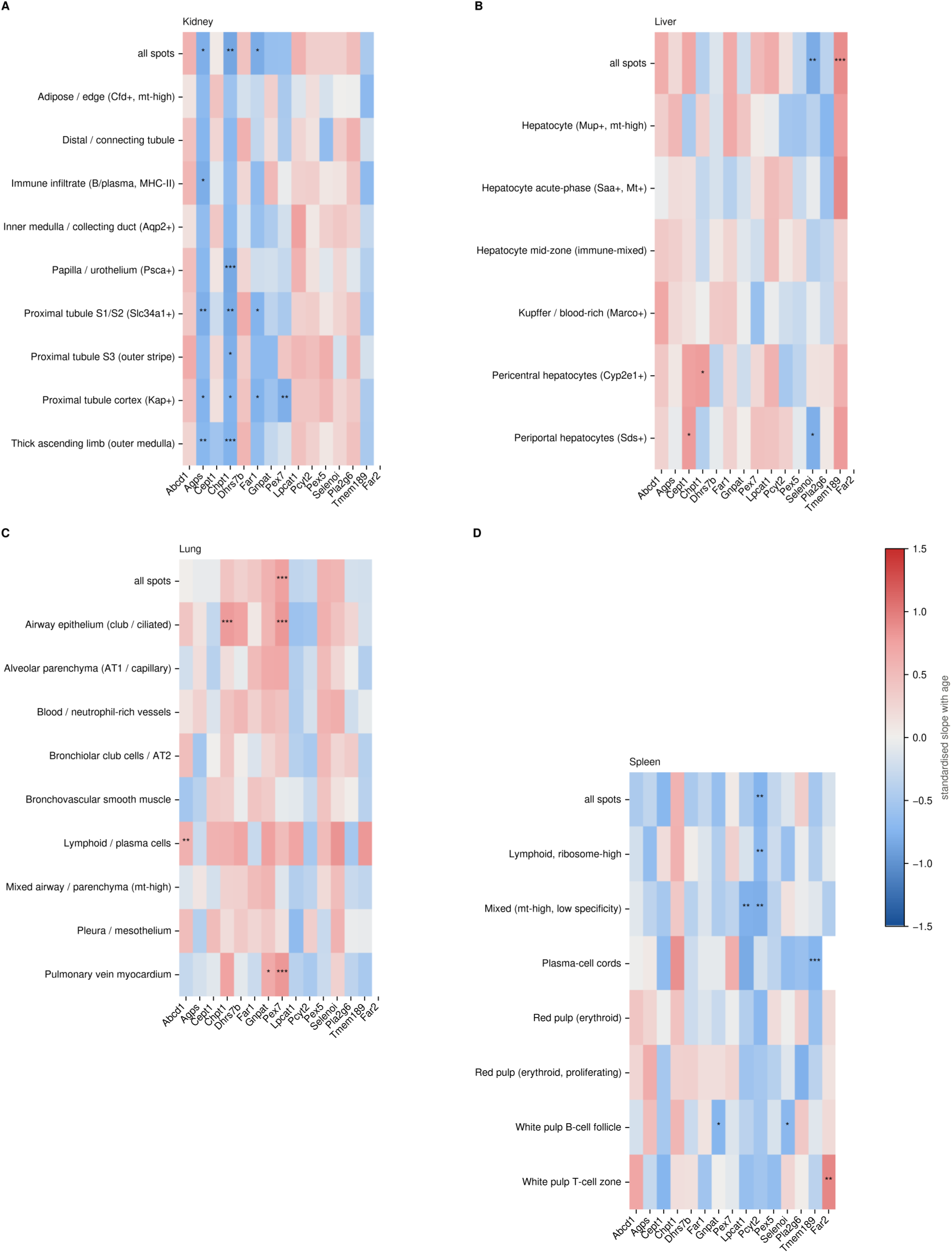
Single plasmalogen-pathway genes per niche (part 2 of 2). As Supplementary Fig. 26. **A,** Kidney: *Agps*, *Chpt1* and *Far1* fall in all spots and across tubule niches (*Far1* q = 0.048, *Agps* q = 0.014, *Chpt1* q = 0.005). **B,** Liver: *Pla2g6* falls and *Tmem189* rises. **C,** Lung: *Pex7* and *Chpt1* rise. **D,** Spleen: *Pcyt2* falls in all spots (q = 0.001) and several niches. GSE283283 Visium (male C57BL/6JN; 4, 17 and 26 months; 10–14 mice per tissue); niches labelled from marker genes before any age test (pre-registered plan).

**Supplementary Fig. 28.**
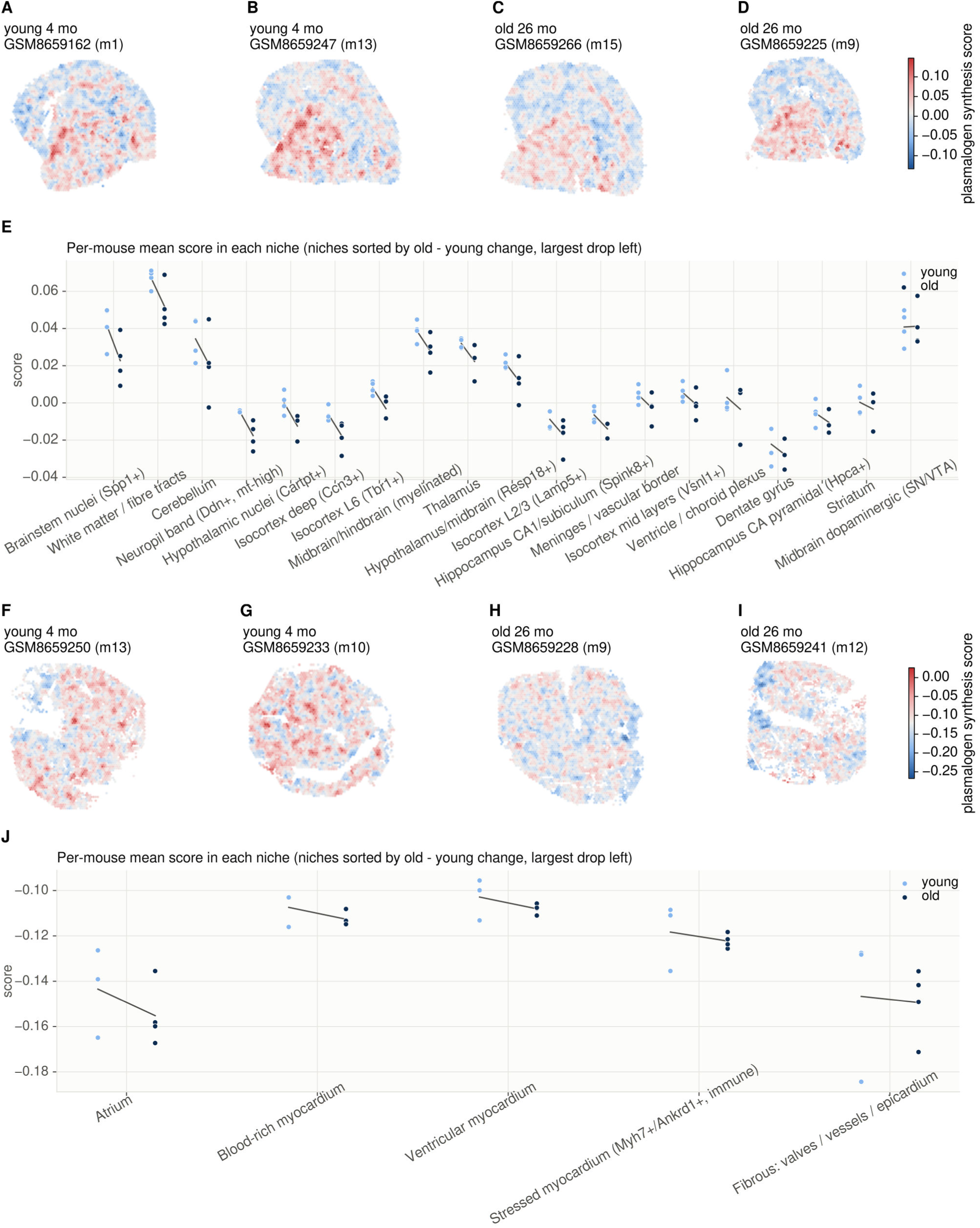
Spatial maps of the plasmalogen-synthesis score, brain and heart. **A–E,** Spatial maps of the plasmalogen-synthesis score, brain. Two young and two old brain sections on one colour scale (display smoothed over the 6 nearest spots), with each niche’s mean score per mouse below the maps, so that a whole-section shift can be told apart from a change confined to some niches. GSE283283 Visium, young 3.9 versus old 25.8 months (3–5 mice per age and tissue; 17-month mice drawn for context, not tested); unit = mouse; Welch t-test and Hedges’ g, Benjamini–Hochberg (BH) within tissue. Secondary, descriptive analysis (not pre-registered). **F–J,** Spatial maps of the plasmalogen-synthesis score, heart. Two young and two old heart sections on one colour scale (display smoothed over the 6 nearest spots), with each niche’s mean score per mouse below the maps, so that a whole-section shift can be told apart from a change confined to some niches. GSE283283 Visium, young 3.9 versus old 25.8 months (3–5 mice per age and tissue; 17-month mice drawn for context, not tested); unit = mouse; Welch t-test and Hedges’ g, Benjamini–Hochberg (BH) within tissue. Secondary, descriptive analysis (not pre-registered).

**Supplementary Fig. 29.**
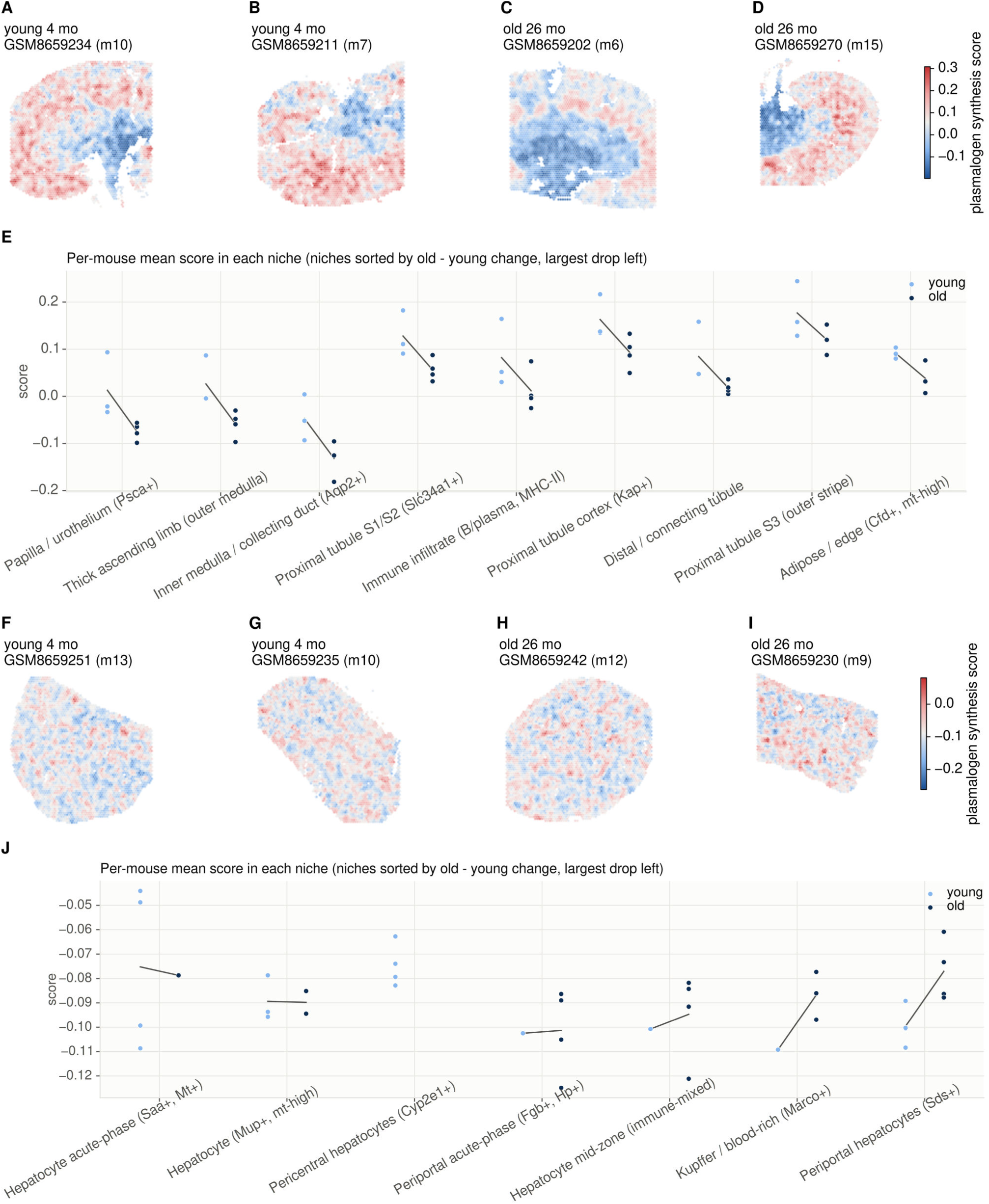
Spatial maps of the plasmalogen-synthesis score, kidney and liver. **A–E,** Spatial maps of the plasmalogen-synthesis score, kidney. Two young and two old kidney sections on one colour scale (display smoothed over the 6 nearest spots), with each niche’s mean score per mouse below the maps, so that a whole-section shift can be told apart from a change confined to some niches. GSE283283 Visium, young 3.9 versus old 25.8 months (3–5 mice per age and tissue; 17-month mice drawn for context, not tested); unit = mouse; Welch t-test and Hedges’ g, Benjamini–Hochberg (BH) within tissue. Secondary, descriptive analysis (not pre-registered). **F–J,** Spatial maps of the plasmalogen-synthesis score, liver. Two young and two old liver sections on one colour scale (display smoothed over the 6 nearest spots), with each niche’s mean score per mouse below the maps, so that a whole-section shift can be told apart from a change confined to some niches. GSE283283 Visium, young 3.9 versus old 25.8 months (3–5 mice per age and tissue; 17-month mice drawn for context, not tested); unit = mouse; Welch t-test and Hedges’ g, Benjamini–Hochberg (BH) within tissue. Secondary, descriptive analysis (not pre-registered).

**Supplementary Fig. 30.**
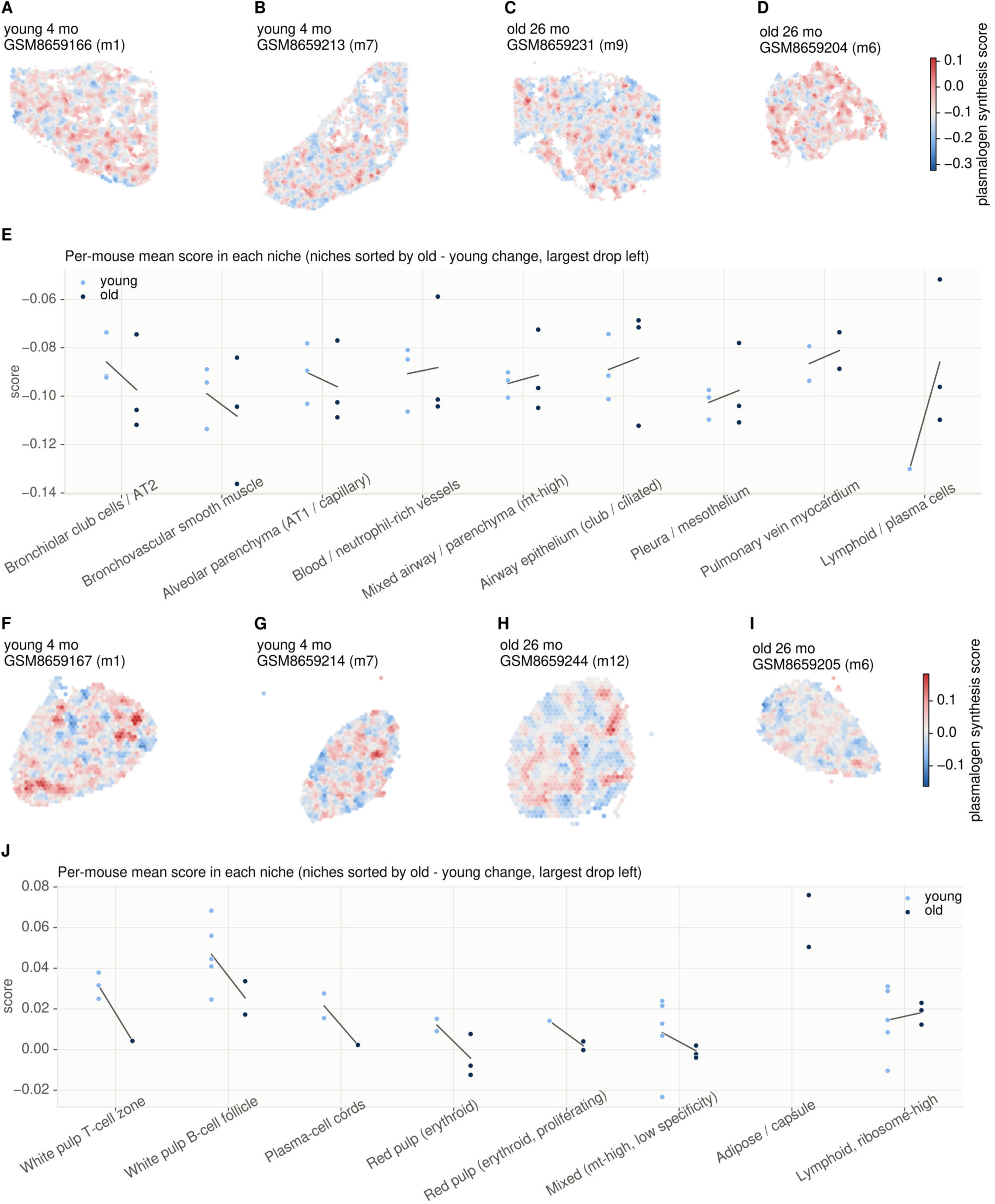
Spatial maps of the plasmalogen-synthesis score, lung and spleen. **A–E,** Spatial maps of the plasmalogen-synthesis score, lung. Two young and two old lung sections on one colour scale (display smoothed over the 6 nearest spots), with each niche’s mean score per mouse below the maps, so that a whole-section shift can be told apart from a change confined to some niches. GSE283283 Visium, young 3.9 versus old 25.8 months (3–5 mice per age and tissue; 17-month mice drawn for context, not tested); unit = mouse; Welch t-test and Hedges’ g, Benjamini–Hochberg (BH) within tissue. Secondary, descriptive analysis (not pre-registered). **F–J,** Spatial maps of the plasmalogen-synthesis score, spleen. Two young and two old spleen sections on one colour scale (display smoothed over the 6 nearest spots), with each niche’s mean score per mouse below the maps, so that a whole-section shift can be told apart from a change confined to some niches. GSE283283 Visium, young 3.9 versus old 25.8 months (3–5 mice per age and tissue; 17-month mice drawn for context, not tested); unit = mouse; Welch t-test and Hedges’ g, Benjamini–Hochberg (BH) within tissue. Secondary, descriptive analysis (not pre-registered).

**Supplementary Fig. 31.**
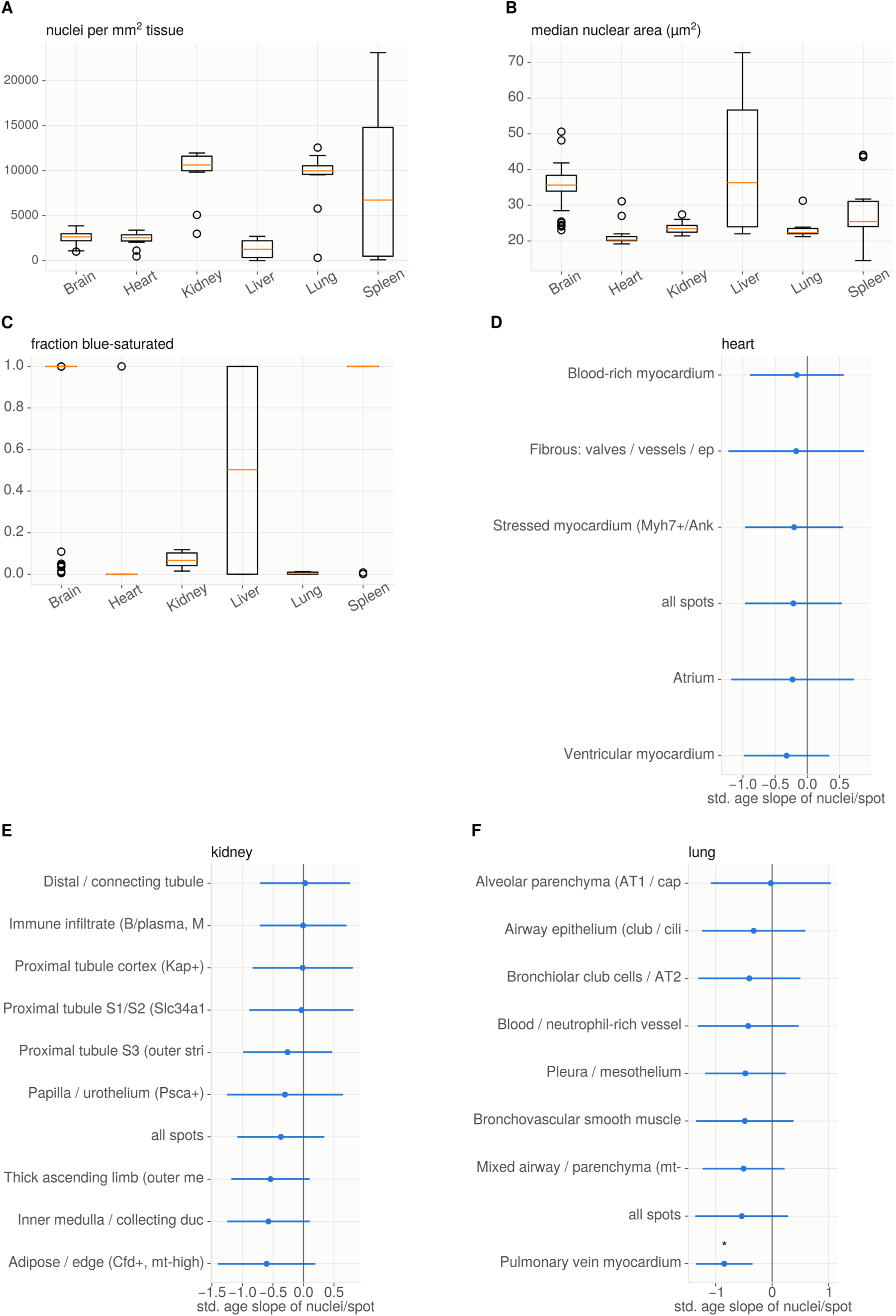
Nuclear segmentation of the H&E images: quality control and cellularity with age. **A–C,** Quality control of the nuclear segmentation per section (nuclei segmented in the full-resolution H&E images; pre-registered Addenda A and B): **A,** nuclei per mm² of tissue; **B,** median nuclear area; **C,** fraction of blue-saturated (over-stained) nuclei. Registration checks required the nuclei count to separate in-from off-tissue spots (AUC ≥ 0.70 and ≥ 0.05 above the AUC with mirrored coordinates). Spleen sections did not pass and are excluded. **D–F,** Cellularity with age. Standardised age slope (± 95% CI) of mean nuclei per spot per niche (OLS with HC3 errors, BH within tissue; * q < 0.05). **D,** Heart and **E,** kidney: no niche changes. **F,** Lung: the pulmonary-vein myocardium loses nuclei (q = 0.008). Brain, liver and spleen give no estimate and are not shown. Nuclei segmented in the full-resolution H&E images (pre-registered Addenda A and B).

**Supplementary Fig. 32.**
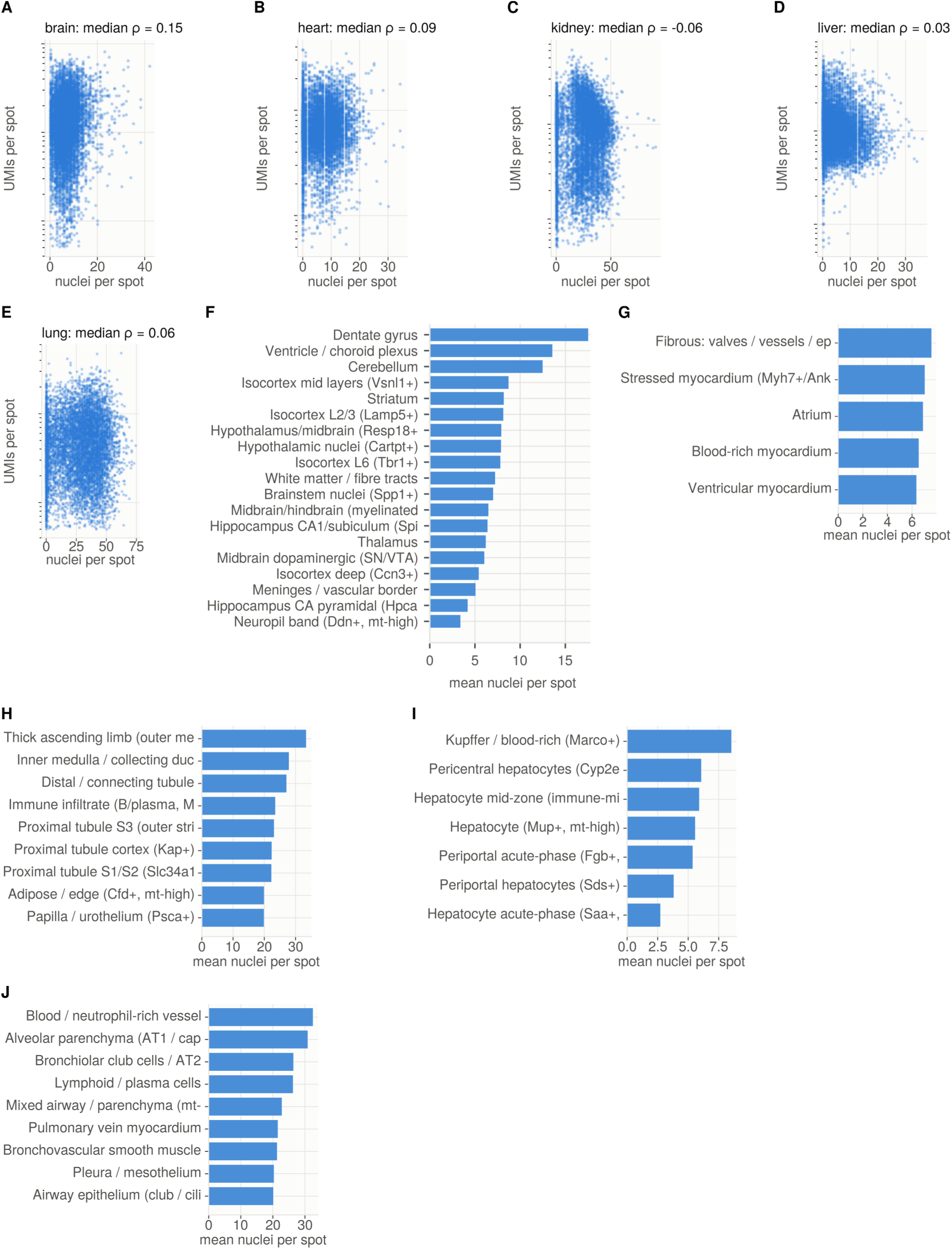
Image validation of the transcriptomic niches. **A–E,** Nuclei per spot against UMIs per spot in brain, heart, kidney, liver and lung (median Spearman ρ per section −0.06 to +0.15; spleen excluded): within sections the two are only weakly related. **F–J,** Mean nuclei per spot per niche in brain, heart, kidney, liver and lung: dentate gyrus, choroid plexus and cerebellum carry the most nuclei in brain, tubular niches in kidney. Nuclei segmented in the full-resolution H&E images (pre-registered Addenda A and B).

**Supplementary Fig. 33.**
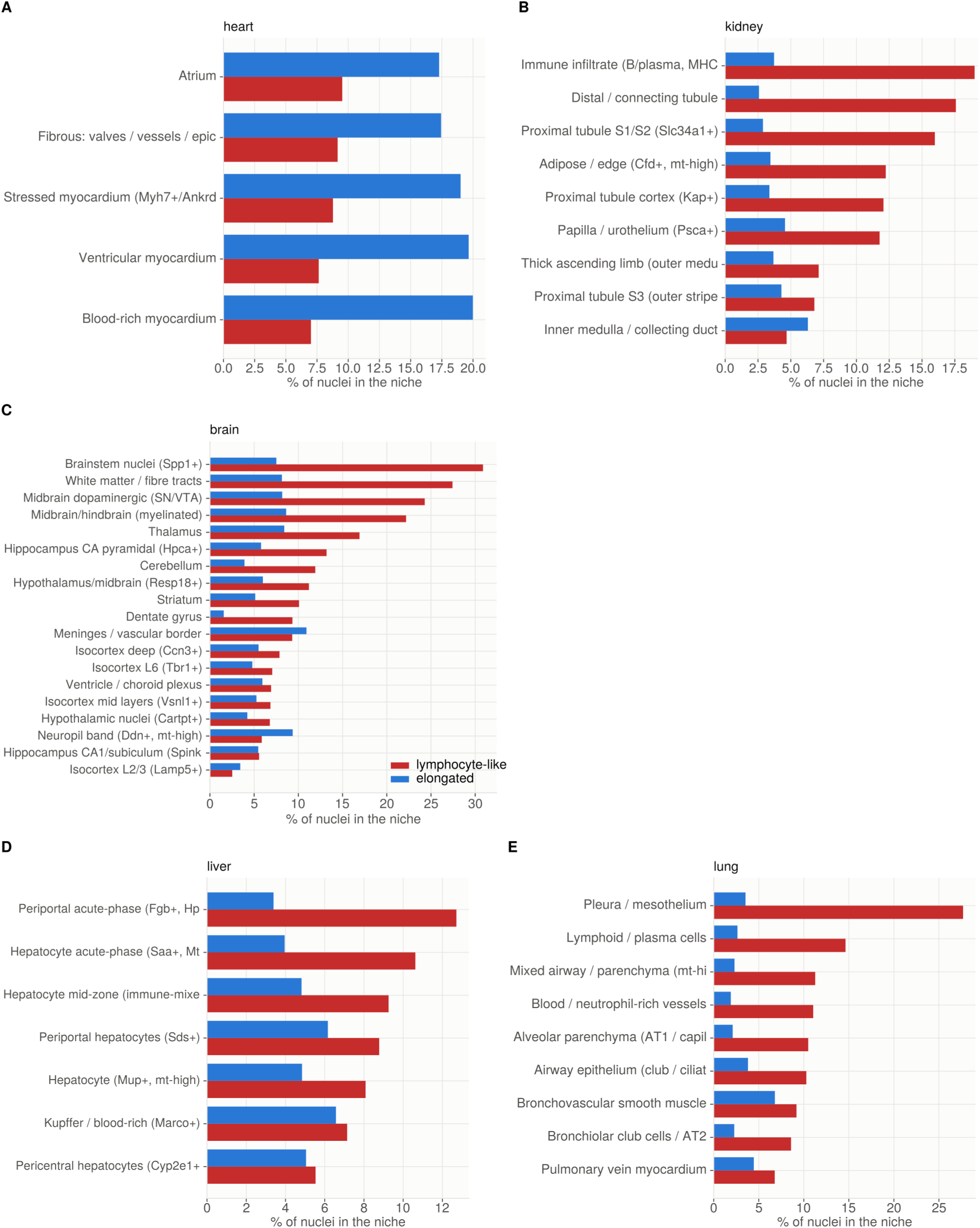
Nuclear morphology classes per niche. Lymphocyte-like (red) and elongated (blue) nuclei as % of nuclei in each niche (section-relative rules fixed in pre-registered Addendum B): **A,** heart; **B,** kidney (immune-infiltrate niche highest in lymphocyte-like nuclei); **C,** brain (brainstem and white-matter niches highest); **D,** liver; **E,** lung (pleural and lymphoid niches highest). Spleen sections were excluded. Nuclei segmented in the full-resolution H&E images (pre-registered Addenda A and B).

**Supplementary Fig. 34.**
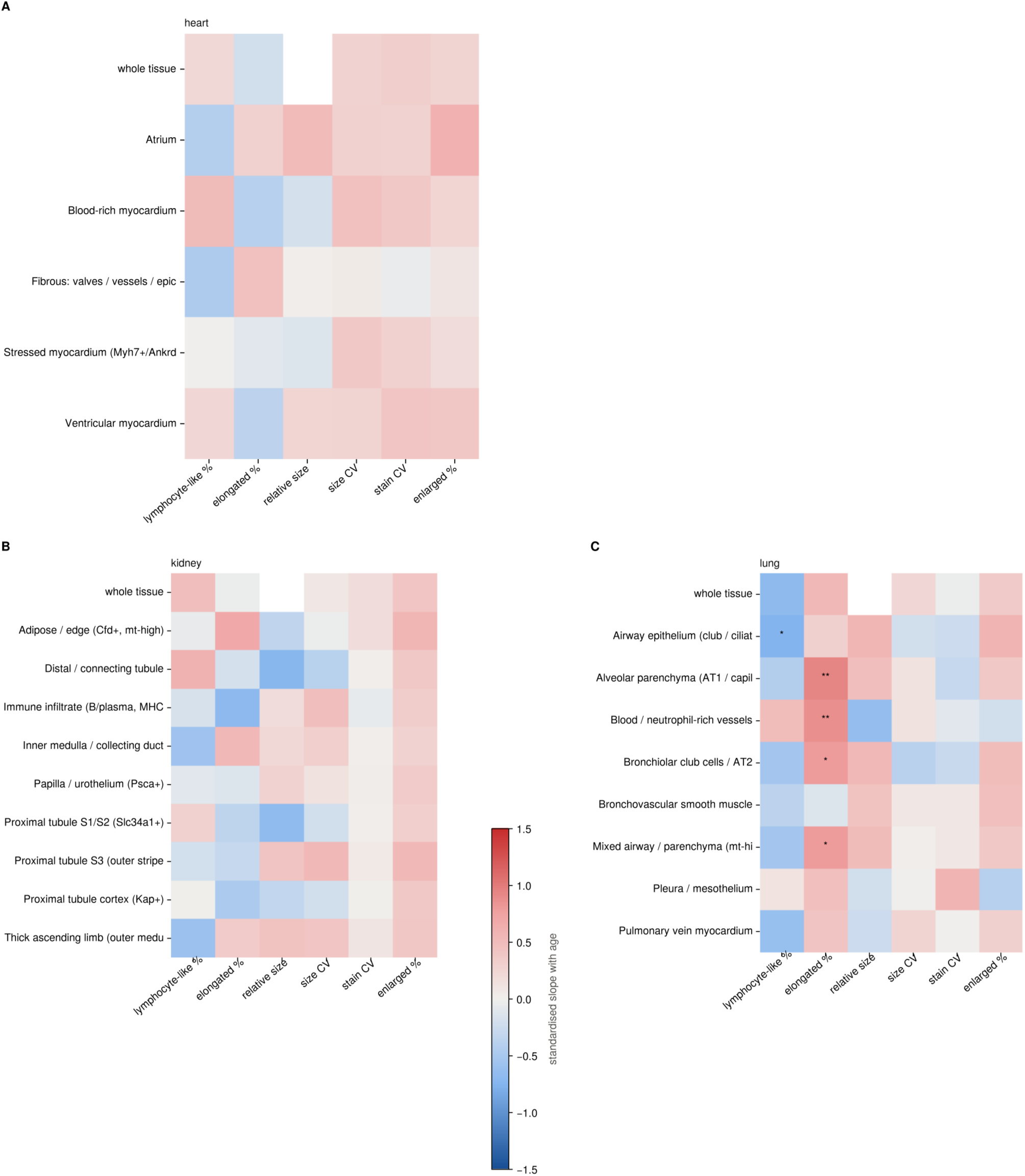
Nuclear morphology with age. Standardised age slope of lymphocyte-like and elongated nuclear fractions, relative nuclear size, coefficient of variation of size and stain and the fraction of enlarged nuclei, per mouse, whole tissue and per niche, with a focus-adjusted sensitivity model (pre-registered Addendum B; * q < 0.05): **A,** heart and **B,** kidney, without significant change. **C,** Lung: elongated nuclei become more frequent in four niches and lymphocyte-like nuclei fewer in airway epithelium (q < 0.05). Brain gives no estimate, and liver (whole tissue only) does not change (Supplementary Data 5, sheet image_morphology_age). Nuclei segmented in the full-resolution H&E images (pre-registered Addenda A and B).

**Supplementary Fig. 35.**
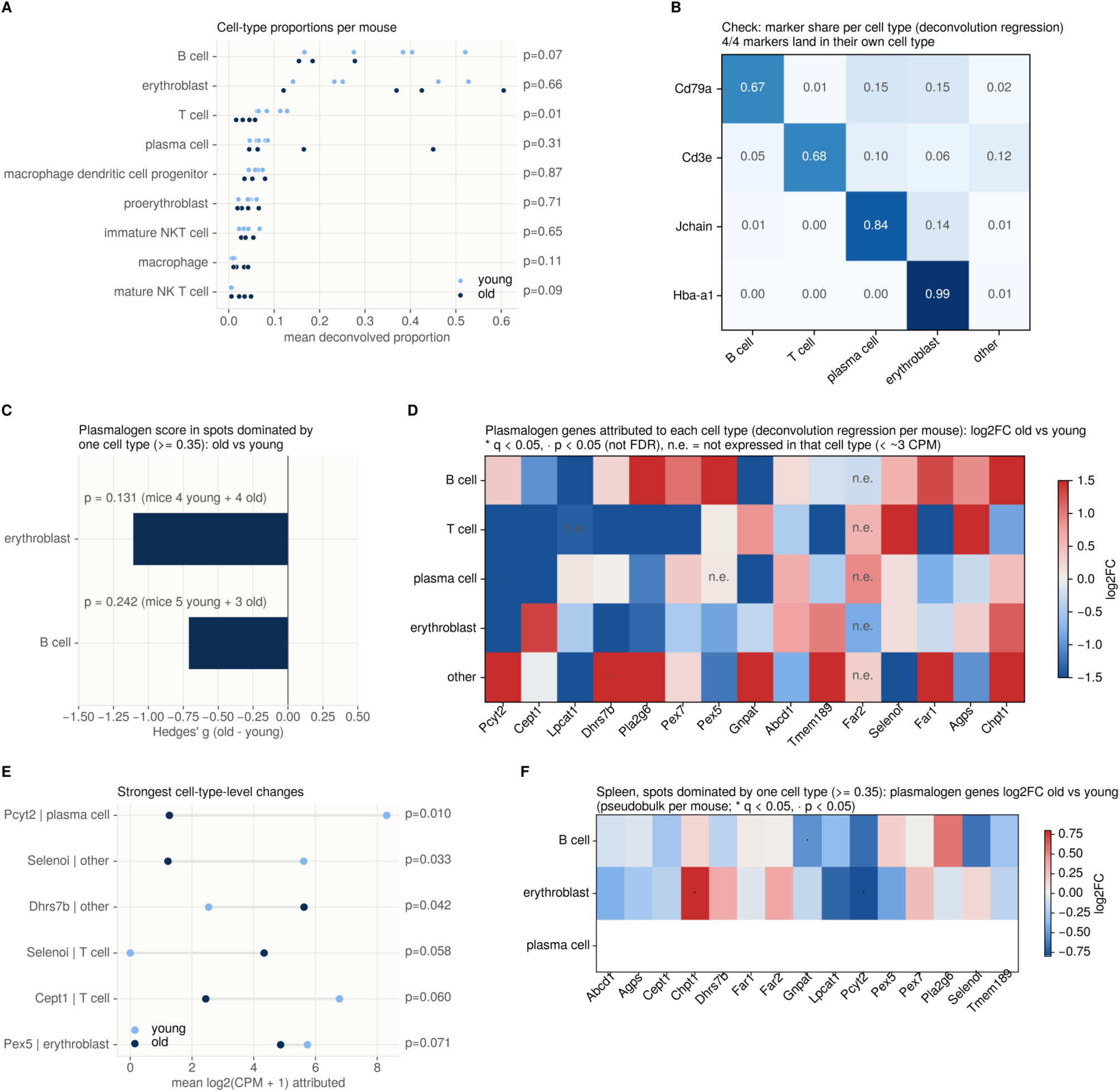
Spleen: cell-type composition, plasmalogen-synthesis score and pathway genes by dominant cell type. **A,** Deconvolved cell-type proportions per mouse, young versus old (Welch p): B cells 35% → 20% (p = 0.07), T cells 9.1% → 3.8% (p = 0.013, q = 0.16), plasma cells 6.8% → 18.1% (p = 0.31). **B,** Check of the attribution regression: share of each marker gene’s expression assigned to each cell type (*Cd79a*/B, *Cd3e*/T, *Jchain*/plasma, *Hba-a1*/erythroblast; 4 of 4 land in their own type). **C,** Plasmalogen-synthesis score in spots dominated (≥ 0.35) by erythroblasts (g = −1.1, p = 0.13) or B cells (g = −0.7, p = 0.24). **D,** Plasmalogen-pathway genes attributed to each cell type by non-negative regression of spot CPM on deconvolved proportions (per mouse): log2 fold change old versus young (n.e., not expressed). **E,** The strongest cell-type-level changes (*Pcyt2* in plasma cells, p = 0.010). The spleen decline is therefore partly compositional (fewer B and T cells). **F,** Plasmalogen-pathway genes in spots dominated (≥ 0.35) by one cell type: log2 fold change old versus young (pseudobulk per mouse; · p < 0.05, no gene q < 0.05). *Gnpat* falls in B-cell-dominated spots and *Pcyt2* (falling) and *Chpt1* (rising) change in erythroblast-dominated spots, each nominally; too few plasma-cell-dominated spots (empty row). GSE283283 Visium, young 3.9 versus old 25.8 months (3–5 mice per age and tissue); unit = mouse; Welch t-test and Hedges’ g, Benjamini–Hochberg (BH) within tissue. Secondary, descriptive analysis (not pre-registered).

**Supplementary Fig. 36.**
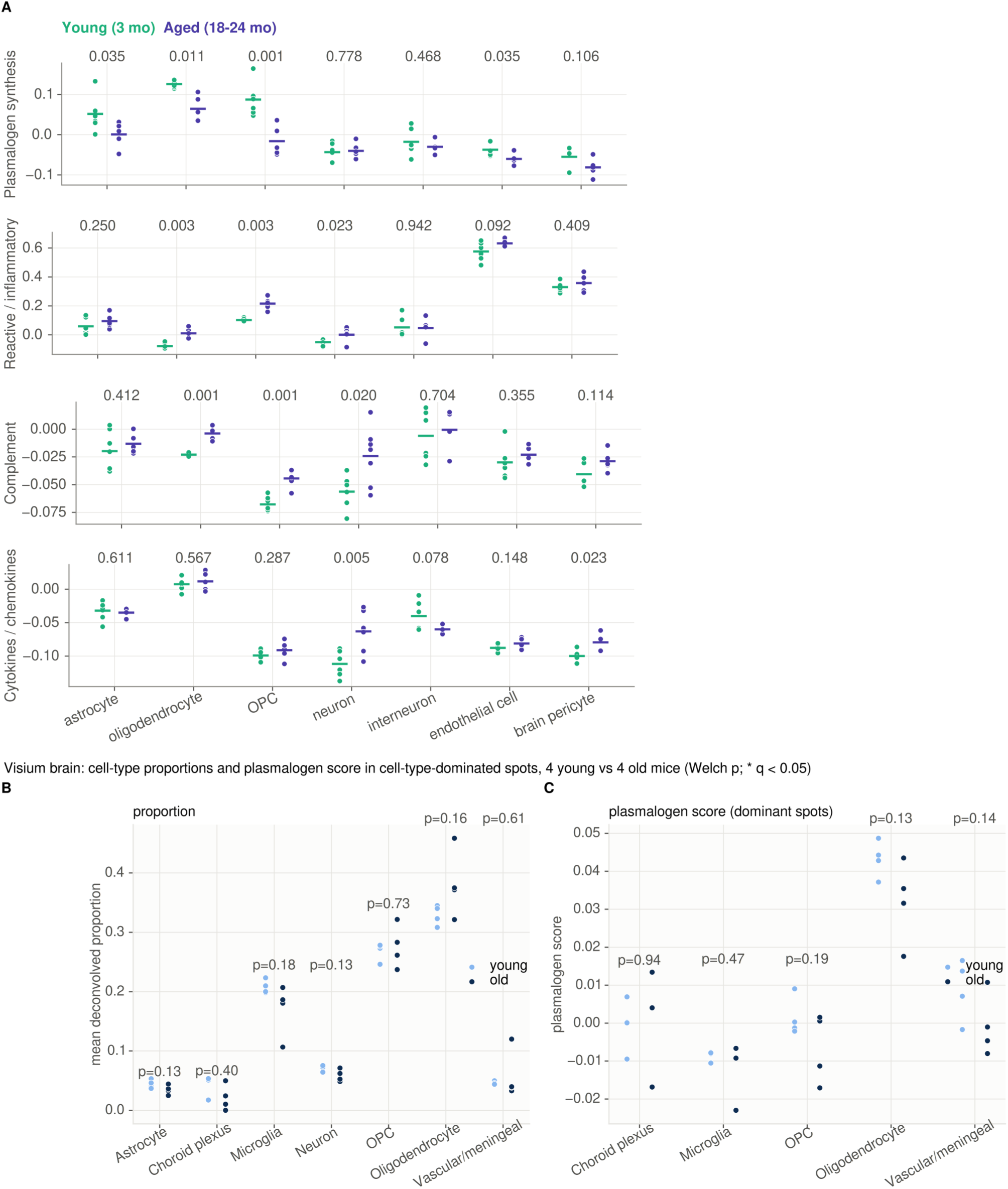
Non-myeloid brain cell types: plasmalogen-synthesis and reactive programmes; Visium brain: deconvolved cell types and their plasmalogen-synthesis score. **A,** Non-myeloid brain cell types: per-mouse plasmalogen-synthesis, reactive/inflammatory, complement and cy-tokine/chemokine scores in astrocytes, oligodendrocytes, OPCs, neurons, interneurons, endothelial cells and pericytes, young versus aged (printed values, Welch p). Plasmalogen synthesis falls in astrocytes (p = 0.035), oligodendrocytes (0.011), OPCs (0.001) and endothelial cells (0.035), not in neurons (0.78); oligodendrocytes and OPCs raise reactive and complement scores, neurons reactive, complement and cytokine scores. Tabula Muris Senis FACS brain (Smart-seq2; myeloid and non-myeloid sorts from the same mice), young 3 months versus aged 18–24 months; per-mouse mean scanpy score per cell type (≥ 5 cells); Welch t-test, Hedges’ g, BH. Secondary, descriptive analysis (not pre-registered). **B,C,** Visium brain: mean deconvolved cell-type proportion per mouse (**B**) and the plasmalogen-synthesis score in spots dominated (proportion ≥ 0.5) by one cell type (**C**), 4 young versus 4 old mice; no comparison reaches p < 0.1, reflecting the few dominated spots and mice. Visium brain, GSE283283 (4 young vs 4 old mice) and GSE233983 (10 vs 10 mice, both sexes). Secondary, descriptive analysis (not pre-registered).

**Supplementary Fig. 37.**
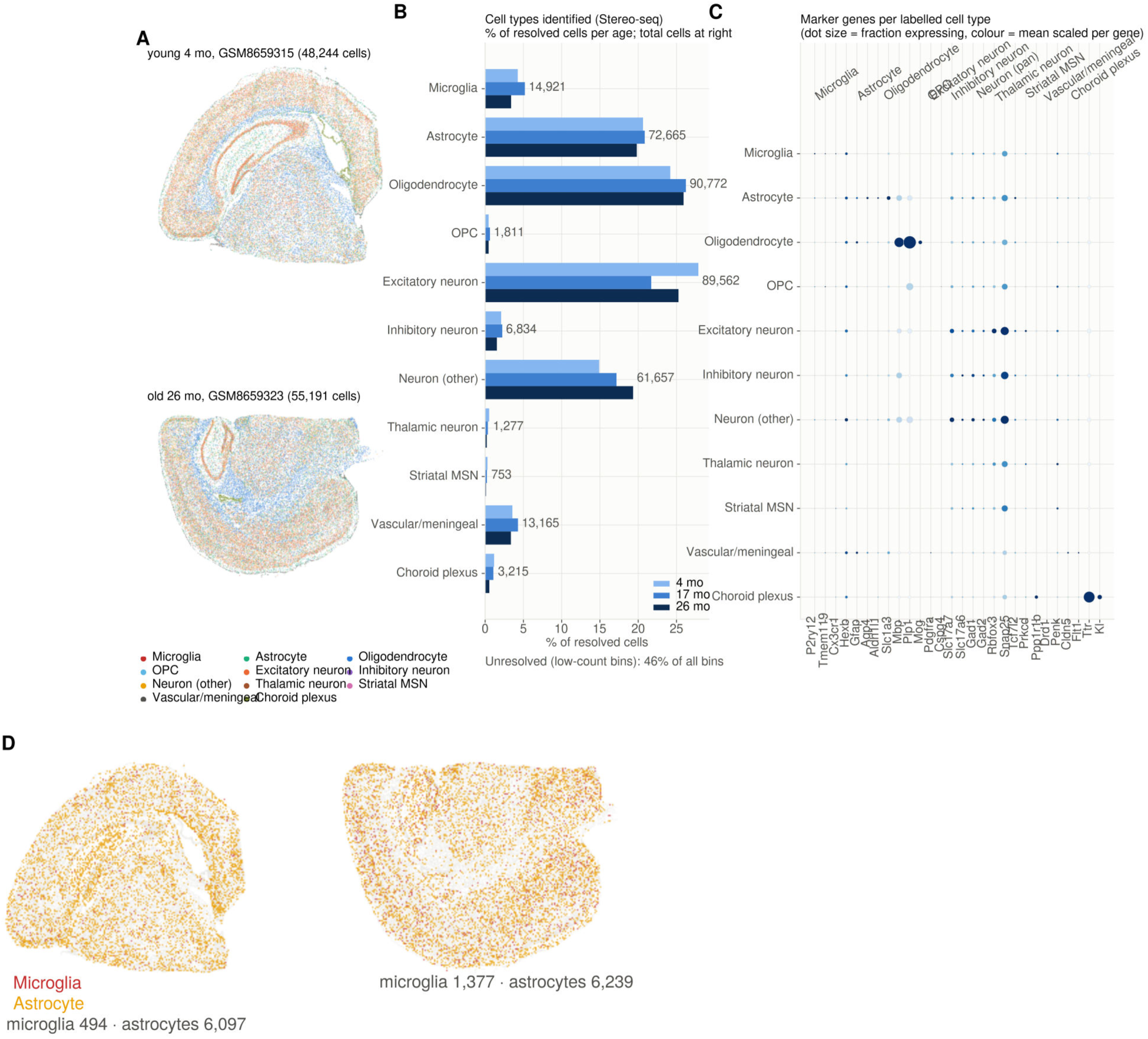
Stereo-seq brain cell types: atlas, marker genes and glial positions. Stereo-seq brain cell bins (GSE283283; one mouse per age, 4–5 sections each), labelled by a two-level marker rule (pre-registered amendment 2). **A,** Cell types on a young (4 months, 48,244 cells) and an old (26 months, 55,191 cells) section. **B,** Percentage of resolved cells per type and age (total cells at right; 46% of bins unresolved). **C,** Marker genes per labelled cell type (dot size, fraction expressing; colour, mean scaled per gene). **D,** Microglia (red) and astrocyte (orange) positions on the same young and old sections (microglia 494 versus 1,377, astrocytes 6,097 versus 6,239; one mouse per age, descriptive). The plasmalogen-synthesis and cytokine/chemokine landscapes of these sections are shown in Figs. 5D and 6F.

**Supplementary Fig. 38.**
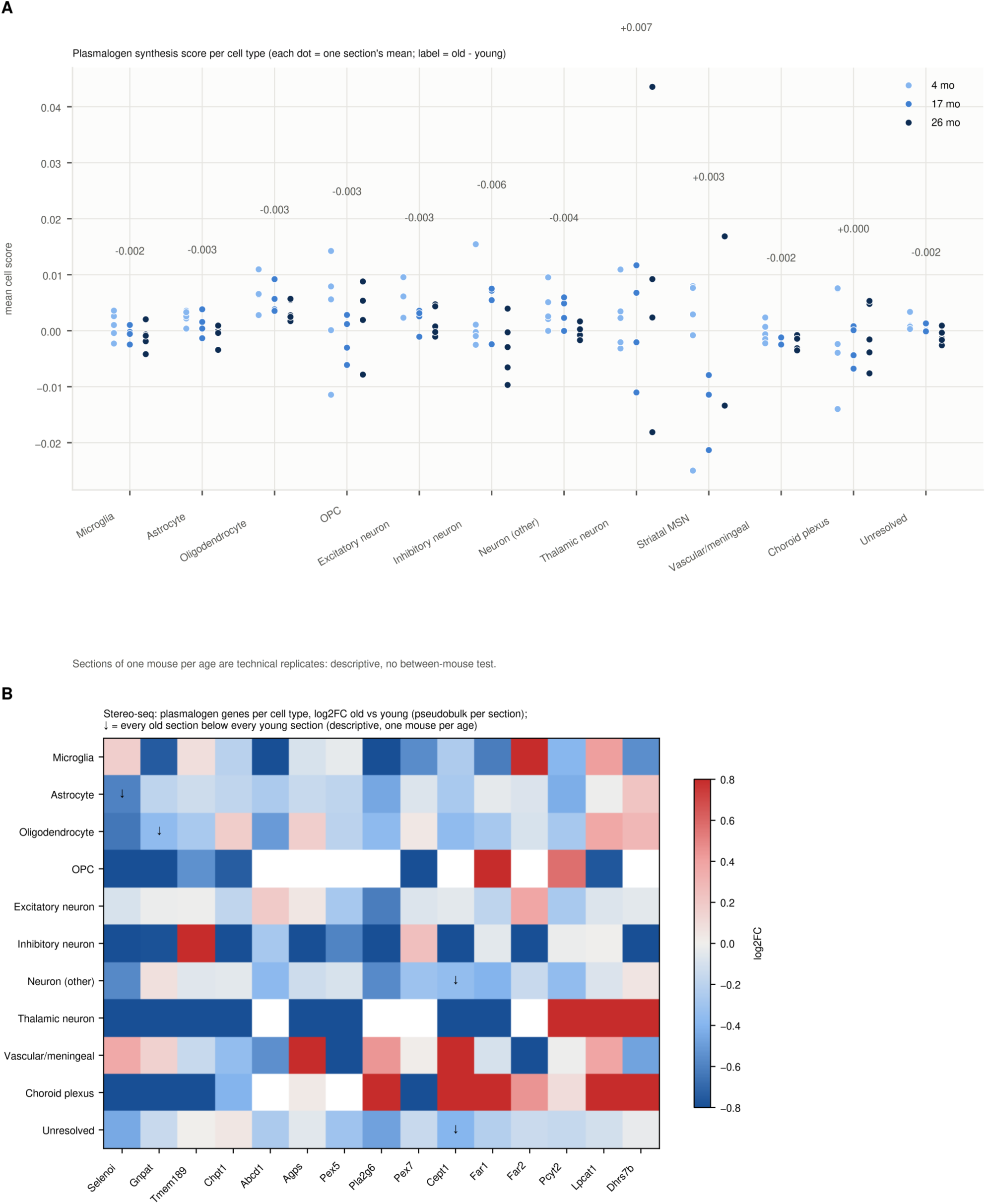
Stereo-seq: plasmalogen-synthesis score per cell type, by section; Stereo-seq: plasmalogen-pathway genes per cell type. **A,** Stereo-seq: plasmalogen-synthesis score per cell type, by section. Section-mean plasmalogen-synthesis score per cell type at 4 and 26 months; lower at 26 months in 8 of 11 resolved types (astrocytes g = −1.89, oligodendrocytes −1.13, microglia −0.79 between sections; thalamic neurons, striatal MSNs and choroid plexus higher). Sections of one mouse are technical replicates: descriptive, no p-values. **B,** Stereo-seq: plasmalogen-pathway genes per cell type. Old-minus-young difference of each plasmalogen-synthesis gene per Stereo-seq cell type (section means; descriptive).

**Supplementary Fig. 39.**
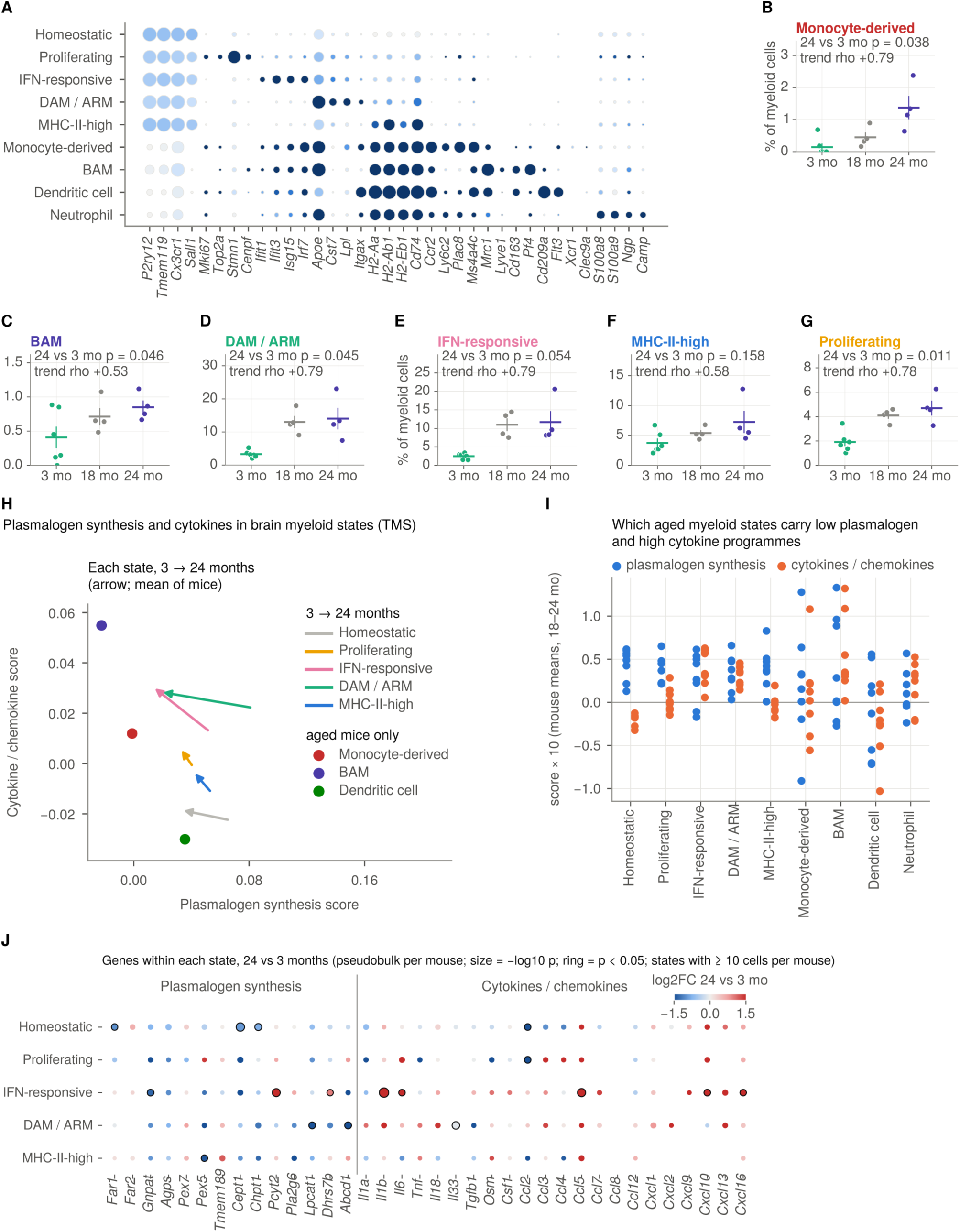
Brain myeloid states in ageing and the programmes within each state. TMS FACS brain myeloid cells (13,417 cells, 14 mice: 6 at 3, 4 at 18 and 4 at 24 months; UMAP of the states in Fig. 6E). States: homeostatic, proliferating, IFN-responsive, DAM/ARM and MHC-II-high microglia (marker scores > 2 SD above the 3-month mean; thresholds fixed before age was examined), monocyte-derived cells, border-associated macrophages (BAM), dendritic cells and neutrophils (cell numbers in Fig. 6E). **A,** Marker genes per state (dot size, fraction expressing; colour, mean expression). **B–G,** Percentage of myeloid cells per mouse at 3, 18 and 24 months (24 versus 3 months, Welch p; Spearman trend ρ): monocyte-derived (**B**; p = 0.038), BAM (**C**; p = 0.046), DAM/ARM (**D**; p = 0.045), IFN-responsive (**E**; p = 0.054), MHC-II-high (**F**; p = 0.16) and proliferating (**G**; p = 0.011). Homeostatic microglia fall from 87.7% to 58.7% (q = 0.03; main Fig. 6). Monocyte origin is inferred from Ccr2/Ly6c2, not lineage-traced. **H,** Each myeloid state drawn as an arrow from 3 to 24 months in plasmalogen-synthesis × cytokine/chemokine score space (mean of mice); every microglial state moves towards lower plasmalogen-synthesis and higher cytokine scores; BAM, monocyte-derived and dendritic cells are present in aged mice only. **I,** Per-mouse scores of each state in aged mice. **J,** Plasmalogen-synthesis and cytokine/chemokine genes within each state, 24 versus 3 months (pseudobulk per mouse; colour, log2 fold change; size, −log10 p; ring, p < 0.05; states with ≥ 10 cells per mouse): the plasmalogen-synthesis score falls within homeostatic (g = −2.14, q = 0.03) and DAM/ARM microglia (−2.07, q = 0.03), and IFN-responsive microglia raise *Il1b* (+2.37 log₂, q < 10⁻⁴) and *Ccl5* (+7.0, q = 0.025). The Hallmark inflammatory score falls within states, so the tissue-level inflammatory rise reflects the shift between states. Secondary, descriptive analysis.

**Supplementary Fig. 40.**
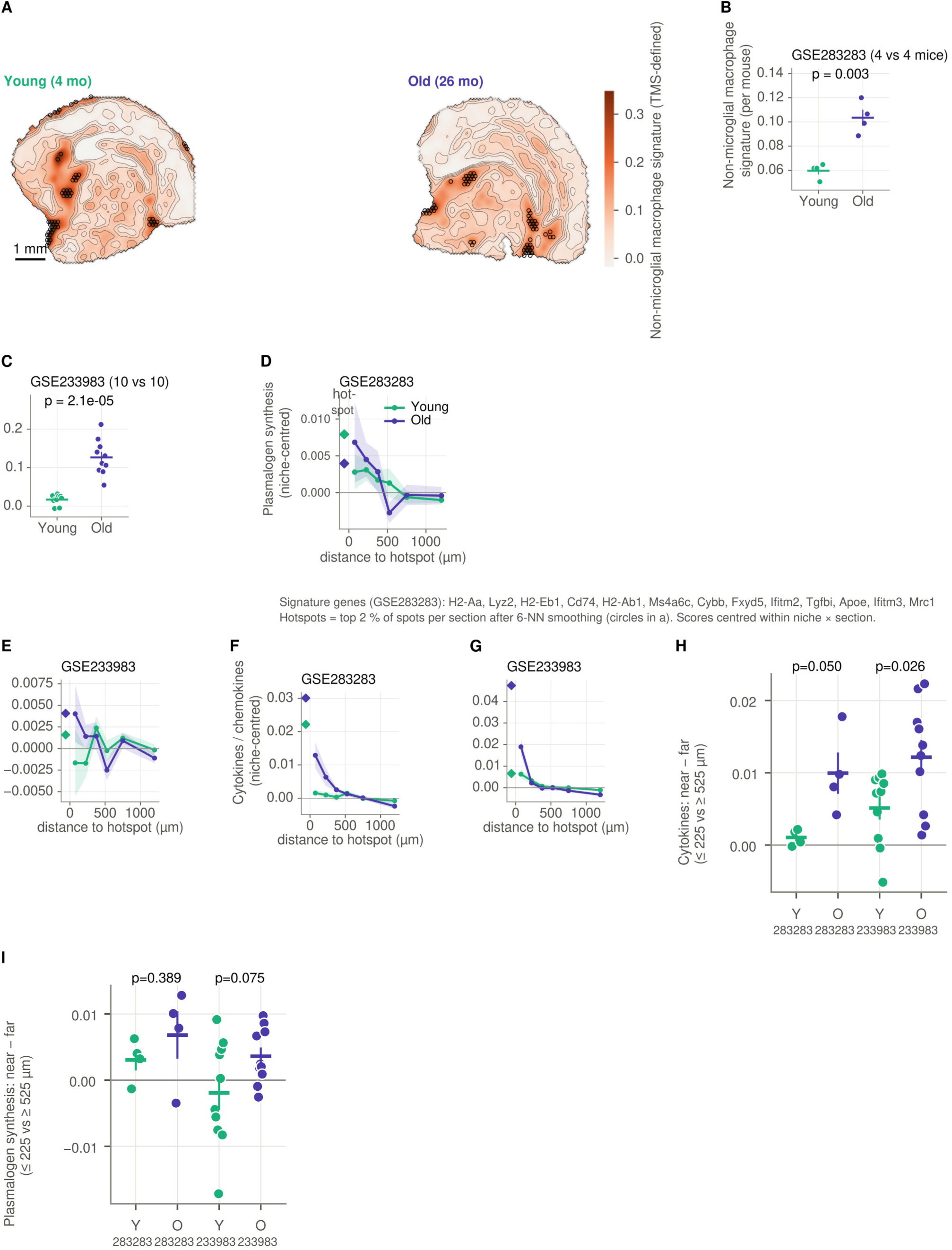
Monocyte-signature hotspots in brain tissue. TMS-derived non-microglial macrophage (monocyte) signature scored on Visium brain. **A,** Signature on a young and an old section; circles, hotspots (top 2% of spots per section after 6-nearest-neighbour smoothing). **B,C,** Signature per mouse, young versus old, in GSE283283 (**B**; p = 0.003) and GSE233983 (**C**; p = 2 × 10⁻⁵). **D,E,** Plasmalogen-synthesis score (niche-centred) with distance from hotspots in GSE283283 (**D**) and GSE233983 (**E**). **F,G,** The same for the cytokine/chemokine score. **H,** Cytokine score near (≤ 225 µm) minus far (≥ 525 µm) from hotspots per mouse: the gradient steepens with age (p = 0.050 and 0.026). **I,** The same for the plasmalogen-synthesis score: not lower near hotspots (p = 0.39 and 0.075). Visium brain, GSE283283 (4 young vs 4 old mice) and GSE233983 (10 vs 10 mice, both sexes). Secondary, descriptive analysis (not pre-registered).

**Supplementary Fig. 41.**
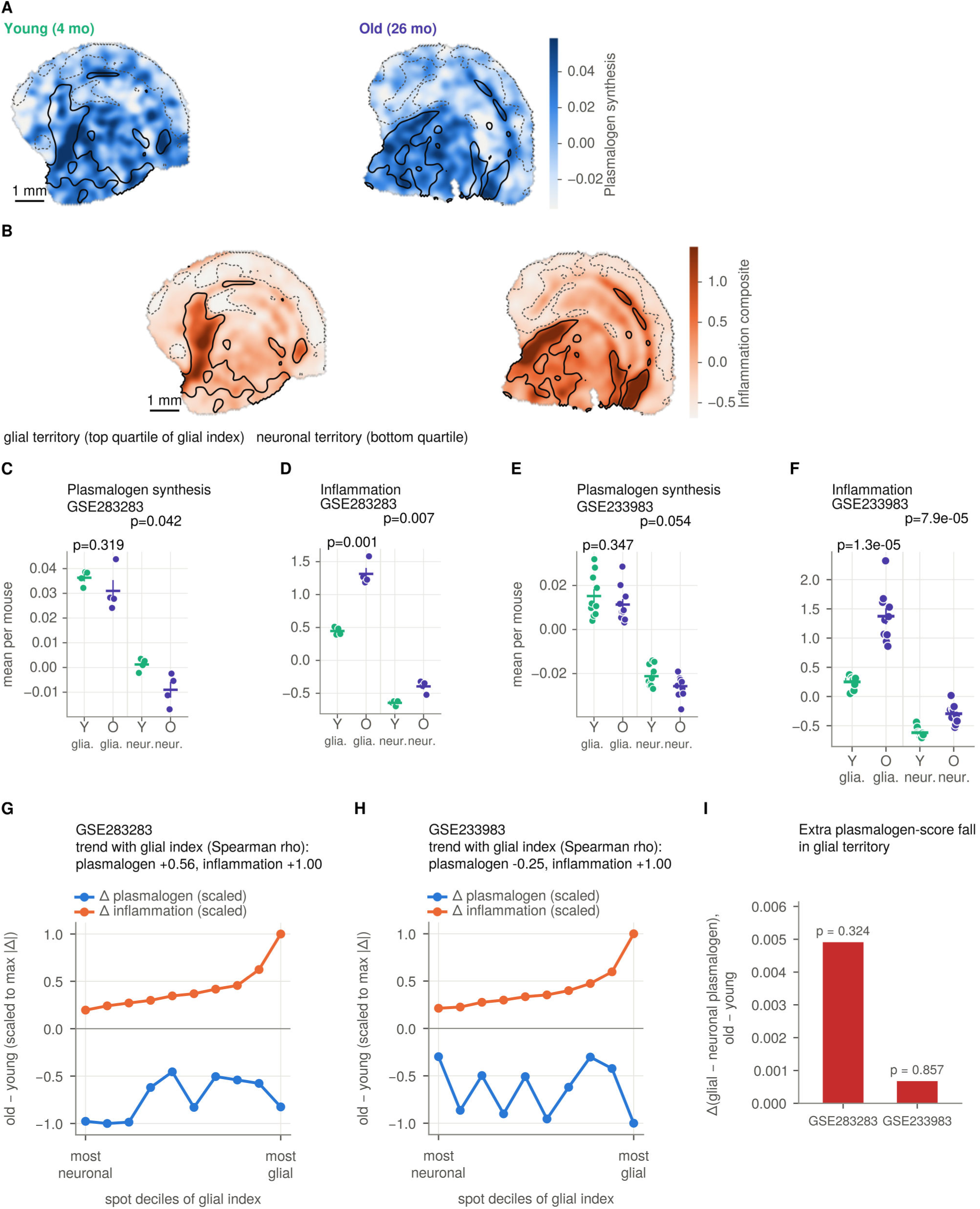
Glial and neuronal territories of ageing brain tissue; change of both scores along the neuronal → glial axis. **A–F,** Glial and neuronal territories of ageing brain tissue. Per section, spots in the top and bottom quartile of a glial index (TMS-derived oligodendrocyte, astrocyte and microglia signatures minus neuron signature) define glial and neuronal territories. Inflammation rises in both, most in glial territory (g = +5.6, p = 0.001; replication +3.4, p = 1 × 10⁻⁵); the plasmalogen-synthesis score falls in both, nominally only in neuronal territory (−1.83, p = 0.04; replication −0.88, p = 0.054). Spots (55 µm) mix cells, so neuronal territory also contains glia. Visium brain, GSE283283 (4 young vs 4 old mice) and GSE233983 (10 vs 10 mice, both sexes). Secondary, descriptive analysis (not pre-registered). **G–I,** Change of both scores along the neuronal → glial axis. Old-minus-young change of the plasmalogen-synthesis and inflammation scores in bins of the glial index, per mouse, in both Visium brain datasets. Visium brain, GSE283283 (4 young vs 4 old mice) and GSE233983 (10 vs 10 mice, both sexes). Secondary, descriptive analysis (not pre-registered).

**Supplementary Fig. 42.**
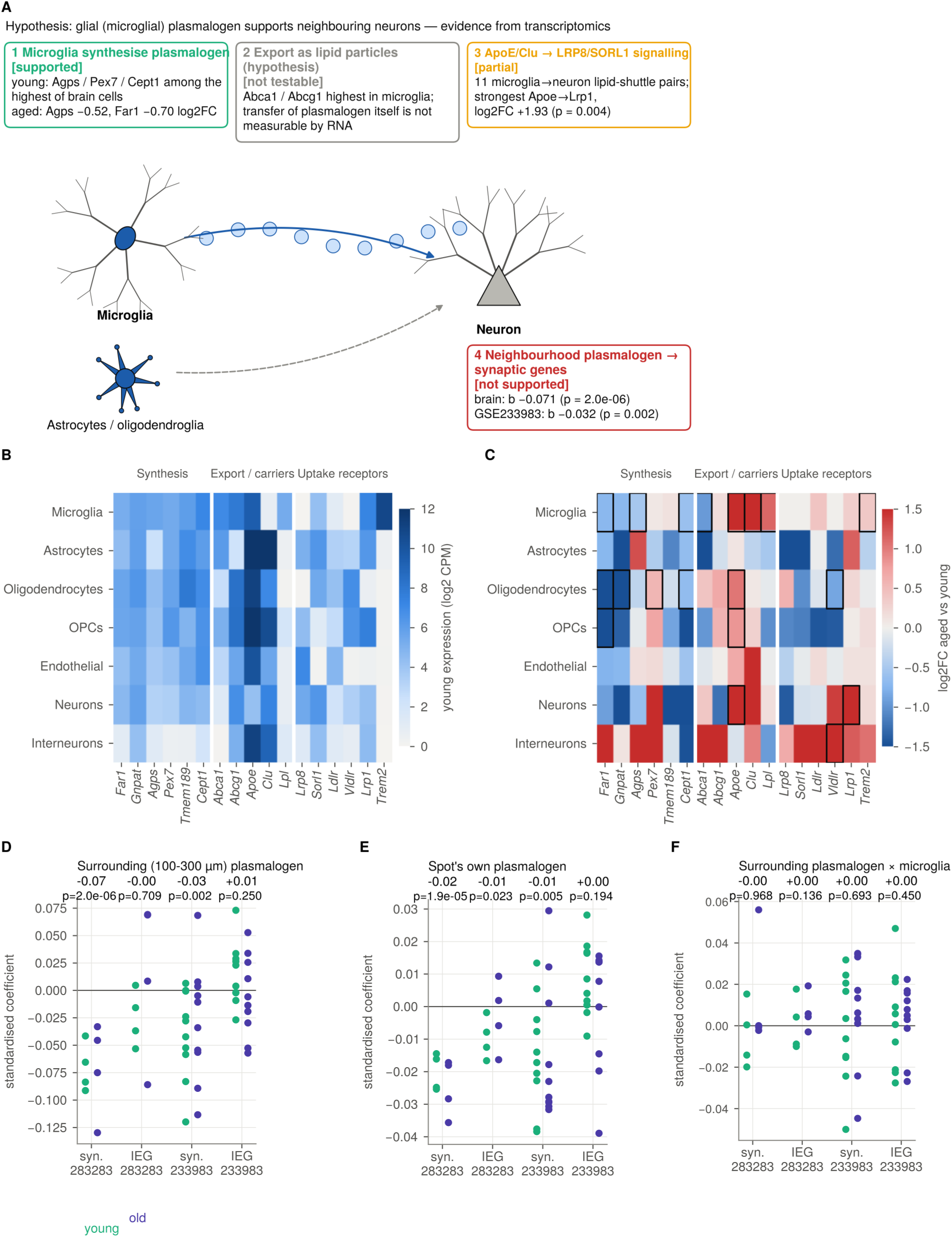
Glia-to-neuron lipid support: hypothesis, lipid-carrier and receptor genes, and neuronal-function genes. **A,** Glia-to-neuron lipid support: hypothesis and what transcripts can test. Schematic of the hypothesis that glia supply plasmalogen-containing lipids to neurons and that its loss impairs neuronal function, annotated with the transcript-level evidence of panels B–F. Lipid transfer itself is not measured. Exploratory. **B,C,** Expression of synthesis, lipid-carrier and uptake-receptor genes by brain cell type. Young expression and age change of plasmalogen-synthesis, lipid-export/carrier (Abca1, Abcg1, Apoe, Clu, Lpl) and uptake-receptor (Lrp8, Sorl1, Ldlr, Vldlr, Lrp1) genes in TMS astrocytes, microglia, neurons and other types; black box, p < 0.05. Apoe rises in aged microglia (+2.2 log₂) and neurons (+5.7). Tabula Muris Senis FACS brain (Smart-seq2; myeloid and non-myeloid sorts from the same mice), young 3 months versus aged 18–24 months; per-mouse mean scanpy score per cell type (≥ 5 cells); Welch t-test, Hedges’ g, BH. Secondary, descriptive analysis (not pre-registered). **D–F,** Neuronal-function genes and the plasmalogen-synthesis score of the neighbourhood. Synaptic-transmission (syn.) and activity-gene (IEG) scores per spot regressed, per section, on the plasmalogen-synthesis score of the surrounding ring (100–300 µm), the spot’s own score and cell-type signatures; coefficients per mouse (green, young; purple, old), printed mean and p. **D,** Ring coefficient: negative for synaptic genes in both datasets (p = 2 × 10⁻⁶ and 0.002), not for IEGs. **E,** The spot’s own score. **F,** Ring × microglia interaction: none. No evidence that neuronal-function genes depend on neighbouring plasmalogen synthesis. Visium brain, GSE283283 (4 young vs 4 old mice) and GSE233983 (10 vs 10 mice, both sexes). Secondary, descriptive analysis (not pre-registered).

**Supplementary Fig. 43.**
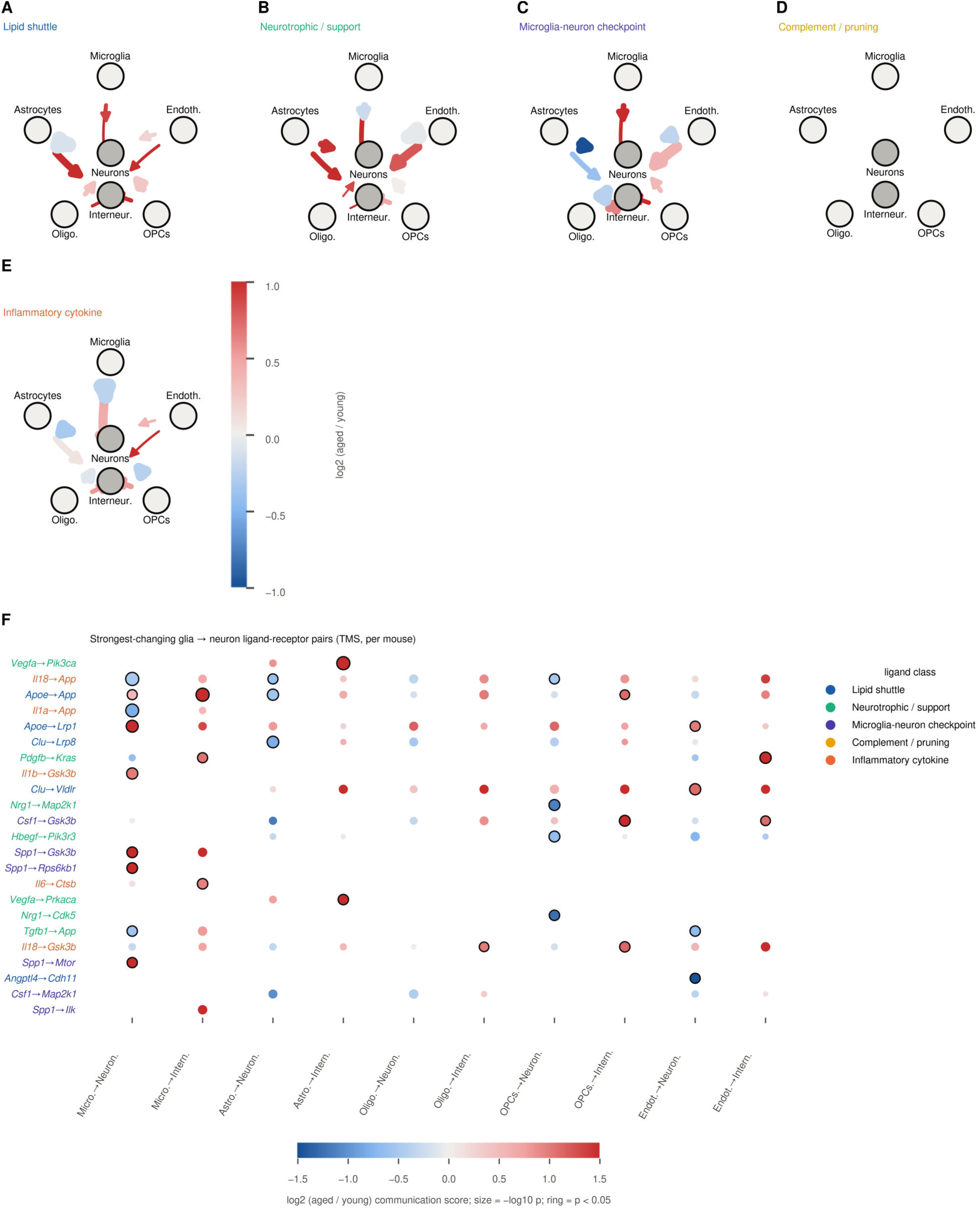
Glia-to-neuron ligand–receptor changes; Glia-to-neuron ligand–receptor pairs: dot plot. **A–E,** Glia-to-neuron ligand–receptor changes. OmniPath ligand–receptor pairs, sender glia to receiver neurons/interneurons (score = √(ligand × receptor), per mouse), coloured by category; edges, pairs with p < 0.05 for age (none for complement/pruning, **D**). No pair passes BH q < 0.1 within its sender–receiver combination. Tabula Muris Senis FACS brain (Smart-seq2; myeloid and non-myeloid sorts from the same mice), young 3 months versus aged 18–24 months; per-mouse mean scanpy score per cell type (≥ 5 cells); Welch t-test, Hedges’ g, BH. Secondary, descriptive analysis (not pre-registered). Exploratory. **F,** Glia-to-neuron ligand–receptor pairs: dot plot. Young and aged scores and log₂ fold change of the glia-to-neuron pairs in the lipid-shuttle, neurotrophic, checkpoint and inflammatory categories (e.g. microglia Apoe → neuronal Lrp1 +1.93 log₂, p = 0.004, q = 0.15). Tabula Muris Senis FACS brain (Smart-seq2; myeloid and non-myeloid sorts from the same mice), young 3 months versus aged 18–24 months; per-mouse mean scanpy score per cell type (≥ 5 cells); Welch t-test, Hedges’ g, BH. Secondary, descriptive analysis (not pre-registered). Exploratory.

**Supplementary Fig. 44.**
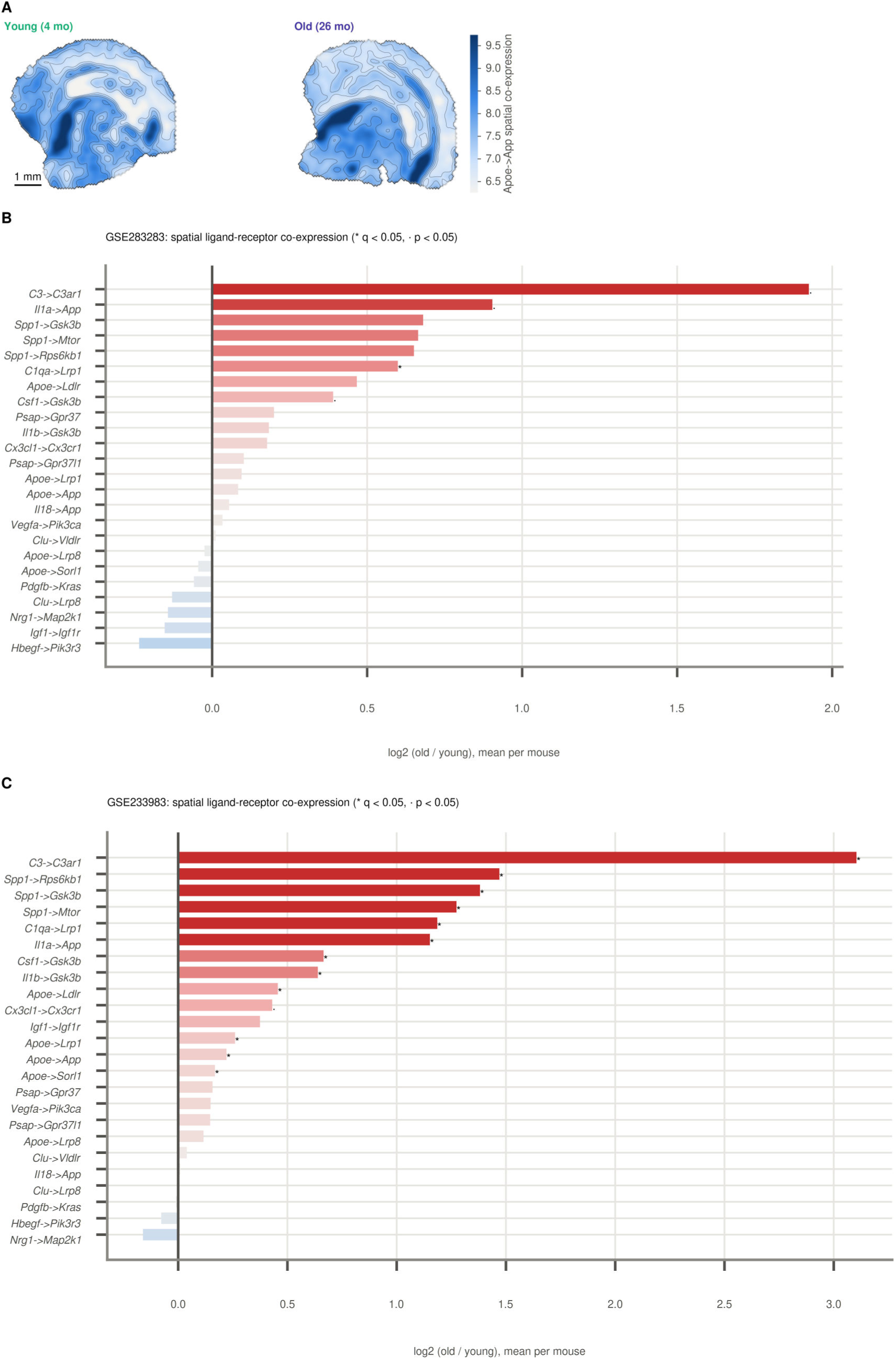
Spatial ligand–receptor co-expression. Per spot, ligand(spot) × mean recep-tor(spot + 6 neighbours). **A,** *Apoe*–*App* co-expression on a young and an old section. **B,C,** Old/young log2 ratio per pair (mean per mouse; * q < 0.05, · p < 0.05) in GSE283283 (**B**; *C1qa*–*Lrp1* q = 0.02) and GSE233983 (**C**; *C3*–*C3ar1*, *Spp1*–*Rps6kb1*/*Gsk3b*/*Mtor*, *C1qa*–*Lrp1*, *Il1a*–*App*, *Csf1*–*Gsk3b*, *Il1b*–*Gsk3b*, *Apoe*–*Ldlr*/*Lrp1*/*App*/*Sorl1* rise, q < 0.05). Visium brain, GSE283283 (4 young vs 4 old mice) and GSE233983 (10 vs 10 mice, both sexes). Secondary, descriptive analysis (not pre-registered). Exploratory.

## References

1. Braverman, N. E. & Moser, A. B. Functions of plasmalogen lipids in health and disease. Biochimica et Biophysica Acta (BBA) - Molecular Basis of Disease 1822, 1442–1452 (2012).

2. Hossain, M. S., Mawatari, S. & Fujino, T. Biological Functions of Plasmalogens. Advances in experimental medicine and biology 1299, 171–193 (2020).

3. Cipolla, C. M. & Lodhi, I. J. Peroxisomal Dysfunction in Age-Related Diseases. Trends in Endocrinology & Metabolism 28, 297–308 (2017).

4. López-Otín, C., Blasco, M. A., Partridge, L., Serrano, M. & Kroemer, G. The Hallmarks of Aging. Cell 153, 1194–1217 (2013).

5. López-Otín, C., Blasco, M. A., Partridge, L., Serrano, M. & Kroemer, G. Hallmarks of aging: An expanding universe. Cell 186, 243–278 (2023).

6. Chung, H. Y. et al. Molecular inflammation: Underpinnings of aging and age-related diseases. Ageing Research Reviews 8, 18–30 (2009).

7. Brodde, A. et al. Impaired neurotransmission in ether lipid-deficient nerve terminals. Human Molecular Genetics 21, 2713–2724 (2012).

8. Park, H. et al. Peroxisome-derived lipids regulate adipose thermogenesis by mediating cold-induced mitochondrial fission. Journal of Clinical Investigation 129, 694–711 (2019).

9. Kumar, A. et al. Age-dependent decline in plasmalogen biosynthesis impairs stress-induced mitochondrial fission in Drosophila. Experimental Cell Research 458, 114974 (2026).

10. Hossain, M. S. et al. Reduction of Ether-Type Glycerophospholipids, Plasmalogens, by NF-κB Signal Leading to Microglial Activation. The Journal of neuroscience: the ofiicial journal of the Society for Neuroscience 37, 4074–4092 (2017).

11. Youssef, M., Ibrahim, A., Akashi, K. & Hossain, M. S. PUFA-Plasmalogens Attenuate the LPS-Induced Nitric Oxide Production by Inhibiting the NF-kB, p38 MAPK and JNK Pathways in Microglial Cells. Neuroscience 397, 18–30 (2019).

12. Ali, F., Hossain, M. S., Sejimo, S. & Akashi, K. Plasmalogens Inhibit Endocytosis of Toll-like Receptor 4 to Attenuate the Inflammatory Signal in Microglial Cells. Molecular neurobiology 56, 3404–3419 (2019).

13. Hossain, M. S., Mawatari, S. & Fujino, T. Plasmalogen-Mediated Activation of GPCR21 Regulates Cy-tolytic Activity of NK Cells against the Target Cells. Journal of immunology (Baltimore, Md.: 1950) 209, 310–325 (2022).

14. Maeba, R. et al. Plasmalogens in human serum positively correlate with high-density lipoprotein and decrease with aging. Journal of atherosclerosis and thrombosis 14, 12–8 (2007).

15. Bozelli, J. C. & Epand, R. M. Plasmalogen Replacement Therapy. Membranes 11, 838 (2021).

16. Ginsberg, L., Rafique, S., Xuereb, J. H., Rapoport, S. I. & Gershfeld, N. L. Disease and anatomic specificity of ethanolamine plasmalogen deficiency in Alzheimer’s disease brain. Brain Research 698, 223–226 (1995).

17. Han, X., Holtzman, D. M. & McKeel, D. W. Plasmalogen deficiency in early Alzheimer’s disease subjects and in animal models: molecular characterization using electrospray ionization mass spectrometry. Journal of Neurochemistry 77, 1168–1180 (2001).

18. Fabelo, N. et al. Severe Alterations in Lipid Composition of Frontal Cortex Lipid Rafts from Parkinson’s Disease and Incidental Parkinson’s Disease. Molecular Medicine 17, 1107–1118 (2011).

19. Gu, J. et al. Plasmalogens Eliminate Aging-Associated Synaptic Defects and Microglia-Mediated Neuroinflammation in Mice. Frontiers in Molecular Biosciences 9, 815320 (2022).

20. Papsdorf, K. & Brunet, A. Linking Lipid Metabolism to Chromatin Regulation in Aging. Trends in Cell Biology 29, 97–116 (2019).

21. Hahn, O. et al. Atlas of the aging mouse brain reveals white matter as vulnerable foci. Cell 186, 4117–4133.e22 (2023).

22. Wang, L. et al. Spatial transcriptomics of the aging mouse brain reveals origins of inflammation in the white matter. Nature Communications 16, 3231 (2025).

23. GTEx Consortium. The GTEx Consortium atlas of genetic regulatory effects across human tissues. Science 369, 1318–1330 (2020).

24. Li, G.-H. et al. A multi-omics molecular landscape of 30 tissues in aging female rhesus macaques. Nature Methods 22, 2658–2669 (2025).

25. Schaum, N. et al. Ageing hallmarks exhibit organ-specific temporal signatures. Nature 583, 596–602 (2020).

26. Honsho, M., Asaoku, S. & Fujiki, Y. Posttranslational Regulation of Fatty Acyl-CoA Reductase 1, Far1, Controls Ether Glycerophospholipid Synthesis. Journal of Biological Chemistry 285, 8537–8542 (2010).

27. Tabula Muris Consortium. A single-cell transcriptomic atlas characterizes ageing tissues in the mouse. Nature 583, 590–595 (2020).

28. André, A. et al. Cerebral plasmalogens and aldehydes in senescence-accelerated mice P8 and R1: A comparison between weaned, adult and aged mice. Brain Research 1085, 28–32 (2006).

29. Rodemer, C. et al. Inactivation of ether lipid biosynthesis causes male infertility, defects in eye development and optic nerve hypoplasia in mice. Human Molecular Genetics 12, 1881–1895 (2003).

30. Teigler, A., Komljenovic, D., Draguhn, A., Gorgas, K. & Just, W. W. Defects in myelination, paranode organization and Purkinje cell innervation in the ether lipid-deficient mouse cerebellum. Human Molecular Genetics 18, 1897–1908 (2009).

31. Sol, J. et al. Ether lipids and sphingolipids drive sex-specific human aging dynamics. Redox Biology 85, 103779 (2025).

32. Hossain, M. S., Mawatari, S. & Fujino, T. Plasmalogens inhibit neuroinflammation and promote cognitive function. Brain research bulletin 192, 56–61 (2023).

33. Bottelbergs, A. et al. Peroxisome deficiency but not the defect in ether lipid synthesis causes activation of the innate immune system and axonal loss in the central nervous system. Journal of Neuroinflammation 9, 61 (2012).

34. Jin, K. et al. Brain-wide cell-type-specific transcriptomic signatures of healthy ageing in mice. Nature 638, 182–196 (2025).

35. Lee, D. et al. Single-cell atlas of transcriptomic vulnerability across multiple neurodegenerative and neuropsychiatric diseases. Preprint at 10.1101/2024.10.31.24316513 (2024).

36. Yazar, S. et al. Single-cell eQTL mapping identifies cell type–specific genetic control of autoimmune disease. Science 376, eabf3041 (2022).

37. Yang, W. et al. Single-cell, multi-region profiling of the macaque brain across the lifespan. Preprint at 10.1101/2025.10.31.685880 (2025).

38. Ma, S. et al. Caloric Restriction Reprograms the Single-Cell Transcriptional Landscape of Rattus Norvegicus Aging. Cell 180, 984–1001.e22 (2020).

39. Gao, S. M. et al. Aging atlas reveals cell-type-specific effects of pro-longevity strategies. Nature Aging 4, 998–1013 (2024).

40. Roux, A. E. et al. Individual cell types in C. elegans age differently and activate distinct cell-protective responses. Cell Reports 42, 112902 (2023).

41. Lu, T.-C. et al. Aging Fly Cell Atlas identifies exhaustive aging features at cellular resolution. Science 380, eadg0934 (2023).

42. Davie, K. et al. A Single-Cell Transcriptome Atlas of the Aging Drosophila Brain. Cell 174, 982–998.e20 (2018).

43. Lyu, P. et al. Comparative single-cell multiomic analysis reveals evolutionarily conserved and species-specific cellular mechanisms mediating natural retinal aging. Preprint at 10.1101/2025.09.09.673285 (2025).

44. Fleischer, J. G. et al. Predicting age from the transcriptome of human dermal fibroblasts. Genome Biology 19, 221 (2018).

45. Zhang, Y. et al. Purification and Characterization of Progenitor and Mature Human Astrocytes Reveals Transcriptional and Functional Differences with Mouse. Neuron 89, 37–53 (2016).

46. Galatro, T. F. et al. Transcriptomic analysis of purified human cortical microglia reveals age-associated changes. Nature Neuroscience 20, 1162–1171 (2017).

47. BrainSeq Consortium. Developmental and genetic regulation of the human cortex transcriptome illuminate schizophrenia pathogenesis. Nature Neuroscience 21, 1117–1125 (2018).

48. Li, M. et al. Integrative functional genomic analysis of human brain development and neuropsychiatric risks. Science 362, eaat7615 (2018).

49. Gatto, N. et al. Directly converted astrocytes retain the ageing features of the donor fibroblasts and elucidate the astrocytic contribution to human CNS health and disease. Aging Cell 20, e13281 (2020).

50. Mertens, J. et al. Directly Reprogrammed Human Neurons Retain Aging-Associated Transcriptomic Signatures and Reveal Age-Related Nucleocytoplasmic Defects. Cell Stem Cell 17, 705–718 (2015).

51. Wu, C. et al. Spatially resolved transcriptome of the aging mouse brain. Aging Cell 23, e14109 (2024).

52. Ding, J. et al. A metabolome atlas of the aging mouse brain. Nature Communications 12, 6021 (2021).

53. Milacic, M. et al. The Reactome Pathway Knowledgebase 2024. Nucleic Acids Research 52, D672–D678 (2023).

54. Liberzon, A. et al. The Molecular Signatures Database Hallmark Gene Set Collection. Cell Systems 1, 417–425 (2015).

55. Subramanian, A. et al. Gene set enrichment analysis: A knowledge-based approach for interpreting genome-wide expression profiles. Proceedings of the National Academy of Sciences 102, 15545–15550 (2005).

56. Love, M. I., Huber, W. & Anders, S. Moderated estimation of fold change and dispersion for RNA-seq data with DESeq2. Genome Biology 15, 550 (2014).

57. Korsunsky, I. et al. Fast, sensitive and accurate integration of single-cell data with Harmony. Nature Methods 16, 1289–1296 (2019).

58. Becht, E. et al. Dimensionality reduction for visualizing single-cell data using UMAP. Nature Biotechnology 37, 38–44 (2018).

59. Wolf, F. A., Angerer, P. & Theis, F. J. SCANPY: large-scale single-cell gene expression data analysis. Genome Biology 19, 15 (2018).

60. Cable, D. M. et al. Robust decomposition of cell type mixtures in spatial transcriptomics. Nature Biotechnology 40, 517–526 (2021).

61. Chen, A. et al. Spatiotemporal transcriptomic atlas of mouse organogenesis using DNA nanoball-patterned arrays. Cell 185, 1777–1792.e21 (2022).

62. Shamim, N., Fujiyama, A., Honsho, M., Mawatari, S. & Fujino, T. Ether-lipid deficiency remodels the testicular interstitium beyond spermatogenic arrest in Gnpat-knockout mice. bioRxiv [preprint; add DOI when posted] Preprint at (2026).

63. Benjamini, Y. & Hochberg, Y. Controlling the False Discovery Rate: A Practical and Powerful Approach to Multiple Testing. Journal of the Royal Statistical Society Series B: Statistical Methodology 57, 289–300 (1995).

64. Mogilenko, D. A., Shchukina, I. & Artyomov, M. N. Immune ageing at single-cell resolution. Nature Reviews Immunology 22, 484–498 (2021).

65. Leal, S. L. & Yassa, M. A. Neurocognitive Aging and the Hippocampus across Species. Trends in Neuro-sciences 38, 800–812 (2015).

66. Ledesma, M. D., Martin, M. G. & Dotti, C. G. Lipid changes in the aged brain: Effect on synaptic function and neuronal survival. Progress in Lipid Research 51, 23–35 (2012).

67. Moqri, M. et al. Biomarkers of aging for the identification and evaluation of longevity interventions. Cell 186, 3758–3775 (2023).

68. Moqri, M. et al. Validation of biomarkers of aging. Nature Medicine 30, 360–372 (2024).

